# Coevolutionary mining of prokaryotic non-coding elements with a genome language model

**DOI:** 10.64898/2026.09.22.753630

**Authors:** David B. Li, Garyk Brixi, Alexandra S. Kim, Mateus B. Fiamenghi, Claudia L. Driscoll, Simone A. Evans, Alex Gao, Natalia N. Ivanova, Nikos C. Kyrpides, Karl Deisseroth, Max E. Wilkinson, Michael A. Fischbach, Brian L. Hie

## Abstract

Microbial genomes encode non-coding RNAs (ncRNAs) and nucleic-acid-interacting proteins essential to biology and biotechnology. However, discovery of these systems remains protein-centric and based on similarity to known sequences. Here, we introduce Minerva, a framework for coevolutionary mining that uses genome language models to predict pairwise interactions directly from sequence. Minerva enables fast, alignment-free prediction of base pairing, monomeric protein contacts, and repetitive sequence motifs. We recover known systems such as CRISPR, detect open-reading-frame signatures at the DNA level without supervision, and predict hundreds of unannotated regions per bacterial genome containing multiple hairpins or repetitive motifs. In Pseudomonadota, we find that the widespread TwoAYGGAY structured RNA family carries large secondary-structure extensions and is often flanked by repetitive motifs. In prophages, we discover that Unknown Group 27 reverse transcriptase systems encode variable arrays of structurally conserved yet sequence-diverse ncRNAs that template complementary DNA hairpin products. Coevolutionary mining with Minerva enables efficient biological discovery across microbial genomes and metagenomes.

## Introduction

Microbial genomes encode diverse non-coding RNA and DNA elements that govern how organisms regulate, defend, and adapt. Despite their foundational importance, many non-coding sequences across prokaryotic genomes and metagenomes remain unannotated (*1–3*). This gap exists, in part, because standard discovery pipelines have prioritized protein-coding regions and homology to previously characterized sequences.

CRISPR offers a striking example: CRISPR loci were recognized purely from a pattern of DNA repeats years before their role in bacterial immunity was understood (*4–6*). Following the characterization of CRISPR as a programmable RNA-guided system, many distantly related nucleic-acid-guided systems have been discovered. Whereas the original CRISPR discovery arose from a property of the DNA sequence itself, these later systems have been prioritized by sequence or structural homology to conserved protein domains (*7–9*). The ability to read non-coding signals directly, without relying on a protein query, could yield many more nucleic-acid-interacting systems with distinct evolutionary histories and functions.

Directly predicting conserved interactions across genomes would enable the discovery of functional elements missed by homology-based approaches. Functionally coupled amino acids or nucleotides undergo correlated, compensatory mutations to maintain their interactions, leaving detectable coevolutionary imprints across sequences (*10–12*). Coevolutionary support across diverse genomes serves as a powerful filter because many energetically favorable interactions, such as incidental RNA base pairing, have no functional role. We use the term “coevolutionary mining” to describe the broader framework of predicting and leveraging evolutionary couplings for biological discovery.

Previous coevolutionary mining approaches have relied on multiple sequence alignments (MSAs) of homologs to detect covariation through statistical models. Advances in this MSA-centric approach, exemplified by the Infernal suite and CMfinder, have enabled bioinformatic ncRNA discovery and helped grow the Rfam database to more than four thousand families (*13–20*). In some cases, the covariation signal has even resolved RNA–protein interactions directly from paired MSAs (*20*). However, detecting covariation requires identifying and aligning a sufficient number of divergent homologs, which is particularly difficult for structured RNAs because secondary structure is often conserved beyond primary sequence (*21, 22*). Constructing suitable MSAs is also computationally intensive and often requires iterative refinement and manual curation (*17*).

Language models trained on biological sequences offer a promising alternative. These models learn statistical patterns that reflect biological structure and function from large unlabeled corpora of protein or nucleotide sequences (*23–30*). Masked language models (MLMs) are particularly well suited for capturing coevolutionary signals, since their training objective requires predicting hidden parts of a sequence from the surrounding context. For example, the identity of a masked nucleotide in an RNA stem is constrained by its base-pairing partner, so the objective rewards learning couplings maintained by compensatory substitutions.

Here, we present Minerva, a framework for query- and alignment-free coevolutionary mining from genome language models. Minerva efficiently predicts base pairing, protein contacts, and repetitive sequence motifs from genomic sequence inputs. We apply this tool to map unannotated structure in intergenic regions, uncover extensions to a known RNA family, and discover a phage-encoded ncRNA array that we experimentally show templates complementary DNA (cDNA) synthesis.

## Results

### Minerva accurately predicts pairwise coevolutionary maps from sequence

Given a genomic region, Minerva produces a two-dimensional coevolutionary map whose entries reflect how strongly, and in what way, each pair of positions interacts (**Fig. 1A**). This representation naturally captures arbitrary pairwise relationships, including pseudoknots, overlapping hairpins in a riboswitch, and repeat interactions. These maps support the discovery of new interactions, the annotation of known elements, and pattern-level search and retrieval of related sequences (**Fig. 1B**). Minerva predicts these maps by passing the input sequence through a genome language model and extracting coevolutionary interactions using two complementary methods. Both methods operate on standard parts of the transformer neural network architecture (*31, 32*), so the Minerva framework can be applied across different biological language models.

**Fig. 1.**
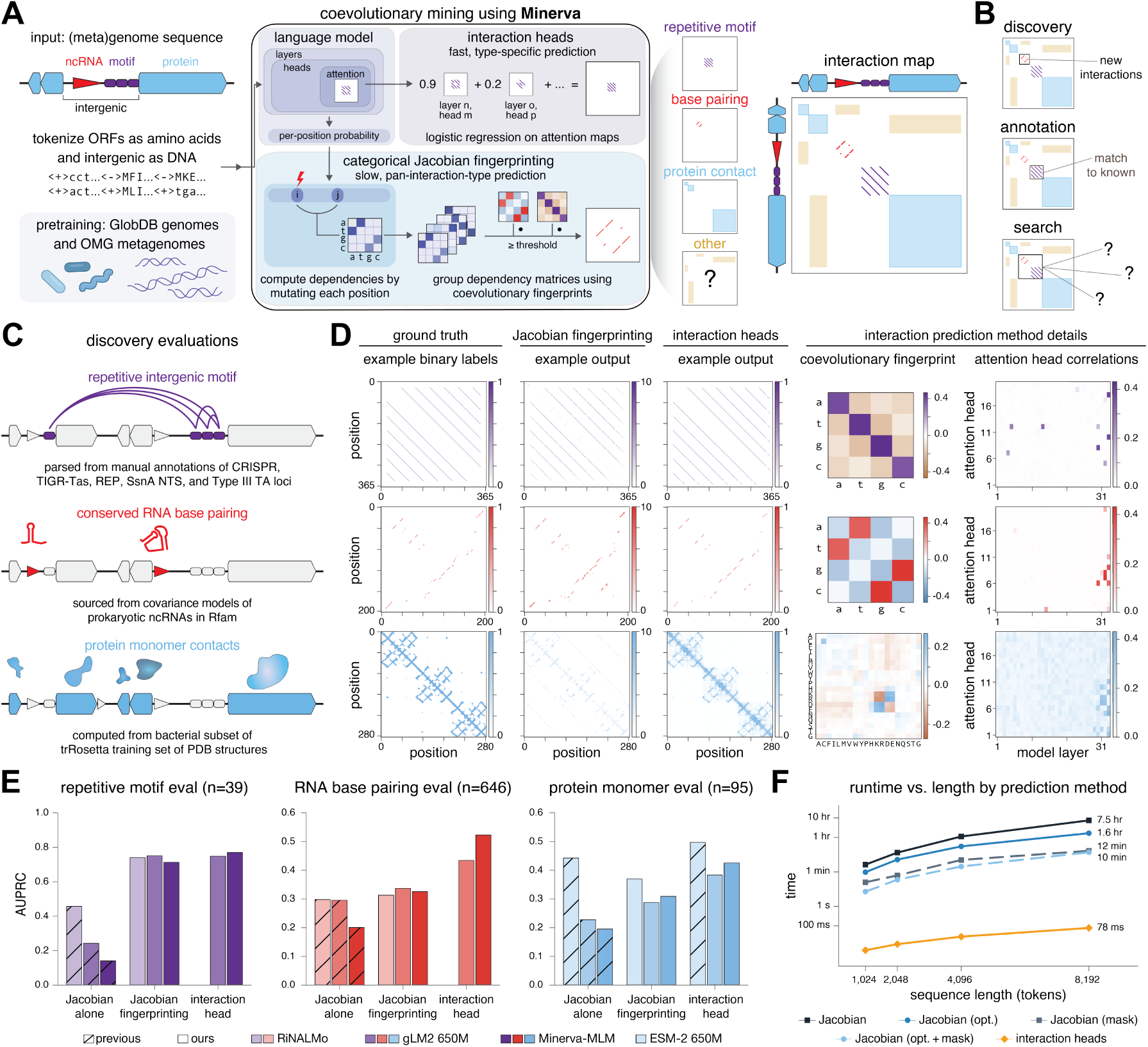
Minerva extracts pairwise coevolutionary maps from genome language models through interaction heads and categorical Jacobian fingerprinting. **(A**) Schematic of Minerva framework with input, output, interaction heads, and categorical Jacobian fingerprinting depicted. Pretraining and tokenization details depicted are for the Minerva-MLM genome language model; the framework generalizes to other models. **(B)** Intended use cases for the interaction map output of coevolutionary mining. **(C)** Depiction of evaluations for coevolutionary discovery covering conserved RNA base-pairing in bacterial ncRNAs, protein contacts in protein monomers, and repeat interactions in repetitive intergenic motifs. **(D)** Example ground truth and predictions for the discovery evaluations, as well as heatmaps of the reference Jacobian fingerprints used and of attention head correlations for the different interaction types. Examples shown are for a REP region from NC_010943, Rfam family RF04177, and chain A of PDB 2RJM. **(E)** Aggregate performance of categorical Jacobian, categorical Jacobian fingerprinting, and two-layer interaction heads across the repetitive-motif, conserved RNA base-pairing, and protein monomer-contact evaluations. Colors indicate the underlying language model; hatching distinguishes previously described methods from methods introduced here. **(F)** Run time as a function of input length for the categorical Jacobian and its optimized, mask-based, and combined variants, and for interaction heads. Annotations give the time per sequence at 8,192 tokens.

The quality of Minerva’s interaction predictions depends on the underlying pre-trained model. We therefore trained our own genome language model, Minerva-MLM. We initialized Minerva-MLM from gLM2 and retained gLM2’s mixed-modality tokenization, which uses amino acid tokens for protein sequences and DNA tokens for intergenic sequences (*25*). To improve modeling of long intergenic regions, which tend to be enriched for ncRNAs (**fig. S1A**), we continued pretraining for a trillion tokens from the OMG and GlobDB datasets (*25, 33*). Our primary checkpoint has a 4,096-token context window, corresponding to approximately 10 kb of DNA. We further extended the context window to 8192 tokens for the Minerva-MLM-8k model (**fig. S1**).

Common benchmarks for covariation prediction have primarily focused on monomeric protein contacts (*34*). To guide method development, we introduce two new evaluations spanning biologically distinct systems: RNA covariation from Rfam-derived base pairing and repetitive DNA motifs such as CRISPR arrays and repetitive extragenic palindromic (REP) elements, evaluated in their native genomic contexts (**fig. S2; Methods**). We use these alongside a protein monomer contact benchmark derived from the Protein Data Bank (**Fig. 1C**; *34*).

Our first interaction extraction method, categorical Jacobian fingerprinting (Jacobian fingerprinting for short), relies on *in silico* saturation mutagenesis. We mutate each position in the sequence to every possible nucleotide or amino acid and record the resulting changes in predicted probabilities at other positions. This yields a set of pairwise dependency matrices, referred to as the categorical Jacobian (*28*). Previous methods collapse each matrix into a scalar interaction score, discarding its internal structure (*28, 30, 35, 36*). Instead, we treat these distinct matrix patterns as “fingerprints” that can distinguish specific types of molecular interactions by comparison with reference patterns (**Fig. 1, A and D; figs. S3 and S4; Methods**). We derived these reference fingerprints by clustering dependency matrices from benchmark examples with known interaction labels. Repetitive-motif and base-pairing fingerprints reflect matching and reverse-complement rules, respectively. The protein-contact fingerprint resembles the Miyazawa–Jernigan statistical potential, with its dominant components corresponding to amino-acid physicochemical properties (**fig. S5;** *37–40*).

Although Jacobian fingerprinting flexibly represents diverse types of coevolutionary relationships, it requires a forward pass for each mutation, making genome-scale scanning impractical. We therefore asked whether the same interactions could be recovered from attention maps in a single forward pass of the model. Specific attention heads in Minerva-MLM were specialized for repetitive motifs, RNA base pairing, or protein covariation (**Fig. 1D; fig. S6)**. Lightweight logistic regression models, which we call interaction heads, further improved performance by combining multiple attention heads on a small set of supervised examples for each benchmark (**Methods;** *27*). Interaction heads using only the final two layers (instead of all layers) achieve high AUPRC across all benchmarks while reducing memory usage and improving throughput. We therefore use these two-layer interaction heads for our analyses and provide a last-six-layer variant with slightly higher accuracy (**fig. S7A)**.

Across all three benchmarks, both Jacobian fingerprinting and interaction heads outperform previous straight-to-scalar categorical Jacobian methods (*28, 35*) as measured by covariation prediction AUPRC (**Fig. 1E; figs. S7B and S8 to S11**). Interaction heads perform best overall, exceeding all other methods on more than 84% of individual Rfam families tested in the base-pairing benchmark (**fig. S11A; Supplementary Text**). The continued pretraining of Minerva-MLM relative to gLM2 further improved interaction head performance, albeit with reduced performance for the Jacobian-based methods, suggesting that coevolutionary information in model internal representations and output probabilities can diverge during training.

Performance relative to specialized language models depended on the interaction type. On the protein monomer benchmark, the protein language model ESM-2 outperforms both genome language models, consistent with its exclusive training on protein sequences (*23*). In contrast, Minerva-MLM interaction heads surpass the specialized RNA language model RiNALMo when predicting base pairing under conditions favorable to RiNALMo, using isolated RNA sequences (**Fig. 1E; figs. S7B and S11, A and B**). Neither the orientation nor RNA boundaries are available in the discovery setting, and placing the same RNA in its natural genomic context substantially reduces RiNALMo performance while slightly improving performance for Minerva-MLM (**fig. S11C**).

In addition to their higher accuracy, interaction heads are approximately five orders of magnitude faster than categorical Jacobian-based methods, requiring 79 ms per sequence compared with over 1.5 hours for an accelerated mask-token-based categorical Jacobian (**Methods**) and 6 hours for the original categorical Jacobian implementation with Minerva-MLM (**Fig. 1F**). This speedup makes genome-wide coevolutionary discovery practical. We thus adopt the Minerva-MLM interaction heads for primary genome scanning, and reserve Jacobian fingerprinting for more detailed locus-level interrogation, as the Jacobian can represent a broader unsupervised set of evolutionary relationships (*35*).

### Fine-tuning improves interaction predictions and codon periodicity emerges without supervision

Fine-tuning has previously improved the generative performance of DNA language models (*26, 41*) and interaction prediction by RNA language models, including through the categorical Jacobian (*30, 42*). We therefore tested whether family-specific fine-tuning could further improve Minerva predictions at individual loci (**Fig. 2A**). Because fine-tuning modifies the underlying model, it can improve predictions from both Minerva extraction methods.

**Fig. 2.**
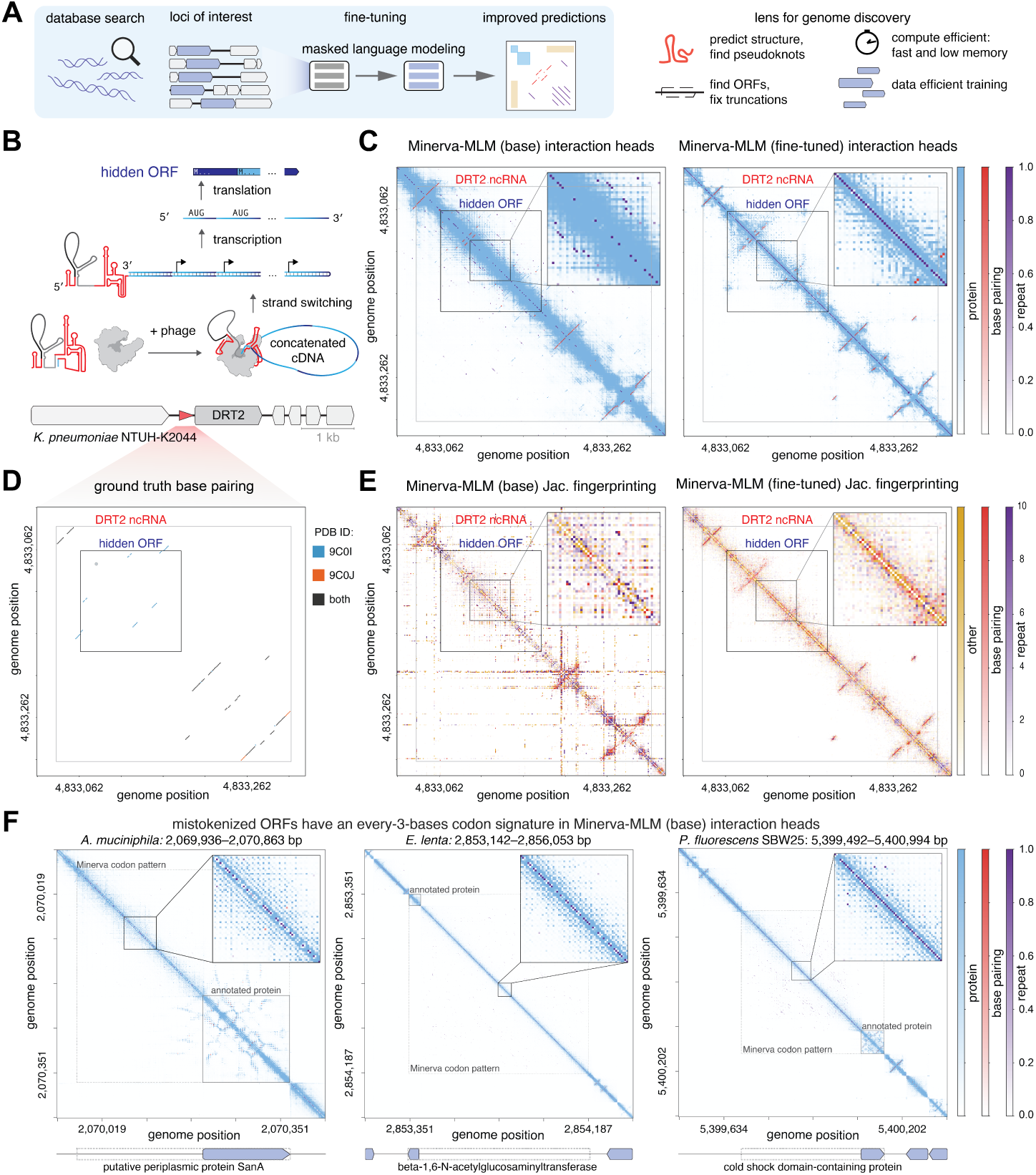
Fine-tuning improves interaction predictions, and codon periodicity emerges without supervision. **(A)** Schematic of family-specific fine-tuning and its use to improve interaction predictions at loci of interest. **(B)** Schematic of the DRT2 bacterial antiphage defense system. Upon phage infection, the DRT2 ncRNA is reverse transcribed to produce a concatemeric cDNA encoding a repetitive hidden ORF. **(C)** Minerva-MLM interaction-head predictions across the ncRNA and hidden ORF of the *Klebsiella pneumoniae* NTUH-K2044 DRT2 system before and after family-specific fine-tuning. Boxed regions are enlarged in the insets. **(D)** Reference base-pairing interactions derived from cryo-EM structures of the DRT2–ncRNA complex. Colors indicate interactions observed in PDB 9C0I, PDB 9C0J, or both structures. Hidden-ORF boundaries are also labeled. **(E)** Minerva-MLM categorical Jacobian fingerprinting predictions across the same DRT2 region before and after family-specific fine-tuning. Boxed regions are enlarged in the insets. **(F)** Minerva-MLM interaction-head predictions across three partially annotated protein-coding loci. An every-three-nucleotide interaction pattern extends beyond the annotated Bakta CDS boundaries, consistent with codon periodicity in the unannotated regions. The nearest upstream in-frame start codon is indicated for each locus.

We evaluated this approach on DRT2, a recently characterized bacterial antiphage defense system consisting of an Unknown Group 2 reverse transcriptase and an ncRNA that produces a repetitive open reading frame (ORF) upon phage infection (**Fig. 2B**; *43, 44*). Cryo-EM structures of both components, together with evidence that the ncRNA structure and hidden ORF are functionally conserved (*43, 44*), make DRT2 a well-characterized system for evaluating complex, system-specific predictions.

Interaction heads using the base Minerva-MLM model predict several hairpins present in the cryo-EM structure but miss others, including a pseudoknot involving the 3’ hairpin (**Fig. 2, C and D**). Fine-tuning on 13,034 DRT2 loci strengthened existing interactions and revealed additional base pairing, recovering the pseudoknot (**Fig. 2C; figs. S12 and S13**). The pseudoknot signal emerged after fine-tuning on as few as 200 loci, suggesting that relatively small sets of homologous sequences can support family-specific adaptation (**fig. S13**). Categorical Jacobian fingerprinting showed similar improvements (**Fig. 2E**).

The hidden ORF, which escaped conventional annotation and was only revealed through characterization of the DRT2 cDNA product (*43, 44*), was also detectable using the base Minerva-MLM model. The protein covariation interaction head and Jacobian fingerprinting both reveal a repeating three-nucleotide pattern across the ORF consistent with codon periodicity (**Fig. 2, C and E**). Unlike the RNA-structure signal, this pattern did not clearly improve with fine-tuning, suggesting that it is learned during pretraining on diverse sequences.

We observe the same codon-periodicity signal in protein-coding regions mistokenized as DNA due to mettannotator annotation errors (*45*). For several such loci, extending the annotated protein to the nearest in-frame start codon at the boundary of the periodic signal recovered full-length ORFs matching RefSeq proteins with 100% identity by BLASTp (**Fig. 2F**). More generally, these observations show that Minerva coevolutionary maps capture sequence relationships beyond the interaction classes they were explicitly trained to detect. Together, these capabilities suggest a unified discovery workflow: genome-scale scanning with base-model interaction heads to flag potentially novel interactions and unannotated ORFs, followed by targeted fine-tuning to resolve complex structures using both interaction heads and Jacobian fingerprinting.

### Genome-scale coevolutionary mining reveals intergenic structure

We applied Minerva genome-wide to search for intergenic structure beyond known annotations. Our dataset comprised 150 bacterial genomes selected from the hCom2 human gut microbiome community (*46*), selectagent and pathogen lists, and laboratory model organisms (**Fig. 3A; table S1**). We tiled each genome in overlapping chunks using the Minerva-MLM interaction heads and stored the resulting coevolutionary maps. The complete scan took 100 minutes on a single H100 GPU and required 18.3 GB of storage, producing 266–1,740 chunk maps per genome (**Methods**).

**Fig. 3.**
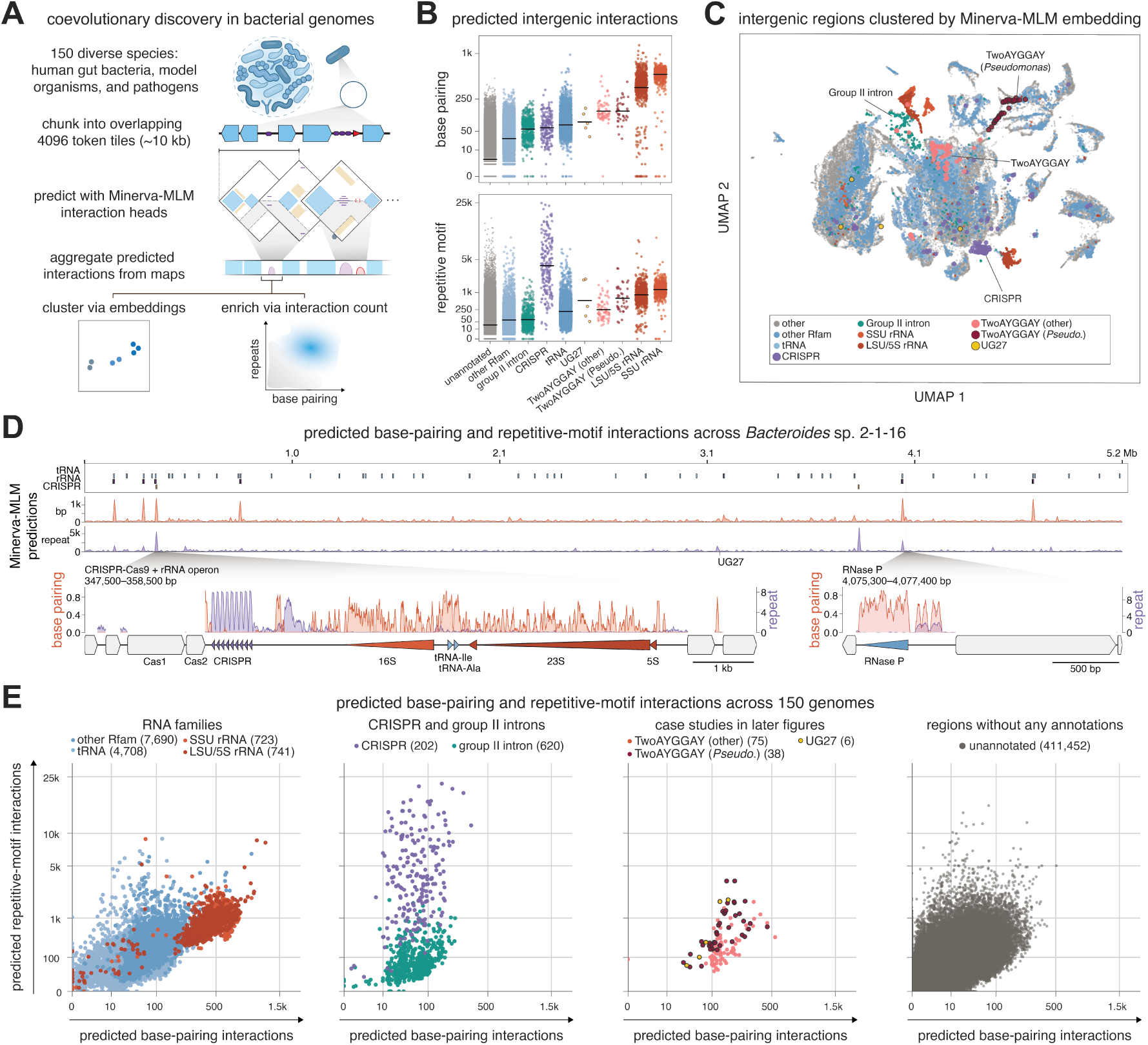
Genome-scale coevolutionary mining reveals structured intergenic loci across 150 bacterial genomes. **(A)** Minerva scanning workflow, applied to 150 bacterial genomes. Genomes are tiled using Minerva-MLM interaction heads to predict repeating and base-pairing nucleotides in intergenic sequence. Scores above threshold are then counted per intergenic region, enabling enrichment of structured intergenic regions. **(B)** Predicted base-pairing interactions (top) and repetitive-motif interactions (bottom) per intergenic region, grouped by mettannotator annotation class. Horizontal lines show the mean value per category. **(C)** UMAP of Minerva-MLM embeddings for the top 10% most structured intergenic regions across the 150 genomes as predicted by Minerva-MLM, colored by annotation class. Selected clusters labeled. **(D)** Per-position Minerva interaction scores as one-dimensional genome tracks for *Bacteroides* sp. 2-1-16, with annotated tRNA, rRNA, and CRISPR loci above. Insets, the CRISPR–Cas9 and rRNA operon locus (347,500–358,500 bp) and the RNase P locus (4,075,300–4,077,400 bp), showing base-pairing (red) and repeat (purple) signal relative to element boundaries. The UG27 locus is marked. **(E)** Predicted repetitive-motif interactions versus predicted base-pairing interactions for all intergenic regions across the 150 genomes, split by annotation class: RNA families, CRISPR and group II introns, the two case studies pursued below, and regions without annotation. Region counts in parentheses. Both axes cube-root scaled.

Whereas our earlier benchmarks evaluated predictions on individual annotated elements, we now asked whether aggregate Minerva signals could identify structured loci across whole genomes, using mettannotator annotations as reference (*45*). By counting the number of unique predicted repetitive-motif and base-pairing interactions across entire intergenic regions, we observed the expected enrichments such as CRISPR loci carrying the strongest repetitive-motif signal and rRNAs showing the highest base-pairing signal (**Fig. 3B**). tRNA-containing regions were enriched for both signals, consistent with their dense secondary structure and frequent clustering. Intergenic regions flanking proteins associated with structured nucleic acids, including transposases, were also enriched for predicted structure (**fig. S14**).

As a complementary representation, Uniform Manifold Approximation and Projection (UMAP) of Minerva-MLM embeddings from regions with high predicted covariation revealed groupings corresponding to several known element classes, including CRISPR loci, rRNAs, and group II introns (**Fig. 3C**). The embedding space also reflected sequence composition and phylogeny (**fig. S15**), indicating that purely embedding-based discovery of new families may require further post-training or decomposition (*29, 47, 48*). For genomescale visualization, summarizing the two-dimensional coevolutionary maps as one-dimensional tracks likewise highlighted known structured elements, including rRNAs and CRISPR arrays (**Fig. 3D; fig. S16**).

We then used predicted base pairing and repetitive interactions to survey unannotated intergenic regions (**Fig. 3E**). Many unannotated regions showed signals comparable to known structured RNAs and repeat arrays, including regions highly enriched for base pairing, repetitive motifs, or both. Within these genomes, Minerva predicted that 57,787 loci contain at least two neighboring hairpins with stems of five or more base pairs, with 40,175 (69.5%) outside of existing annotations. A more stringent threshold of four or more hairpins identified 9,044 loci, of which 41.1% were unannotated. This suggests that a large fraction of structured noncoding loci in bacterial genomes remains uncharacterized, including complex multi-hairpin elements. Two classes of loci were particularly notable: those containing matches to the TwoAYGGAY structured RNA family (*14, 19*) and those associated with Unknown Group 27 (UG27) reverse transcriptases (*49*), both of which we investigate below.

### Structure extension and repeat association in Pseudomonadota TwoAYGGAY RNAs

The canonical TwoAYGGAY structured RNA motif (Rfam RF01731) consists of two sequential hairpins emerging from a common basal stem, each with an AYGGAY-containing terminal loop (*19*). Intergenic regions containing TwoAYGGAY matches were strongly enriched for both predicted base pairing and repetitive motifs. In genomes other than *Pseudomonas fluorescens* SBW25, these loci showed 3.7-fold enrichment in repetitive-motif signal and 10.9-fold enrichment in base-pairing signal relative to all intergenic regions. In SBW25, the base-pairing signal was comparable (10.6-fold enrichment) and the repetitive-motif signal was substantially stronger (13.6-fold enrichment), prompting us to examine the organization of TwoAYGGAY loci in this genome in greater detail.

Inspection of TwoAYGGAY loci in SBW25 with the Minerva-MLM base-pairing interaction head revealed predicted interactions extending substantially beyond existing Rfam annotations (**Fig. 4A; fig. S17**). When oriented relative to the Rfam match, the predicted structure contains three additional hairpins, an elongated basal stem, and a 3’ pseudoknot. The repetitive-motif interaction head further predicts distinct flanking repetitive sequences, with shorter 5–6-nt GMTBT(T) repeats at the 5’ end and longer 100–200-nt repeats at the 3’ end (**Fig. 4A**).

**Fig. 4.**
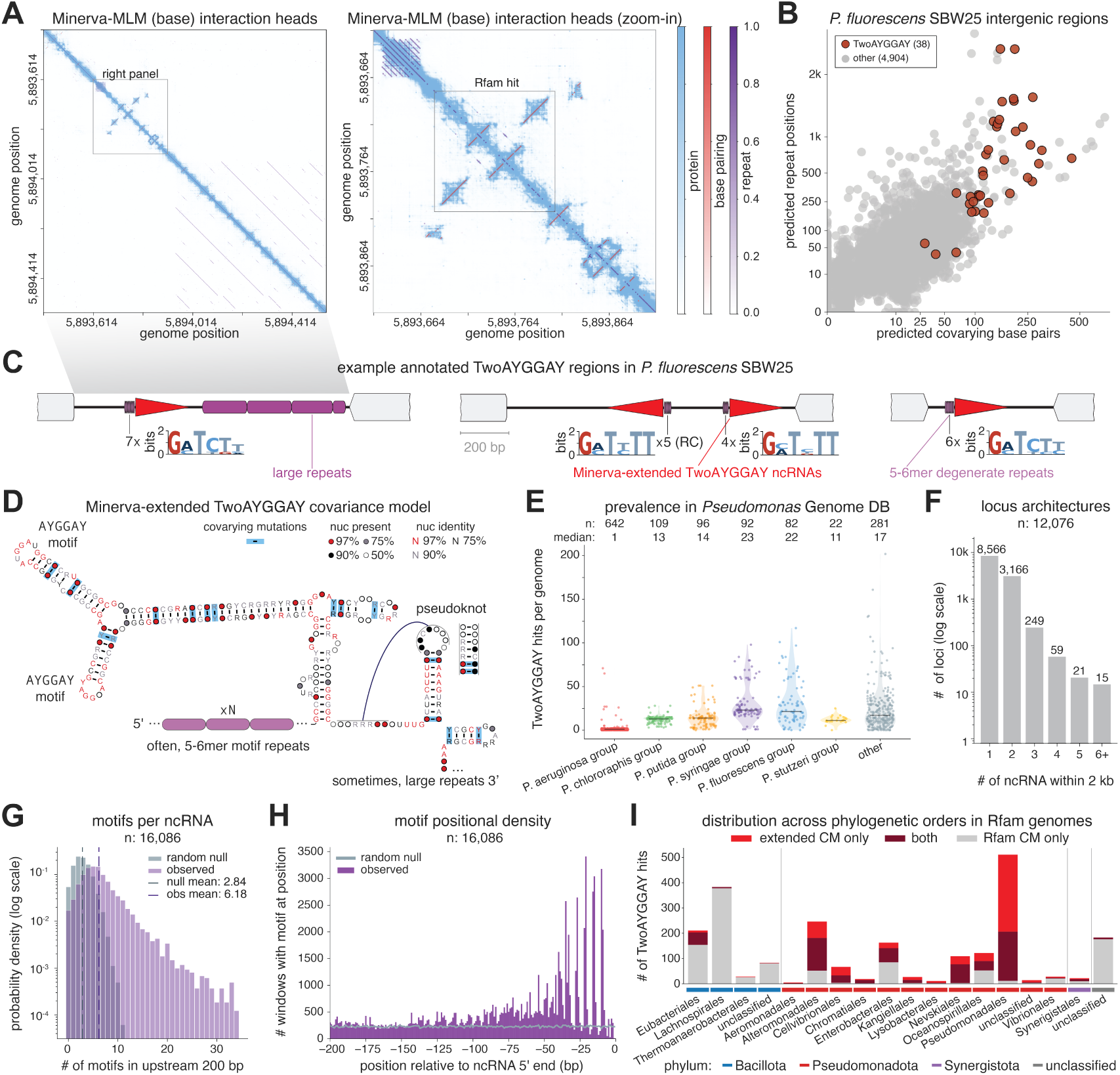
Minerva reveals extended structure and repeat association in Pseudomonadota TwoAYGGAY RNAs. **(A)** Minerva-MLM base-model interaction head predictions for a TwoAYGGAY-containing intergenic region in *Pseudomonas fluorescens* SBW25 (left), with the boxed region enlarged (right). Rfam RF01731 match boxed in the right panel. **(B)** Predicted repetitive-motif interactions versus base-pairing interactions across SBW25 intergenic regions. TwoAYGGAY-containing regions (n = 38) in red; all others (n = 4,904) in grey. Axes are cube-root scaled. **(C)** Example TwoAYGGAY loci in SBW25, showing the RNA match, upstream degenerate 5–6-mer repeat array with sequence logo and copy number, and downstream large repeats. RC, reverse-complement. **(D)** Minerva-extended TwoAYGGAY covariance model with associated repeat features drawn schematically: upstream 5–6-mer motif repeats (×N) and, at some loci, downstream large repeats. **(E)** TwoAYGGAY copy number across *Pseudomonas* species groups in the *Pseudomonas* Genome Database. Each point, one strain; horizontal bars, medians. Strain counts and median copy number listed above each group. **(F)** Number of extended TwoAYGGAY matches within 2 kb of one another, across all matches in the database (n = 12,076 loci). Counts above bars; y-axis log-scaled. **(G)** Distribution of 5’ motif (GATCT, GCTCT, GCTTT, GCTGT, GCCGT, or GCAGT) counts in the 200 bp upstream of extended TwoAYGGAY matches (n = 16,086), against randomly sampled regions from the same genomes as a null. Vertical lines, observed and null means. Y-axis log-scaled. **(H)** Positional density of 5’ motif matches within the 200 bp upstream of extended TwoAYGGAY matches, plotted relative to the TwoAYGGAY match 5’ end, with the same null as **(G)**. **(I)** TwoAYGGAY hits per phylogenetic order in Rfam genomes, classified by whether they are recovered by the Minerva-extended covariance model only, the Rfam RF01731 covariance model only, or both. Minerva-extended covariance model matches required ≥80% coverage. Phylum indicated below.

Across the SBW25 genome, TwoAYGGAY-containing intergenic regions represent many of the loci with the strongest predicted base-pairing and repetitive-motif signals (**Fig. 4B**). The genome contains 48 TwoAYGGAY matches distributed across 38 intergenic regions; seven matches are truncated (**Fig. 4C; figs. S18 and S19**). Alignment of the 41 non-truncated RNAs is consistent with the Minerva-predicted extended structure and reveals covariation among variable positions, although most positions are highly conserved (**fig. S18B**). The longer 3’ repeat signal also overlaps a previously described organization of repetitive extragenic palindromic (REP) elements (*50*). All 11 tandem group I/III REP arrays previously identified in SBW25 occur at a fixed position downstream of a TwoAYGGAY RNA match and contribute to the Minerva-predicted 3’-associated repeat signal (**Fig. 4A; fig. S19**).

To determine how broadly this extended RNA architecture is distributed across *Pseudomonas*, we fine-tuned Minerva-MLM on TwoAYGGAY matches identified by the original covariance model in the *Pseudomonas* Genome Database. Fine-tuning strengthened the extended stem prediction, although it modestly reduced the repetitive-motif signal (**fig. S20**). We then incorporated these predictions into an iteratively refined covariance model and scanned all complete genomes in the database (**Fig. 4D; Methods**). This scan identified 16,086 matches across 1,148 of 1,324 strains, spanning most *Pseudomonas* species groups but largely absent from the *P. aeruginosa* group (**Fig. 4E**). Copy number varied substantially across genomes. Median copy number ranged from 11 to 23 among non-*P. aeruginosa* groups, with as many as 202 matches in *Pseudomonas* sp. FDA-ARGOS 380. Large differences were also evident among strains of the same species, in some cases spanning tens to hundreds of copies per genome (**Fig. 4E; fig. S21**).

At the local genomic scale, TwoAYGGAY elements frequently occur in close proximity, often forming doublets (**Fig. 4F; fig. S22; Supplementary Text**). Of these doublets, 68.4% were divergently oriented, with a median inter-element distance of 249 bp (IQR, 163–312 bp). Both doublet formation and divergent orientation were significantly enriched relative to empirical null distributions accounting for the genomic distribution of individual TwoAYGGAY elements (*p* < 10^-4^ for both; **Methods**). The short 5’ repeat motif was also enriched upstream of TwoAYGGAY matches, with a mean of 6.18 copies within the upstream 200 bp compared with 2.84 in randomly sampled regions (**Fig. 4G; fig. S23; Methods; Supplementary Text**). Individual loci contained more than 30 upstream repeats in some cases. These repeats occurred in a consistent orientation and register upstream of the RNA match, although the spacing and precise position of the array relative to the 5’ boundary varied across loci (**Fig. 4H; figs. S23 and S24**).

Finally, we assessed whether the extended secondary structure occurs more broadly across prokaryotes by applying the Minerva-extended covariance model to all prokaryotic genomes in the Rfam database containing at least one match to the original TwoAYGGAY model. The Minerva-predicted extension was concentrated in Pseudomonadota (**Fig. 4I; fig. S25**). Within this phylum, we observed variants lacking the hairpin within the extended basal stem (**fig. S26**) and others in which the terminal 3’ hairpin was replaced by additional AYGGAY-motif-containing hairpins (**fig. S27**). By contrast, TwoAYGGAY matches in Bacillota generally lacked the extension and were instead predicted to adopt an elongated basal stem resembling the originally described architecture (**figs. S28 and S29; Supplementary Text;** *19*). Overall, these results reveal substantial structural variation among TwoAYGGAY RNAs across prokaryotic lineages, with the extended architecture largely confined to Pseudomonadota.

### Minerva reveals a variable ncRNA array associated with UG27 reverse transcriptases

Like TwoAYGGAY-containing regions, UG27 reverse transcriptase loci emerged from the Minerva genome scan with strong repetitive-motif and base-pairing signals, showing 11.3-fold enrichment in repetitive-motif signal and 6.5-fold enrichment in base-pairing signal relative to all intergenic regions. These reverse transcriptases belong to the class 2 family of Unknown Group reverse transcriptases, several members of which produce diverse cDNA products from nearby RNA templates (*43, 44, 49, 51, 52*). UG27 loci encode three conserved proteins, which we refer to as the reverse transcriptase (RT), small accessory protein (sORF), and large accessory protein (bORF), together with a long intergenic region previously hypothesized to contain an RNA template (**fig. S30;** *49*). The five UG27 systems in our 150-genome dataset were all located in integrated prophages in bacteria from the hCom2 community (**Fig. 5A**).

**Fig. 5.**
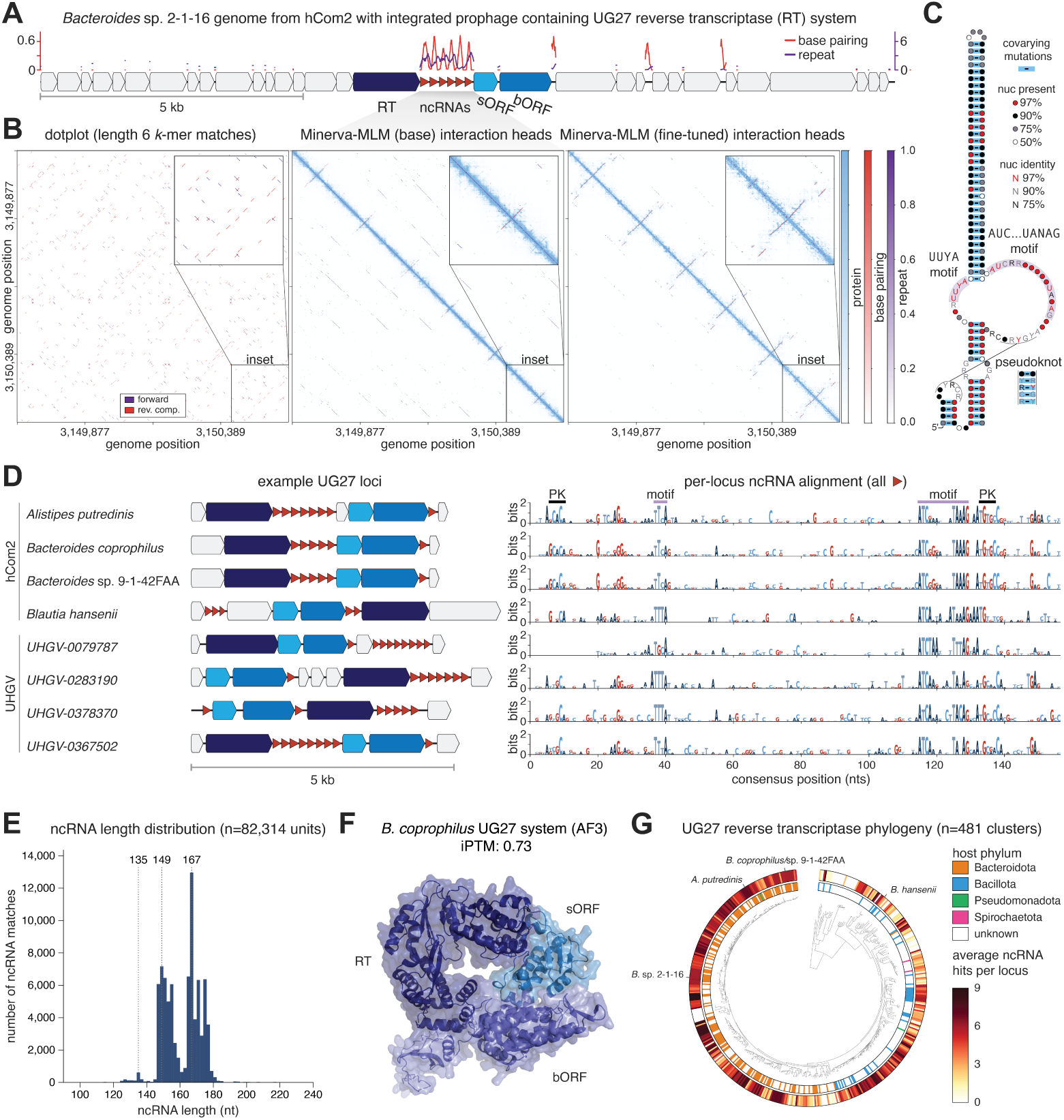
UG27 reverse transcriptase systems encode variable arrays of structurally conserved ncRNAs. **(A)** Locus visualization of the *Bacteroides* sp. 2-1-16 UG27 system with its surrounding prophage context and counts of Minerva-predicted base-pairing and repetitive-motif interactions plotted above. **(B)** Dotplot, Minerva-MLM base model, and Minerva-MLM fine-tuned interaction head predictions for an intergenic region of the *Bacteroides* sp. 2-1-16 UG27 system. The inset shows a zoom-in on one ncRNA unit and its surroundings. **(C)** Depiction of covariance model for UG27 ncRNA with additional features labeled. **(D)** UG27 loci examples from the Unified Human Gut Virome Catalog and the hCom2 bacterial community (left), with corresponding ncRNA alignments guided by the covariance model (right). **(E)** Length distribution of UG27 ncRNAs across 14,118 deduplicated loci (82,314 ncRNA units). **(F)** AlphaFold3-predicted structure of the three protein components in the *Bacteroides coprophilus* UG27 system. **(G)** Phylogenetic tree of UG27 reverse transcriptases clustered at 90% sequence identity with predicted bacterial host phylum and average number of ncRNAs per locus.

Using Minerva to interrogate individual UG27 loci, we noticed a striking pattern within the intergenic regions. Minerva-MLM interaction heads predicted multiple neighboring hairpins linked by repetitive-motif interactions, suggestive of a structured array of approximately 150-nt units (**Fig. 5B**). Family-specific finetuning of Minerva-MLM on UG27 loci further revealed an additional hairpin at the base of each ncRNA unit and a pseudoknot, while reducing the repetitive-motif signal (**Fig. 5B; figs. S31 and S32)**. The pseudoknot emerged after fine-tuning on as few as 10 UG27 loci, corresponding to approximately 60 individual array units, and was recovered even using rank-1 low-rank adaptation (LoRA), a highly parameter-efficient fine-tuning method (**fig. S32;** *53*).

We used the repetitive motifs and base-pairing interactions predicted by Minerva to guide alignment of the individual ncRNA units. Despite limited primary-sequence conservation, the resulting alignment supported a conserved secondary-structure architecture, enabled construction of a covariance model, and revealed short conserved sequence features within an asymmetric internal loop that interrupts the main hairpin and brackets the central stem (**Fig. 5C**). Neither these short motifs nor the broader array architecture was readily apparent from conventional sequence dotplots (**Fig. 5B**). Individual units were highly sequence-diverse both within the same array and across loci while retaining the common predicted secondary structure (**Fig. 5D**). Unit lengths were concentrated around 149 and 167 nt, with a smaller population of approximately 135-nt units lacking the pseudoknot (**Fig. 5E**). At the locus level, the order of the three UG27 protein-coding genes varied, and some loci contained additional ncRNA units in neighboring intergenic regions (**Fig. 5D**). AlphaFold3 further predicted that the three UG27 proteins form an interacting complex (**Fig. 5F; fig. S33; Methods;** *54* ), suggesting a system architecture comprising this protein complex and the variable ncRNA array.

To examine how UG27 systems are distributed, we constructed a phylogenetic tree of UG27 reverse transcriptases and annotated each locus with its associated ncRNA array (**Fig. 5G**). UG27 systems are predominantly found in bacteriophages predicted to infect Bacteroidota and Bacillota, with additional clades associated with Pseudomonadota and Spirochaetota. Both ncRNA copy number and unit length varied across the phylogeny, and in several clades Minerva predicted structured units not captured by the current covariance model, indicating sequence diversity beyond the model’s detection range (**fig. S34**). Across metagenomic databases, we also found closely related UG27 loci whose arrays differed by the presence or absence of one or two units, consistent with the gain or loss of individual ncRNA units (**fig. S35**).

### UG27 systems reverse transcribe ncRNA hairpins into short cDNA products

To test whether the UG27 ncRNA array serves as a template for cDNA synthesis, we cloned the five UG27 systems identified in the hCom2 genomes, expressed them heterologously in *E. coli* under a T7 promoter, purified DNA, and performed strand-specific DNA sequencing (**Fig. 6A**). These systems sample disparate branches of the UG27 RT phylogeny (**Fig. 5G; fig. S36**). Given the long and concatemeric cDNA products reported for other class 2 Unknown Group reverse transcriptases, we initially expected UG27 systems to produce similar products. Denaturing PAGE instead revealed short 50–100-nt ssDNA products from all five systems, whose recovery was enhanced by a modified miniprep (**fig. S37A**). These products required RT activity, as mutation of the RT active-site motif from YSDD to YSAA ablated their production (**Fig. 6B; fig. S37B**).

**Fig. 6.**
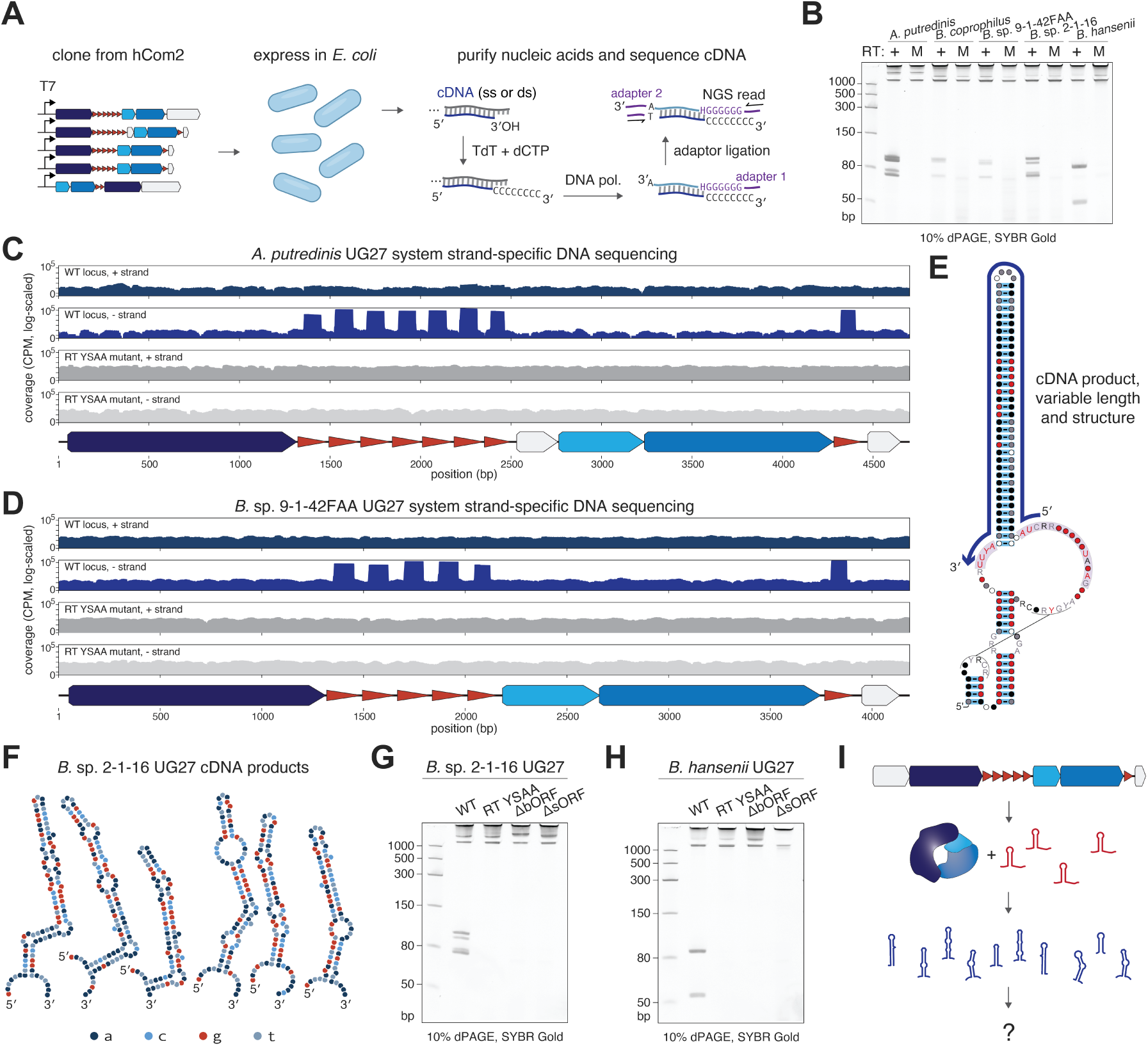
UG27 systems reverse transcribe the central hairpin of each ncRNA unit into short cDNA products. **(A)** Schematic of experimental approach used to identify UG27 system products by heterologous expression in *E. coli*, nucleic-acid purification, and strand-specific DNA sequencing. **(B)** Denaturing 10% TBE-Urea PAGE of purified nucleic acids, revealing abundant ssDNA products. M indicates constructs in which the RT active-site motif was mutated from YSDD to YSAA. **(C)** Strand-specific DNA-sequencing coverage, normalized to counts per million mapped read pairs (CPM), across the *Alistipes putredinis* UG27 locus for the wild-type system and catalytically inactive RT YSAA mutant. Positive- and negative-strand coverage is shown separately on a logarithmic scale. **(D)** Strand-specific DNA-sequencing coverage, as in **(C)**, for the *Bacteroides* sp. 9-1-42FAA UG27 system. **(E)** Location and boundaries of the cDNA product mapped onto the UG27 ncRNA covariance model. **(F)** ViennaRNA-predicted secondary structures of cDNA products generated by the *Bacteroides* sp. 2-1-16 UG27 system, using DNA thermodynamic parameters. **(G)** Denaturing 10% TBE–urea PAGE of nucleic acids purified from *E. coli* expressing the wild-type *Bacteroides* sp. 2-1-16 UG27 system, the RT active-site YSAA mutant, or constructs lacking the bORF or sORF. **(H)** Denaturing PAGE analysis, as in **(G)**, for the *Blautia hansenii* UG27 system. **(I)** Model of the UG27 system showing its three protein components, ncRNA array, and resulting short cDNA hairpins.

Strand-specific sequencing showed that these cDNAs mapped precisely to the central hairpins of the UG27 ncRNAs across all five loci (**Fig. 6, C and D; fig. S38**). RT-inactive controls showed approximately uniform plasmid coverage and recovered the endogenous *E. coli* Ec86 retron product as a positive control for ssDNA capture (**fig. S39**). The cDNA boundaries coincided with the conserved sequence motifs identified from the Minerva-guided alignment; reverse transcription began at the conserved AUC sequence within the AUCNNNNNNNUANAG motif and terminated after the conserved UUYA motif on the opposite side of the central hairpin (**Fig. 6E; fig. S38**). The resulting cDNAs are predicted to form diverse hairpins containing variable bulges and unpaired regions (**Fig. 6F; fig. S40)**.

To determine the protein requirements for cDNA synthesis, we deleted the two accessory ORFs in the *Blautia hansenii* and *Bacteroides* sp. 2-1-16 systems. Deletion of either accessory ORF abolished cDNA production (**Fig. 6, G and H**). Combined with the requirement for RT catalytic activity, these results indicate that all three UG27 proteins are required for cDNA synthesis during heterologous expression in *E. coli*. These findings support a model in which the three UG27 proteins reverse transcribe the central hairpin of each unit within the variable ncRNA array, generating short cDNA hairpins of diverse sequence and length (**Fig. 6I**).

## Discussion

We anticipate that Minerva will accelerate sequence analysis and discovery in three ways: (i) by enabling detailed interrogation of specific loci and their interactions, potentially enhanced by family- or group-specific fine-tuning; (ii) by supporting search and retrieval of related loci and guiding construction of downstream representations such as alignments and covariance models; and (iii) by systematically identifying genomic regions with previously unrecognized structure, including new ncRNAs and associated ribonucleoprotein systems.

TwoAYGGAY and UG27, both prioritized through genome-wide scanning, illustrate distinct limitations of conventional homology-based discovery. For TwoAYGGAY, the Rfam model did not capture lineage-specific structural extensions, while the UG27 ncRNA array was difficult to align and detect because its units are highly sequence-diverse and retain only short discontinuous conserved motifs. In both cases, Minerva predictions and family-specific fine-tuning enabled construction of improved covariance models, making alignments and sequence analysis tractable.

The resulting analyses also raise distinct biological questions. The AYGGAY loop motifs in TwoAYGGAY elements have been proposed as binding sites for Csr/Rsm family post-transcriptional regulators (*55*), and additional AYGGAY-containing hairpins in some variants suggest that these RNAs may act as multivalent Csr/Rsm sponges. Their extreme copy-number variation, flanking repeats, and enrichment as divergent doublets further point to a mobile-element-like mechanism of expansion, although any link between mobility and regulation remains unresolved. For UG27, the biochemical output is clear, but the biological role of its cDNA products remains unknown. We did not observe obvious sequence complementarity to phage or bacterial genomes, which leaves open whether these cDNAs recognize nucleic-acid structures, interact with proteins, or act through another mechanism. Apparent gains and losses of individual array units suggest an evolutionarily modular, potentially reprogrammable system in which cDNA sequence or structure may determine function.

Broadly, coevolutionary maps provide an interpretable interface between genome language models and biological discovery by resolving different types of interactions rather than assigning a single annotation to each sequence. Improving the underlying models will broaden what coevolutionary mining can discover. Minerva-MLM and other bidirectional genome language models have relatively short context windows, limiting their ability to detect long-range genomic interactions. Even within these windows, Minerva-MLM predicts protein contacts less accurately than specialized protein models, and RNA–protein interactions remain difficult to predict. In addition, Minerva-MLM’s representation of coding regions as amino acids prevents us from simultaneously modeling overlapping nucleotide-level features, such as structured RNAs embedded within coding sequences. Fine-tuning can sharpen interaction signals, potentially at the expense of others, as family-specific adaptation strengthened RNA secondary-structure predictions but attenuated repeat- and ORF-associated signals in our DRT2, TwoAYGGAY, and UG27 case studies. Comparing base and fine-tuned predictions helps retain both. Longer context windows, improved predictions for more interaction types, and deeper analysis of model internal representations may therefore extend Minerva toward more comprehensive annotation, search, and alignment.

Minerva functions as a discovery engine that generates hypotheses from sequence data for subsequent computational and experimental analysis. Our genome-wide scan predicts hundreds of uncharacterized structured loci per genome, including patterns outside the interaction types we have evaluated. We expect coevolutionary mining to become a broadly useful complement to homology- and alignment-based approaches, converting latent relationships learned by genome language models into interpretable, testable biological insights.

## Acknowledgements

We thank Daniel Chang, Lillian Brixi, Sofia Lövestam, Pranav Lalgudi, Jenny Shi, Aditi Merchant, Samuel King, Kiarash Jamali, Nathan Johns, Dennis Zhang, Jordan Hoff, Alex Li, Joshua Park, Owen Dunkley, Jonathan Strecker, Soumya Kannan, Guilhem Faure, Liana Merk, Matthew Durrant, Joseph Tey, Sergey Ovchinnikov, and all members of the Hie, Fischbach, and Deisseroth labs for helpful discussions. We also thank the Stanford Microbiome Therapies Initiative (MITI) for maintaining and providing strains from hCom2.

## Author Contributions

D.B.L. and G.B. conceived the study and jointly performed all experiments and analyses. A.S.K. contributed to the development of categorical Jacobian fingerprinting. M.B.F., N.N.I., and N.C.K. provided JGI data. C.L.D., S.A.E., and A.G. provided reagents and contributed to exploratory experiments on UG27 function. K.D., M.E.W., M.A.F., and B.L.H. supervised the study. D.B.L., G.B., M.E.W., M.A.F., and B.L.H. wrote the initial draft of the manuscript. All authors reviewed and approved the manuscript. D.B.L. and G.B. have agreed that either author may list their name first on their respective CVs and academic materials.

## Data, Code, and Materials Availability

Code for the Minerva framework is made available at https://github.com/garykbrixi/minerva, including for training, fine-tuning, and locus-level prediction. Minerva-MLM model checkpoints are shared on Hugging Face (<u>Minerva-MLM</u>, <u>Minerva-MLM-8k</u>).

## Materials and Methods

### Pretraining of Minerva-MLM and Minerva-MLM-8k

Minerva-MLM and Minerva-MLM-8k use the 650-million-parameter transformer architecture and mixed-modality tokenization introduced for gLM2 (*25*). Each model has 33 transformer layers with 20 attention heads per layer and a hidden dimension of 1,280 and uses rotary positional embedding (RoPE). Annotated protein-coding regions are represented as uppercase amino-acid tokens, and intergenic regions are represented as lowercase nucleotide tokens. <+> and <−> tokens at gene boundaries indicate gene orientation, and <+> is used preceding each intergenic segment.

We initialized Minerva-MLM from the pretrained gLM2 650M checkpoint and continued pretraining on a mixture of the Open MetaGenomic (OMG) corpus and the GlobDB release r226 dataset (*25, 33*). OMG is sampled at 25% and GlobDB is sampled at 75% of the training data mixture. Minerva-MLM was trained for 1.2 million steps with a context length of 4,096 tokens. We subsequently initialized Minerva-MLM-8k from Minerva-MLM, increased the RoPE base parameter from 10,000 to 20,000 and the context length to 8,192 tokens, and continued training for an additional 620,000 steps. Additional model and training hyperparameters are provided in **table S2**.

Under uniform prediction, the expected cross-entropy losses are approximately log(20) for amino-acid tokens and log(4) for nucleotide tokens. We initially weighted nucleotide-token loss by log(20)/log(4) ≈ 2.16 to equalize the two baselines. During preliminary training experiments, however, this correction was insufficient to prevent nucleotide loss from plateauing or increasing while the aggregate loss continued to improve. We therefore empirically increased the nucleotide-token loss weight to 4.16 for both Minerva-MLM and Minerva-MLM-8k, retaining a weight of 1 for amino-acid-token loss.

### Discovery benchmark datasets and evaluations

We evaluated Minerva on three classes of pairwise sequence relationships: protein monomer contacts, conserved RNA base pairing, and repetitive genomic motifs (**Fig. 1C; fig. S2**). For each benchmark, we divided examples into disjoint training and evaluation sets. The training sets were used to select reference fingerprints for categorical Jacobian fingerprinting and fit attention-based interaction heads, and performance was reported on the held-out evaluation sets. Because these benchmarks are necessarily constructed from previously characterized interactions, they are intrinsically biased toward known biological systems. We therefore restricted these comparisons to zero-shot scoring methods and lightweight probes, aiming to evaluate interaction information already encoded by the pretrained models rather than their capacity to learn benchmark-specific annotations.

#### Protein monomer-contact evaluation

To assess recovery of long-range protein monomer contacts, we used a curated subset of proteins in the trRosetta training set derived from Protein Data Bank (PDB) structures (*34*). Contact maps were computed using an 8 Å distance threshold between Cα atoms in the protein structure to define contacting residue pairs. We filtered the dataset to focus on prokaryotic proteins, identified through RCSB PDB and NCBI Taxonomy APIs to confirm prokaryotic origin (Bacteria or Archaea domains). From the filtered prokaryotic proteins, we used 118 proteins total, with 23 as a training set and 95 as an evaluation set (reflecting a 20/80 split), with proteins ranging from 200 to 1024 amino acids in length. Proteins were provided to each model directly without genomic context. We focused on long-range contacts, defined as those with at least 24 residues of sequence separation.

#### Conserved RNA base-pairing evaluation

We constructed an RNA base-pairing benchmark from the Rfam database. Beginning with all 4,178 Rfam families, we assigned each family a taxonomic-domain label using per-sequence taxonomy queries against the public Rfam MySQL mirror, with query logic adapted from the EMBL–EBI “Rfam/rfam-taxonomy” scripts (https://github.com/Rfam/rfam-taxonomy). Queries were run on May 16, 2025, separately for significant full-alignment hits and seed-alignment hits. Within each hit set, a domain was designated as the major domain if it accounted for at least 90% of classified hits; otherwise, the family was labeled as mixed, except when the only other category was unclassified sequences. Seed- and full-alignment assignments were combined as “ <seed>/<full>” when they differed. We retained families whose combined assignment contained Bacteria, including cases such as “Mixed/Bacteria” and “unclassified sequences/Bacteria”, but excluded “Bacteria/Eukaryota” families to avoid including families for which a small bacterial seed was not representative of the full alignment. The broadly conserved 5S rRNA, tRNA, and tRNA-Sec families were included regardless of their computed assignments. This procedure identified 1,093 bacterial families, compared with 1,030 assigned strictly to Bacteria (**fig. S2A**). Full multiple-sequence alignments in Stockholm format were available from the Rfam FTP mirror for 1,065 families; for the remaining 28, we retrieved the corresponding seed alignments from the Rfam website. Five families whose member sequences could not be retrieved through NCBI Entrez (RF02973, RF02988, RF03073, RF03097, and RF03113) were excluded. We filtered for successfully retrieved families with an alignment depth of at least 10 sequences, yielding a curated set of 680 bacterial RNA families (**fig. S2**).

For each family, we parsed the Rfam consensus secondary structure in WUSS notation to construct a reference map containing both nested and pseudoknotted base pairs and sampled without replacement up to 10 aligned sequences. For each sampled sequence, we parsed the source NCBI accession and alignment coordinates, retrieved the annotated region with 7,500 nucleotides of flanking sequence on each side from NCBI GenBank, and reverse-complemented minus-strand matches to match the orientation of the Rfam alignment. Consensus base pairs were projected from alignment coordinates onto each sequence, retaining a pair only when neither position aligned to a gap. The downstream extraction pipeline sampled one eligible sequence per family and required at least four reference base pairs with a sequence separation of at least three nucleotides. Five short families (four of which are CRISPR direct repeats) failed this contact requirement, and RF02541 was excluded because its 5,241-column alignment exceeded the 4,096-token extraction limit. RF00177, an approximately 1,980-nt bacterial small-subunit rRNA family, was additionally excluded from comparisons with RiNALMo because it exceeded that model’s context window.

The common comparison set therefore contained 673 families, of which 27 were assigned to training and 646 to held-out evaluation. For Minerva-MLM and gLM2, protein-coding regions in the retrieved genomic context were predicted de novo with Pyrodigal and represented as aminoacid tokens, whereas intergenic regions were represented as nucleotide tokens. The annotated ncRNA interval was excluded from ORF prediction to prevent it from being translated into amino-acid tokens. Predictions were evaluated only for nucleotide pairs within the annotated RNA boundaries. Thus, the benchmark tests whether a model can recover the conserved, family-level base-pairing architecture of an RNA from an individual member sequence.

#### Repetitive-motif evaluation

We constructed a benchmark of 50 genomic loci representing five classes of natural repetitive elements, with 10 loci in each class. The training set comprised 20 loci containing CRISPR arrays or type III toxin–antitoxin systems, whereas the held-out evaluation set comprised 30 loci containing REP elements, SsnA-associated non-template-strand (NTS) repeats, or TIGR-Tas systems. CRISPR repeats were identified using CRISPR Recognition Tool v1.2 (*56*) through the CRT plugin in Geneious Prime. Type III toxin-antitoxin loci were obtained from table S2 of (*57*) and TADB 3.0 (*58*). TIGR-Tas and SsnA-associated loci were obtained from (*8*) and (*59*), respectively. REP-containing loci were identified using RepRanger (*60*).

Repeat-unit boundaries within each locus were manually curated, guided by tool identifications. Corresponding positions across repeat units were designated as positive reference interactions, producing a two-dimensional ground-truth position-pair map for each locus. Predictions were evaluated over nucleotide pairs within the annotated intergenic region, excluding pairs separated by fewer than six nucleotides. Performance was summarized both across the complete held-out set and separately for each repeat-system class.

### Categorical Jacobian computation and optimization

For an input sequence of length L, we computed the categorical Jacobian as previously described (*28*). Briefly, each position was systematically perturbed to other token values, and the resulting changes in the model’s per-position output logits were recorded. This procedure produces a fourth-order coupling tensor J of shape (L, A, L, A), where the two alphabet dimensions A describe the input substitution and output-token response, respectively. That tensor, which is the categorical Jacobian, was symmetrized, mean-centered along each alphabet dimension, and reduced to an (L × L) contact map by taking the Frobenius norm over the two alphabet dimensions. Average product correction (APC) was then applied to the resulting L × L score matrix.

We evaluated two perturbation schemes. In full-substitution mode, each nucleotide was replaced with the other three nucleotide tokens, and each amino acid was replaced with the other 19 amino-acid tokens. The alphabet size (A) was 24 for Minerva-MLM, reflecting the four nucleotide and 20 amino-acid tokens in its mixed-modality vocabulary; 20 for ESM-2; and four for RiNALMo. The resulting alphabet-resolved coupling blocks were used for categorical Jacobian fingerprinting. In mask-only mode, each eligible position was instead replaced once with the mask token. This produces a computationally less expensive position-pair score but lacks the input-substitution axis required for categorical Jacobian fingerprinting.

The original implementation for gLM2 processed the perturbations associated with one sequence position in each model forward pass (https://github.com/TattaBio/gLM2/blob/main/categorical_jacobian_gLM2.ipynb). We introduced three optimizations. First, perturbations at multiple positions were combined into larger inference batches, reducing the number of forward passes required. Second, substitutions were restricted to tokens of the same modality: nucleotide positions were substituted only with nucleotides, and amino-acid positions only with amino acids. For a sequence containing 20% nucleotide and 80% amino-acid tokens, this requires approximately 0.2L×3 + 0.8L×19 ≈ 15.8L perturbed sequences, rather than testing every canonical sequence token at every position.

Additionally, we automatically skipped mutation of special tokens including beginning- and end-of-sequence tokens (BOS, EOS). Third, the logits or derived coupling values from each inference batch were transferred immediately to CPU memory, preventing the accumulation of intermediate results in GPU memory. The optimized and original implementations were compared in the computational-efficiency evaluation described below.

### Categorical Jacobian fingerprinting

Categorical Jacobian fingerprinting uses the alphabet-resolved structure of Jacobian coupling blocks to distinguish classes of pairwise interactions. We constructed separate reference fingerprints for RNA base pairs, repetitive nucleotide motifs, and protein contacts using labeled interactions from the corresponding training datasets. RNA base-pairing fingerprints were derived from eight Rfam families (RF00005, RF00010, RF00023, RF00162, RF00169, RF01854, RF02967, and RF04183), repetitive-motif fingerprints from the type III toxin–antitoxin set of sequences, and protein-contact fingerprints from the protein-contact training set. No held-out evaluation examples were used to construct the fingerprints or select similarity thresholds. For each sequence in a reference dataset, we extracted the coupling block for every position pair (i,j) with an APC-corrected Frobenius-norm score greater than 4.0 and a sequence separation greater than two positions for nucleotide interactions or six positions for protein interactions.

Each block had previously been symmetrized and mean-centered along both alphabet dimensions. We retained the modality-relevant portion of each block: (4 × 4) for nucleotide–nucleotide pairs and (20 × 20) for amino-acid–amino-acid pairs. Blocks were pooled across reference sequences, flattened into vectors, and hierarchically clustered using cosine distance and average linkage. For each interaction type, we manually selected a dendrogram cut that isolated a cluster enriched for the corresponding ground-truth interactions. The mean modality-specific coupling block within the selected cluster was used as the reference fingerprint (**figs. S3 and S4**).

After constructing the reference fingerprints, we applied categorical Jacobian fingerprinting to arbitrary query sequences to classify position pairs by interaction type. For each position pair (i, j), we extracted and flattened the modality-specific Jacobian coupling block and compared it to each reference fingerprint using cosine similarity. Nucleotide–nucleotide pairs were compared only with nucleotide fingerprints, and amino-acid–amino-acid pairs only with protein fingerprints. A pair was assigned to an interaction class when its maximum similarity to a reference fingerprint exceeded the threshold selected for that class: 0.35 for RNA base pairing, 0.20 for repetitive motifs, and 0.07 for protein contacts. If a pair exceeded the thresholds for multiple compatible classes, it was assigned to the class with the highest raw cosine similarity. Pairs that did not exceed any threshold were assigned to an “other” channel. This procedure produced separate RNA base-pairing, repetitive-motif, protein-contact, and other interaction maps for each query sequence. We evaluated categorical Jacobian fingerprinting with Minerva-MLM for all three interaction classes, RiNALMo for RNA base pairing and repetitive motifs, and ESM-2 650M for protein contacts.

To examine the physicochemical patterns represented by the protein-contact fingerprints, we compared fingerprints derived from Minerva-MLM and ESM-2 with the Miyazawa–Jernigan statistical potentials (*37, 38*), following an analysis inspired by (*40*). Miyazawa-Jernigan potentials were symmetrized by copying the upper triangle to the lower triangle, corrected by APC, and sign-flipped for comparisons. We performed eigendecomposition separately on each symmetrized (20 × 20) matrix and constructed rank-one matrices from each of the four eigenmodes with the largest absolute eigenvalues. The resulting modes were compared qualitatively with respect to amino-acid hydrophobicity, charge, side-chain size, and residue-specific interactions, including cysteine–cysteine interactions (**fig. S5**).

### Interaction head and attention head analyses

For individual attention-head comparisons, we extracted attention maps from all layers and heads of each model and symmetrized each map by averaging it with its transpose. Individual attention heads were evaluated directly against the corresponding reference interaction maps, and the best-performing head for each interaction class was selected using only the associated training set. Following previous approaches for protein-contact prediction, we constructed interaction heads by training logistic regression models using the concatenated, symmetrized attention values from selected layers as features for each eligible position pair (*27*). We fit separate elastic-net logistic-regression models for each interaction class, evaluating inverse regularization strengths C in {0.01, 0.1, 0.15} and L1 ratios in {0, 0.5, 1}. Final configurations were selected to maximize AUPRC on the corresponding training set. The selected configurations were C=0.01 and L1 ratio=1.0 for protein monomer contacts, C=0.1 and L1 ratio=1.0 for RNA base pairing, and C=0.15 and L1 ratio=0.0 for repetitive motifs.

Extracting attention maps from a given layer requires instantiation of the complete attention matrix, which increases memory usage and prevents the use of FlashAttention kernels (*61*). We therefore compared interaction heads constructed from the last two or last six transformer layers (**fig. S7A**). We used the last-two-layer interaction heads for all figure panels. For combined visualization of all three interaction heads, each position pair was colored by its highest-scoring interaction head, with RNA base-pairing scores above 0.5 taking precedence.

### Performance evaluations and ablations

#### Evaluation metrics and model comparisons

We evaluated predictions on the held-out protein monomer-contact, conserved RNA base-pairing, and repetitive-motif datasets described above. Scores were calculated over all eligible position pairs after excluding pairs separated by fewer than 24 residues for proteins or six nucleotides for RNA and repetitive motifs. The primary performance metric was area under the precision-recall curve (AUPRC). We also calculated area under the receiver operating characteristic curve (AUROC) and precision at T (P@T), where T is the number of positive reference interactions in an example. For the protein-contact evaluation, we additionally calculated precision among the top L, L/2, and L/5 predictions (P@L, P@L/2, P@L/5), where L is the protein length. Aggregate performance was calculated by averaging across examples.

We evaluated gLM2 and Minerva-MLM on all three benchmarks. ESM-2 650M was included as a specialized baseline for protein monomer-contact prediction, and RiNALMo was included as a specialized baseline for RNA base-pairing prediction. Protein sequences were provided to each model without genomic context. For the primary RNA base-pairing comparison in **Fig. 1E**, when evaluating RiNALMo, RNA sequences were provided in isolation without flanking DNA context, a setting favorable to RiNALMo. Additionally, because RiNALMo is trained with the RNA oriented as in Rfam, we performed predictions for both the accession-oriented sequence and its reverse complement. We then took the better performance of the two orientations for each model on the base-pairing evaluation, applied equally to all models. Repetitive motifs were evaluated in their native genomic contexts.

We compared four interaction-extraction approaches: scalar categorical Jacobian scoring using the masked-token-mutation mode, categorical Jacobian fingerprinting, individual attention heads, and logistic-regression interaction heads. Reference fingerprints and interaction heads were selected or fit using only the corresponding training sets, and performance was reported on held-out examples. The best individual attention head was selected using the training set, independently for each interaction type. For comparisons across individual Rfam families, methods were ranked by AUPRC within each held-out family.

#### Sequence-orientation analysis

To assess sensitivity to sequence orientation, each RNA in the base-pairing evaluation was evaluated in both the forward and reverse-complemented orientations with respect to the genome accession. Predictions from reverse-complemented inputs were transformed back to the coordinates of the original sequence before comparison with the reference base-pairing map. We compared orientation-specific performance for the best individual Minerva-MLM attention head and RiNALMo categorical Jacobian fingerprinting (**fig. S11B**).

#### Genomic-context ablation

To determine how surrounding genomic context affected RNA base-pairing prediction, we evaluated each Rfam RNA in isolation and with up to 500 or 2,000 bp of flanking sequence on each side. Inputs were centered on the annotated RNA and truncated to maximum total lengths of 1,024 or 4,096 tokens, respectively. For Minerva-MLM, flanking regions were represented using its mixed-modality genomic tokenization. Because RiNALMo accepts nucleotide sequences, its flanking regions were supplied entirely as nucleotide tokens. Minerva-MLM was evaluated using its interaction head, whereas RiNALMo was evaluated using scalar categorical Jacobian scoring and categorical Jacobian fingerprinting. Predictions were scored only over nucleotide pairs within the annotated RNA boundaries (**fig. S11C**).

#### Language-modeling loss and prediction performance

To test whether language-modeling loss could serve as a measure of interaction-prediction confidence, we first compared the mean nucleotide validation loss of gLM2, Minerva-MLM, and Minerva-MLM-8k with their aggregate performance on the three interaction benchmarks (**fig. S11D**). We then calculated the cross-entropy loss and interaction-head AUPRC for each sequence in the held-out Rfam evaluation and tested their association using Pearson correlation (**fig. S11E**).

#### Computational efficiency evaluation

We benchmarked the computational cost of interaction prediction using the Minerva-MLM model on a single NVIDIA H100 GPU. We compared the original categorical Jacobian implementation as implemented for gLM2 in both substitution and masked mode (https://github.com/TattaBio/gLM2/blob/main/categorical_jacobian_gLM2.ipynb), our optimized Jacobian implementation in both substitution and masked mode, and interaction heads. Runtime was measured using time.perf_counter() with explicit CUDA synchronization immediately before and after each operation. Between measurements, GPU memory was cleared with torch.cuda.empty_cache() and Python garbage collection was invoked to prevent memory accumulation effects. We evaluated sequences of length 1024, 2048, 4096, and 8192 tokens. For interaction heads, we reported amortized per-sequence runtime using a batch size of 16 for inputs up to 4,096 tokens and a batch size of 8 for 8,192-token inputs.

### Family-specific fine-tuning of Minerva-MLM

We explored two fine-tuning strategies: (1) full-parameter updates with a learning rate of 5e-5, linear warmup over 100 steps, and weight decay of 0.01, and (2) low-rank adaptation (LoRA) with a learning rate of 1e-4, no warmup, dropout of 0.05, and ranks of 1, 16, or 32, with alpha set to twice the rank. LoRA adapters were applied to all attention projections (wqkv, wo), feed-forward layers (w1, w2, w3), and the language-model head.

Unless otherwise specified, fine-tuning used bfloat16 mixed precision, gradient checkpointing, an effective batch size of 16, a context length of 4,096 tokens, and AdamW optimization for 12,000 steps with the HuggingFace Trainer. Nucleotide-token loss was weighted by 2.16 relative to amino-acid-token loss, corresponding to the vocabulary-size correction described above. This differs from the empirically increased weight of 4.16 used during continued pretraining. To evaluate whether fine-tuning was feasible on commonly available hardware, we additionally compared full-parameter fine-tuning and LoRA on a free Google Colab instance with an NVIDIA T4 GPU using float16 precision.

We fine-tuned Minerva-MLM separately on DRT2, UG27, and TwoAYGGAY loci, producing models specialized for each of the three types of systems. For each system type, we constructed a fine-tuning dataset by curating a training list of loci, selecting a window of 4096 tokens for each locus centered on the system of interest. DRT2 loci were identified in JGI/IMG using a DRT2 profile HMM and deduplicated by exact window sequence identity, yielding 13,034 loci. UG27 loci were collected as described below and deduplicated at 100% identity for the three protein components, yielding 2,549 loci. TwoAYGGAY loci were selected from RF01731 covariance-model matches in complete genomes from the *Pseudomonas* Genome Database, release PGD r22.1, with an E-value cutoff of 0.01, yielding 6,884 loci.

To assess the effects of fine-tuning, we generated interaction maps using both categorical Jacobian fingerprinting and the pre-trained interaction heads (regressed on the base model) and compared them with predictions from the base Minerva-MLM model. For DRT2, predicted base-pairing interactions were additionally compared with the reference interactions derived from the available cryo-EM structures. Canonical base-pairing interactions in the DRT2 ncRNA were identified using the Python implementation of FR3D (*62*). Structures (PDB 9C0I and 9C0J, representing two conformational states) were parsed from mmCIF files, and all pairwise nucleotide interactions were annotated using FR3D’s pairwise interaction classifier with the 2023 geometric cutoff parameters and idealized hydrogen-bond templates; only cis Watson-Crick/Watson-Crick (cWW) base pairs were retained for downstream analysis.

#### Fine-tuning data-efficiency analysis

To evaluate the effect of DRT2 dataset size, we fine-tuned on randomly sampled subsets of 10, 100, 200, 500, or 1,000 loci, in addition to the complete set of 13,034 loci (**fig. S13**). To assess the effect of UG27 dataset size, we used the collection deduplicated at 100% identity for the three protein components and randomly sampled subsets containing 1, 10, or 100 loci, in addition to the complete set of 2,549 loci (**fig. S32**). For both systems, models were fine-tuned on the complete dataset for 12,000 training steps; the 10-, 100-, 200-, 500-, and 1000-locus subsets for 2,000 steps; and the single-locus dataset for 100 steps, as applicable.

### Genome selection, annotation, and scanning

We collected 150 bacterial genomes representing 116 common human gut bacteria from hCom2 (*46*), 25 pathogens and select agents, and 9 common model organisms, listed in **table S1**. Each genome was annotated using mettannotator v1.5.0 (*45*) in fast mode with Bakta v1.11.4 (*63*) as the gene caller, run via Nextflow v25.10.4 (*64*) with Singularity containers. In fast mode, the pipeline executes gene prediction (Bakta), functional annotation via InterProScan (*65*) and eggNOG-mapper v2.1.11 (*66*), biosynthetic gene cluster prediction via antiSMASH v8.0.1 (*67*) and GECCO v0.9.8 (*68*), pseudogene detection via Pseudofinder v1.1.0 (*69*), antimicrobial resistance gene detection via AMRFinderPlus v4.0.23 (*70*), CRISPR-Cas detection via CRISPRCasFinder v4.3.2 (*71*), carbohydrate-active enzyme annotation via dbCAN v5.1.2 (*72*), ncRNA detection via Infernal v1.1.5 (*14, 17*), tRNA detection via tRNAscan-SE v2.0.9 (*73*), and assembly quality assessment via QUAST v5.2.0. Defense systems were detected using DefenseFinder (*74*). The per-genome GFF3 annotations and DefenseFinder results were consolidated into annotated GenBank files used for downstream analysis and for tokenizing each genome sequence.

Each genome was parsed from GenBank format and tokenized into a mixed-token sequence. Tokenized genomes were tiled into overlapping chunks of 4096 tokens with a stride of 2048 tokens (50% overlap). We ran the Minerva model across all 150 genomes, saving the output predictions from the three last-2-layer interaction heads (repetitive-motif, base-pairing, and protein-monomer) using a single H100 GPU. This analysis was completed within 100 minutes on the H100 GPU, plus 31 minutes for writing the output maps. Genome windows were stored as sparse H5 files with uint16, retaining only values above a per-head threshold (0.1 for repeat or base-pairing, 0.5 for the protein head). To save storage, predictions in protein-coding regions can be masked, resulting in a total storage of 18 GB, 6% of the storage needed for maps that contain coding contacts.

### Intergenic-region, multi-hairpin, and mistokenized ORF analyses

#### Aggregation of intergenic-region predictions

We aggregated the predicted coevolutionary maps from Minerva interaction heads by intergenic region to prioritize loci for follow-up analysis. An intergenic region was defined as the complete nucleotide sequence between adjacent annotated coding sequences (CDSs). For each region, we counted the number of off-diagonal interaction head predictions that exceeded a set threshold (0.9 for base-pairing, 0.5 for repeats). Because genomes were scanned in overlapping tiles, an intergenic region could be split across or represented in multiple tiles. In these cases, we selected the tile with the highest count of predicted interactions as the representative tile of that region. A region was considered annotated if it overlapped any existing annotation; only regions without annotation overlap were classified as unannotated. We also visualized the predicted base-pairing and repetitive motif interactions as one-dimensional tracks by counting the interactions of each type per position and plotting a rolling mean of 21 nucleotides.

#### Enrichment at known systems and near nucleic-acid-interacting proteins

Using these predicted interactions per region, we compared the prioritization of known relevant systems as validation and of new systems for prospective discovery. We examined the repeat and base-pairing counts per annotation type, as defined above. We then compared base-pairing and repetitive-motif interaction counts across annotation classes. UG27 annotations were derived from the covariance model described below. In one of the five UG27 loci identified among the 150 scanned genomes, an annotated CDS overlapped three UG27 ncRNA covariance-model matches. We removed this CDS annotation during post-processing and retokenized the corresponding sequence as nucleotides so that the structured ncRNA array could be analyzed.

We further tested whether particular protein annotations, as named by mettannotator, were enriched in association with base-pairing and repetitive-motif signals from Minerva. Interactions within neighboring intergenic regions extending up to 500 bp upstream or downstream of each CDS were assigned to the corresponding protein. For each protein annotation, the distribution of neighboring interaction counts was compared with the background distribution across all annotated proteins in the 150 genomes using a one-sided Mann–Whitney U test. We chose several well-known nucleic-acid-interacting protein families to visualize (**fig. S14**).

#### Embedding-based analysis of intergenic regions

To use the Minerva-MLM embeddings to group different highly structured intergenic regions, we used UMAP visualization. Regions in the top 10% by either base-pairing or repetitive-motif interaction count were passed through Minerva-MLM with their surrounding genic context. The 1,280-dimensional hidden representations from the final layer were mean-pooled across nucleotide tokens within each intergenic region, excluding protein and orientation-marker tokens, and L2-normalized. The resulting embeddings were projected into two dimensions using UMAP (n_components=2, n_neighbors=15, min_dist=0.3). Points were additionally colored by annotations of interest including species, GC content, and per-region base-pairing and repeat scores (**fig. S15**).

#### Multi-hairpin enrichment analyses

We calculated how many putative non-coding elements are predicted by Minerva within and outside of previous annotations across the 150 genomes. We defined a multi-hairpin locus as hairpins (at least five contiguous anti-diagonal base-pairing interactions over 0.9) that are within 50 nt of each other. These elements were assigned as annotated if any part of the span covers an existing annotation, and we compared the number and percent of annotated versus unannotated elements at different numbers of hairpins.

To test statistical significance of the number of predicted putative non-coding elements against an empirical null, we adopted the approach from RMARK3 (*14*) and created compositionally matched genomes using an HMM. We randomly selected five genomes for this test and constructed 25-state HMMs for each genome fit to the intergenic sequences by Baum-Welch. For each genome we then generated a paired null where every intergenic region was replaced with sequences of equal length emitted by the HMM. We predicted interaction maps for these genomes using Minerva and compared the fold enrichment and empirical FDR between the true and null genomes. Fold enrichment was calculated as the ratio of predicted elements in real versus null genomes, and the empirical FDR as the null count divided by the real count. We found that the loci containing ≥2 hairpins are enriched by 14-fold against the null with an FDR = 0.07, while the loci containing ≥4 hairpins are enriched 132-fold with an FDR = 0.008.

#### Mistokenized ORF analysis

When inspecting Minerva-MLM interaction head predictions from the 150-genome scan, we visually identified mistokenized ORFs by their periodic triplet patterns. We extended putative misannotated ORFs to upstream ATGs with no intervening in-frame stop, producing candidate full protein annotations. These refined protein annotations were then searched using BLASTp against clustered_nr to confirm they match known proteins.

### TwoAYGGAY locus identification and covariance-model construction

Covariance models for extended TwoAYGGAY RNAs were built iteratively using Infernal, guided by Minerva-MLM base-pairing interaction head predictions. The covariance model of TwoAYGGAY in SBW25 (**fig. S18B**) was generated from 41 full-length TwoAYGGAY RNA matches within SBW25 and visualized using R2R. To construct the main covariance model (**Fig. 4D**), complete *Pseudomonas* genome assemblies were downloaded from the *Pseudomonas* Genome Database (PGDB), release pgd_r_22_1 (<u>pseudomonas.com</u>, *75*). An initial search was performed with the Rfam RF01731 (TwoAYGGAY) covariance model, obtained directly from Rfam and indexed with cmpress, using Infernal cmsearch. Minerva-MLM base-pairing interaction head predictions over a randomly subsampled collection of hits were used to guide the construction of a hand-curated seed alignment. This seed alignment was expanded to other homologs with cmalign, and iteratively improved. The resulting main Minerva-extended covariance model was used for subsequent searches and all other figure panels, and visualized using R2R.

For comparisons across *Pseudomonas* species, we searched the same set of complete *Pseudomonas* genome assemblies from PGDB with our final covariance model using cmsearch. Covariance-model hits were retained if their E-value was below 1 × 10^-3^. Because we sometimes observed weaker, overlapping partial hits on the opposite strand at the same genomic locus, antisense partial hits were removed when they overlapped a stronger, opposite-strand hit at the same locus by more than 50% of their length. Strain-level taxonomic metadata (species and NCBI lineage) was obtained for every genome in the database, including genomes with zero hits. Each genome was assigned to a phylogenetic species group based on the “species group” rank in its NCBI lineage (falling back to the species rank when no species-group rank was available), and genomes were grouped into six major species groups (*Pseudomonas aeruginosa, putida, chlororaphis, syringae*, and *fluorescens* groups, and the *Stutzerimonas stutzeri* group) with all remaining taxa combined into a single “other” category.

To examine the prevalence of the extended secondary structure among genomes containing RF01731 matches, we used cmsearch with the Minerva-extended covariance model to scan across the 847 complete, non-eukaryotic genome accessions listed in the Rfam RF01731 sequence table (https://rfam.org/family/RF01731#tab-sequences). Hits against the RF01731 stock covariance model were calculated using Rfam’s family-specific gathering cutoff. To be considered a match to the extended model, hits were required to pass the 1 × 10^-3^ E-value cutoff and filtering as described above, and additionally required to cover at least 80% of the Minerva-extended covariance model, with model coverage computed as the fraction of the covariance model spanned by the alignment to the genomic hit. Each hit was then grouped into “extended CM only”, “Rfam CM only”, or “both” categories, depending on whether there were overlapping hits from each model on the same accession and strand, with any nucleotide overlap. Each accession was assigned a taxonomic lineage (phylum through genus) using the NCBI taxonomy database, and hit-class counts were aggregated at each taxonomic rank, using parent-aware grouping so that unclassified taxa with different parent lineages were not conflated. Minerva-MLM base-pairing interaction head predictions for individual genomes were manually examined, and predicted secondary structures were visualized using R2R.

### TwoAYGGAY genomic architecture analyses

Minerva-extended TwoAYGGAY covariance model hits on the same genome and accession from PGDB were merged into loci by transitively grouping hits whose midpoints fell within 2 kb of one another, and each locus was assigned a multiplicity (singleton, doublet, triplet, or quadruplet-and-above) based on its number of constituent hits. For loci containing exactly two hits, the pair’s relative orientation was classified as convergent, divergent, or tandem based on the strands of the two hits. Using this locus structure, we characterized the distribution of locus multiplicities and the intergenic distance between consecutive hits within multiplet loci, genome-wide inter-hit distances independent of locus assignment, inter-hit distances within multiplet loci stratified by locus size, and doublet orientation frequencies and their associated inter-hit distances.

To analyze the enrichment of TwoAYGGAY doublets, we constructed an empirical null. The null distribution was created from the same PGDB genomes, keeping the coding genes but randomly placing the TwoAYGGAY RNA elements inside the intergenic regions, avoiding direct overlaps of multiple elements. We repeated this 10,000 times for each genome and then compared this null against the observed distributions of TwoAYGGAY matches. We verified our sampler against brute-force enumeration to confirm its correctness, which allowed us to perform this analysis on the 1,083 PGDB genomes. Because our null does not account for RNAs falling within CDS, we excluded the 1,693 observed TwoAYGGAY hits (out of 16,086) which fall within CDS to avoid bias for this significance testing and other analyses, leaving 14,393 elements from 1083 genomes. Enrichment was assessed by Monte Carlo, with the empirical *p*-value calculated as (1+r)/(1+n), where n is the total number of null replicates and r is the number of replicates reaching or exceeding observed values; with n=10,000, the smallest attainable *p*-value is *p* <1 × 10^-4^. Doublets were enriched over the null (*p* <1 × 10^-4^). This was corroborated by a Wilcoxon signed-rank test comparing observed doublet count against the respective null, which was performed using the 715 genomes with at least two TwoAYGGAY elements (*p* <1 × 10^-83^). Because the 2 kb grouping distance is arbitrary, we repeated the test across distance thresholds and found significant enrichment at every value up to 5 kb (*p* < 10^-4^ throughout), indicating that the result does not depend on the particular cutoff chosen. Triplets and larger clusters were enriched relative to independent placement, but after conditioning on the observed excess of doublets they occurred no more often than expected ([6.7% vs 6.4%], *p* = 0.38), consistent with the doublet being the only architecture independently enriched.

We further tested whether the two elements within TwoAYGGAY doublets were oriented independently. Under independent strand assignment, divergent, convergent, and tandem configurations are expected at frequencies of 25%, 25%, and 50%, respectively. Divergent pairs instead accounted for 68.4% of doublets (2,164 of 3,166), a 2.7-fold enrichment over this expectation (χ² goodness-of-fit test, df = 2, χ² =3263.4, *p* < 10^-300^).

### TwoAYGGAY repeat analyses

Repeats in SBW25 TwoAYGGAY RNA matches were annotated in Geneious Prime guided by Minerva-MLM repetitive-motif interaction head predictions, and motif logos were generated using WebLogo 3 in Python. To examine the broader prevalence of the 5′ repeat motif, we examined hits from PGDB as identified above. For each hit, we defined an oriented, strand-aware window spanning 200 bp upstream of the ncRNA’s 5′ end plus 50 bp into the 5′ end of the ncRNA itself and searched this window for a curated set of six 5-mers (GATCT, GCTCT, GCTTT, GCTGT, GCCGT, and GCAGT) matched as a single regular-expression union. This yielded, for every hit, the number and 5′-relative genomic position of all motif occurrences within the 200-bp upstream region. To assess enrichment relative to genomic background, an equal number of 200-bp windows (plus the same 50-bp boundary extension) were drawn at random from the same set of genome sequences, with the probability of drawing a given sequence proportional to its length; the same motif search was applied to these null windows. Observed and null distributions were compared for the number of motif occurrences per window, the positional density of motif occurrences across the window, and the spacing between consecutive motif occurrences within a window.

Motif counts were visualized by locus-multiplicity class and doublet orientation, including the minimum and maximum motif counts within each doublet (**fig. S23**). Upstream motif match positions were additionally represented as binary presence/absence matrices, with hits as rows and upstream match positions as columns. Rows were hierarchically clustered using Hamming distance and average linkage. Clustered matrices were generated for the complete set of Minerva-extended TwoAYGGAY hits in PGDB, the 1,000 Minerva-extended TwoAYGGAY hits with the greatest numbers of upstream motif occurrences, and for several individual genomes of interest, with rows colored by locus multiplicity and doublet orientation (**fig. S24**).

### UG27 loci collection

To identify UG27 loci from databases, we constructed profile HMMs using HMMER3 for each of the three protein components (RT, small accessory protein, and large accessory protein) from the UG27 loci provided in the supplement of (*49*). We searched these HMMs against predicted protein sequences from JGI IMG, the Unified Human Gut Virome catalog (*76*), the Meta-virus resource (MetaVR, *77*), and the Chinese Gut Viral Reference (78), retaining hits with a per-sequence bit score ≥120. Colocalized loci were defined as scaffolds carrying at least two of the three components within 20,000 bp, yielding 14,650 MetaVR, 3,627 JGI, 1,295 UHGV, and 691 CGVR loci prior to cross-database deduplication (Figure S31). Cross-database deduplication was performed at 100% nucleotide identity across the locus window, totaling 14,708 loci. We used JGI, UHGV, and CGVR for Minerva-MLM fine-tuning and loci from all four databases for bioinformatic analysis.

### UG27 ncRNA covariance model construction and array annotation

We constructed an initial alignment of UG27 ncRNA units by manually defining unit boundaries from the repetitive-motif and base-pairing interactions predicted by Minerva in UHGV UG27 loci. The aligned units were used to build an initial covariance model using Infernal (*14*), which was iteratively refined through repeated searches of the UG27 locus collection, realignment of newly identified units, and manual inspection of sequence and predicted secondary-structure conservation. The final covariance model was visualized using R2R (*79*). Because unit length and base pairing along the primary hairpin vary substantially across the family, a single covariance model did not capture the full diversity of UG27 ncRNAs. We therefore constructed a second covariance model for a clade in which both Minerva predictions and manual inspection supported an alternative structure lacking the pseudoknot.

We searched the UG27 locus collection, deduplicated based on 100% nucleotide identity, with both the pseudoknot-containing and pseudoknot-lacking covariance models. Overlapping matches from the two models were resolved by retaining the match with the lower E value, with E-value thresholds of 0.01 used to define an ncRNA unit. We then grouped neighboring units associated with the same UG27 protein-coding locus into ncRNA arrays. This procedure identified 82,314 ncRNA units across 14,118 UG27 loci. Unit length was calculated from the complete covariance-model match, including insertions relative to the consensus model. Comparison of individual arrays was done using sequence alignment with MAFFT in E-INS-i mode (*80*) in Geneious Prime. Alignments of individual ncRNAs in a given locus were constructed using cmalign against the covariance model.

### UG27 phylogenetic and metadata analysis

UG27 loci were deduplicated at 90% RT amino-acid identity using MMseqs2 easy-cluster (--min-seq-id 0.90 -c 0.80 --cluster-mode 2), yielding 649 cluster representatives (*81*). Of these, 481 contained all three protein components (RT, sORF, and bORF), as detected by their respective profile HMMs, and were retained for tree building. For each representative, we validated the protein matches by re-predicting ORFs from the source contig with pyrodigal in metagenomic mode and scanning the resulting proteins against the three component HMMs using pyhmmer (bit-score thresholds 30/20/20 for RT/bORF/sORF). Because some phage contigs encoded additional RTs, including RTs associated with diversity-generating retroelements, noncanonical RT matches outside the expected length range of 380–450 amino acids, including clear fragments (<250 aa), were replaced with the highest-scoring canonical-length RT immediately upstream of either accessory-protein gene on the same contig. The corrected RT sequences plus a DRT2 outgroup were aligned with MAFFT --auto (482 sequences × 1413 columns) and a maximum-likelihood tree was inferred with IQ-TREE v3.1.1 under LG+G4 with 1000 ultrafast-bootstrap and 1000 SH-aLRT replicates. The tree was rooted on DRT2, which was removed from the final visualization.

UG27 ncRNAs were annotated using the two covariance models described above. For each locus, we quantified the number and lengths of ncRNA hits and plotted these measurements alongside the tree. Visual inspection of Minerva predictions suggested that the absence of ncRNA hits in some clades reflects diversified structural units rather than the absence of ncRNA from these loci. For prevalence and host-range analyses, loci were deduplicated at the species level using existing vOTU assignments for MetaVR and UHGV and 95% ANI for JGI/IMG and CGVR. Host taxonomy and geographic origin were obtained from source database metadata: UHGV consensus host lineage (host_lineage_gtdb_r207), falling back to CRISPR-spacer-based prediction where the consensus was empty; JGI/IMG GTDB lineage and geographic fields; and MetaVR per-vOTU host_taxonomy and geographic_location. CGVR loci carry no host or geography metadata. Host phylum was parsed from the GTDB p token, and country strings were mapped to one of six continents by case-insensitive keyword match. For each 90% RT cluster, the cluster representative was assigned the modal host phylum and continent across its member loci; representatives without metadata-bearing members were marked unknown. Geographic distributions were summarized at the continent level (**fig. S34**).

### UG27 protein analysis and AlphaFold3 structure prediction

To assess conservation of the three UG27 protein components, protein sequences for the RT, bORF and sORF were deduplicated using MMseqs2 (*81*) (--min-seq-id 0.90 -c 0.80 --cov-mode 1) and aligned separately using MAFFT (*80*). Sequence logos were generated from the resulting alignments using logomaker (v0.8.7) (*82*), converting alignment column counts to information content. Conserved protein domains were identified with InterProScan version 5.78-109.0 (*65*) using Pfam (PfamA), PROSITE patterns, PROSITE profiles, CDD, SUPERFAMILY, Gene3D, SMART, NCBIfam, and PIRSR databases. The RT contained matches to Pfam reverse transcriptase family PF00078 and the retron-type reverse transcriptase conserved domain cd01646, whereas no conserved domains were detected in either accessory protein (**fig. S30**). Pairwise amino-acid sequence identity among the RT, bORF, and sORF proteins from the five hCom2 UG27 systems was computed from alignments of proteins from all five loci with MAFFT in E-INS-i mode and the BLOSUM62 scoring matrix (*80*) in Geneious Prime (**fig. S36**).

For structure prediction of the UG27 complex, we used AlphaFold 3 with the Singularity container and pre-computed databases (https://github.com/google-deepmind/alphafold3; (*54*)) and two MSA strategies: AlphaFold 3’s default database search (jackhmmer against UniRef90, MGnify, and BFD) and a custom paired MSA from co-localized loci (4,215 sequences per chain which contain all three proteins, deduplicated to 2,211 loci by 100% sequence identity). To construct paired MSAs, we extracted the three UG27 protein components from each locus and aligned them individually using MAFFT. We constructed concatenated alignments with the RT, sORF, and bORF proteins and then deduplicated identical protein sequences from the MSA to be used by AlphaFold3. For each of the five hCom2 UG27 loci, five random seeds with five samples each of all three proteins were generated. The highest-ranked prediction was selected using AlphaFold 3’s global ranking score (**fig. S33**).

### Cloning of UG27 system expression constructs

Stellar chemicompetent cells were prepared using Mix & Go (Zymo, T3001). Genomic DNA of the 5 strains in hCom2 containing UG27 systems was extracted from glycerol stocks using the Monarch Spin gDNA extraction kit (NEB, T3010S). UG27 systems were amplified using NEBNext High-Fidelity 2X PCR Master Mix (NEB, M0541S) and cloned into a T7 expression plasmid by Gibson assembly following PCR purification using a QIAquick PCR purification kit (Qiagen). Constructs containing gene deletions or RT active-site mutations were generated using Gibson assembly or KLD mutagenesis. All constructs were sequence-verified by Plasmidsaurus sequencing. Constructs and primers used for experiments are listed in **table S3**.

### Heterologous expression, DNA extraction, and denaturing PAGE

BL21(DE3) chemicompetent cells were prepared using Mix & Go (Zymo, T3001). Plasmids were transformed into BL21(DE3) bacteria, plated on LB-carbenicillin plates, and grown overnight. One colony was picked into 5 mL of MagicMedia expression medium (Invitrogen, K6803) and grown for 16 hours at 37°C. A 1-mL aliquot of culture was used for a miniprep with the QIAprep spin miniprep kit (Qiagen). For enhanced ssDNA capture, minipreps were modified by the addition of two volumes of 100% ethanol to the clarified lysate after N3 neutralization and centrifugation (e.g. 1.6 mL EtOH for 800 μL supernatant), before loading onto the silica columns (**fig. S37A**). Purified DNA concentration was quantified using a NanoDrop One (ThermoFisher) in dsDNA mode. For denaturing PAGE, 100 ng of each sample was mixed with Novex TBE-urea sample buffer (ThermoFisher), run on a Novex 10% TBE-Urea gel (ThermoFisher) at 250 volts, and stained with SYBR Gold (ThermoFisher).

### Strand-specific DNA sequencing

Strand-specific DNA sequencing was performed largely as previously described (*43*). 1.5 μg of input DNA was incubated with 20 U of terminal deoxynucleotidyl transferase (NEB) and 4 μM deoxycytidine triphosphate (dCTP) or deoxyadenosine triphosphate (dATP) in 1x reaction buffer (without CoCl2) at 37°C for 30 minutes, then 70°C for 10 minutes. Primer ssExt_pT9 or ssExt_pG6 anchor was then annealed to the DNA (95°C 1 minute, 4°C hold), followed by the addition of 1 mM dNTPs and 5 U Klenow fragment (exo-, NEB). The reaction was incubated at 37°C for 30 minutes then 65°C for 5 minutes and products were purified using the QIAquick PCR purification kit (Qiagen). 1 μM each of the adaptor primers (ssd_adapt_top and ssd_adapt_bottom) were annealed in 100 μL H2O with a gradual annealing of 95°C for 1 minute, and then a 10°C decrease every 10 minutes to 5°C. Annealed adaptor (50 nM, 1:20 dilution of the pre-annealed product) was ligated to the recovered DNA using Blunt/TA ligase master mix (NEB). 1 μL of each ligation reaction was amplified with NEBNext in a 50 μL reaction with barcoded primer pairs for 25 cycles. PCR products were pooled, 250–1000 bp products were excised from a 2% E-Gel (ThermoFisher), purified using a QIAquick gel extraction kit (Qiagen), pooled, and then repurified using a QIAquick PCR purification kit (Qiagen). Pooled libraries were run on a MiSeq i100 with 2 × 120 bp cycles.

### Sequencing analysis and cDNA product secondary structure prediction

For each sample, adaptor and homopolymer-tail sequences were removed with cutadapt (v5.2): the 3′ adaptor was trimmed from read 1 (TdT homopolymer tail plus single-strand extension anchor, {A|C}n·CTGTCTCTTATACACATCTCCGAGCCCACGAGAC) and from read 2 (CTGTCTCTTATACACATCTGACGCTGCCGACGA), with quality trimming (-q 20, --nextseq-trim=20, --trim-n) and a 15-nt minimum length. The leading TdT-tail homopolymer (poly-T for dATP tailing, poly-G for dCTP tailing) was then stripped from the read 2 5′ end with a custom script to expose the true 3′ terminus of each ssDNA molecule. Reads were mapped with bwa-mem to a combined reference comprising the *E. coli* BL21(DE3) host genome and the corresponding expression plasmid, and alignments were processed with samtools (v1.23.1; fixmate, coordinate sort, index). Per-strand coverage, 5′-end density, and 3′-end density were tabulated from properly paired reads (one count per pair) using pysam (v0.24.0) and normalized to counts per million properly paired read pairs (CPM) for cross-sample comparison. Coverage profiles were displayed on a symmetric-log axis (linear threshold = 1 CPM) to preserve zero-coverage positions. The ends of the cDNA products were determined by the extra A on the 5′ end due to Klenow and on the 3′ end by the start of the homopolymer tail (disambiguated by the differential tailing using both dATP and dCTP). The data shown are for dCTP-tailed products. cDNA product structures were predicted using ViennaRNA with DNA parameters.

