## Supplementary Figures for "Coevolutionary mining of prokaryotic non-coding elements with a genome language model"

### Supplementary Text

*Loss does not serve as a confidence measure*

We asked whether language-modeling loss could serve as a proxy for interaction-prediction performance and thereby provide a measure of prediction confidence. Across models, average language-modeling loss did not consistently predict interaction performance (**fig. S11D**). Additionally, per-sequence base-pairing performance was uncorrelated (r=-0.002, p=0.96) with per-sequence loss within a model (**fig. S11E**).

*TwoAYGGAY genome architecture*

Using a 2-kb distance cutoff to group neighboring elements, 70.9% occur as singletons, 26.2% as doublets, and only 2.8% as triplets or higher-order clusters (**Fig. 4F**). Triplets and higher-order clusters were not significantly enriched after accounting for the excess of doublets. The divergent bias of doublets noted in the main text is reminiscent of divergently oriented REPs that make up REPINs, a non-autonomous mobile unit (*50*). In doublets, the two elements typically carry unequal numbers of 5′ motif repeats rather than comparable arrays (**fig. S23**).

*Minerva-predicted TwoAYGGAY architectural variation in Caloramator australicus*

In *Caloramator australicus*, Minerva-MLM interaction heads predicted a distinct architecture in which the extended basal stem is flanked on both sides by repeated hairpins. The sequences of these flanking hairpins vary among genomic copies, resembling a direct-repeat organization of hairpin elements (**fig. S29**).

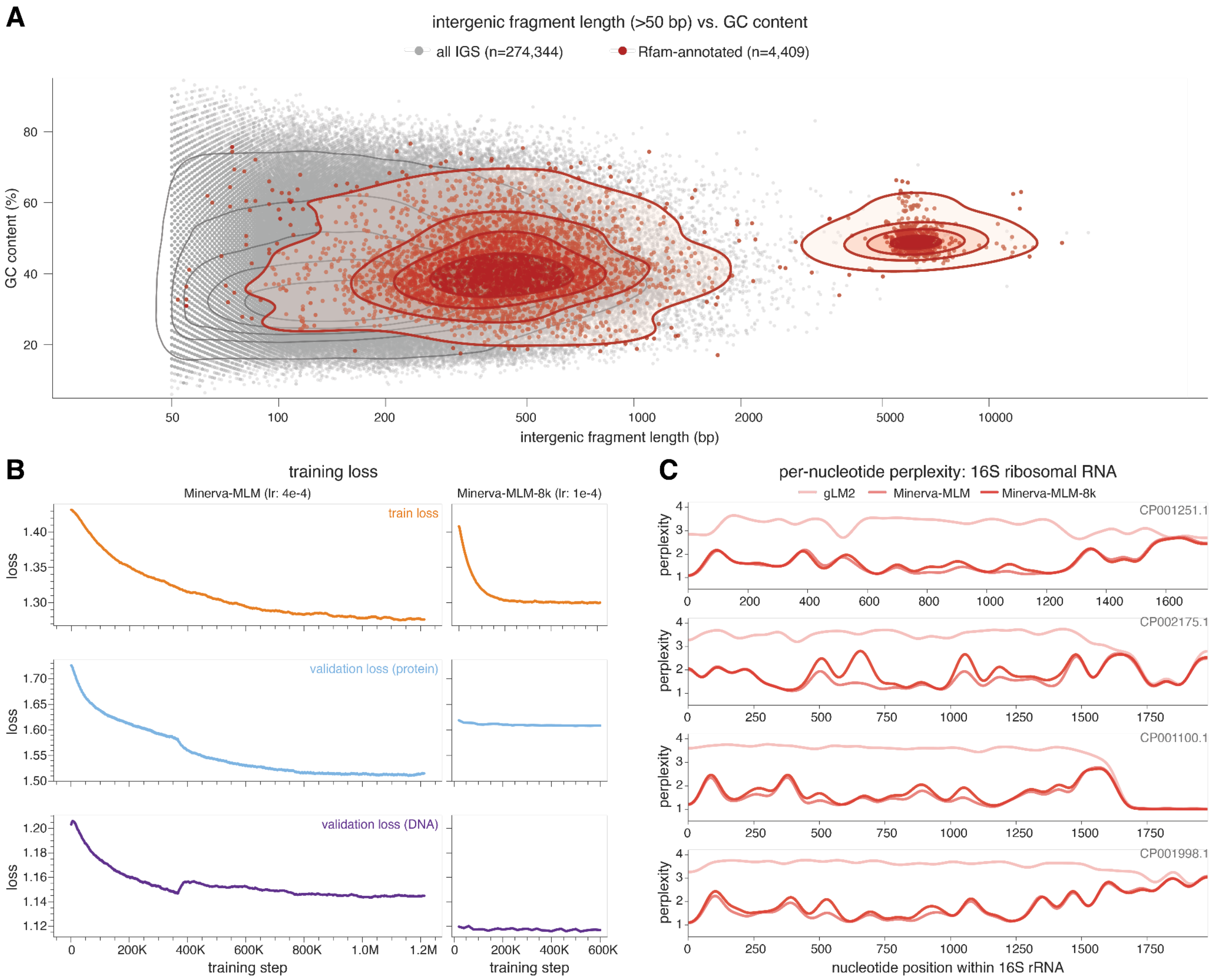


**Figure S1**: **Training the Minerva-MLM genome language model**

**(A)** Intergenic fragment length versus GC content for all intergenic regions greater than 50 base pairs in length from the 150 selected genomes in **table S1**, with regions annotated as matches to Rfam covariance models shown in red.

**(B)** Training loss and per-modality validation losses for Minerva-MLM and Minerva-MLM-8k.

**(C)** Per-nucleotide perplexity across 16S rRNAs from CP001251.1, CP002175.1, CP001100.1, and CP001998.1, compared across Minerva-MLM, Minerva-MLM-8k, and gLM2.

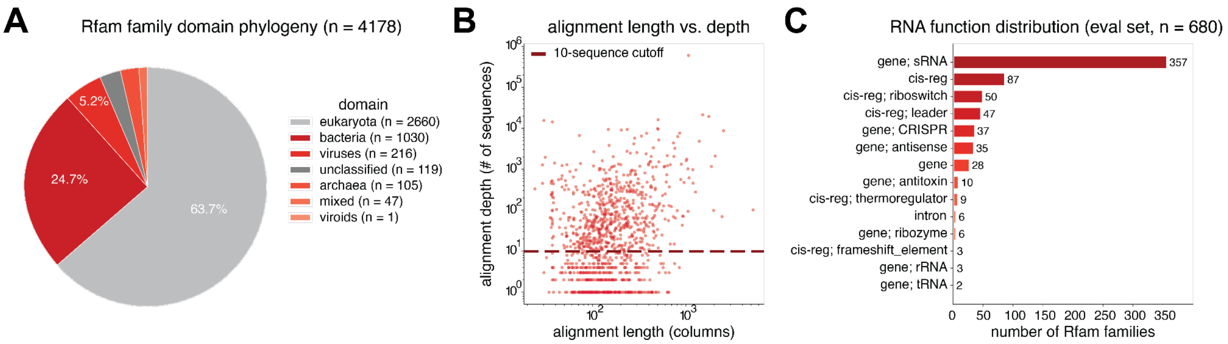


**Figure S2: Composition of the Rfam RNA base-pairing evaluation**

**(A)** Taxonomic distribution of the 4,178 Rfam families considered when constructing the base-pairing evaluation by primary domain of origin. Some unclassified and mixed families are retained in the evaluation as bacterial Rfam families, for a total of 1093 bacterial families.

**(B)** Seed alignment length versus alignment depth for each Rfam family. The dashed line indicates the minimum depth of 10 sequences required for families to be retained for evaluation.

**(C)** Rfam functional class distribution for the 680 families retained in the evaluation set.

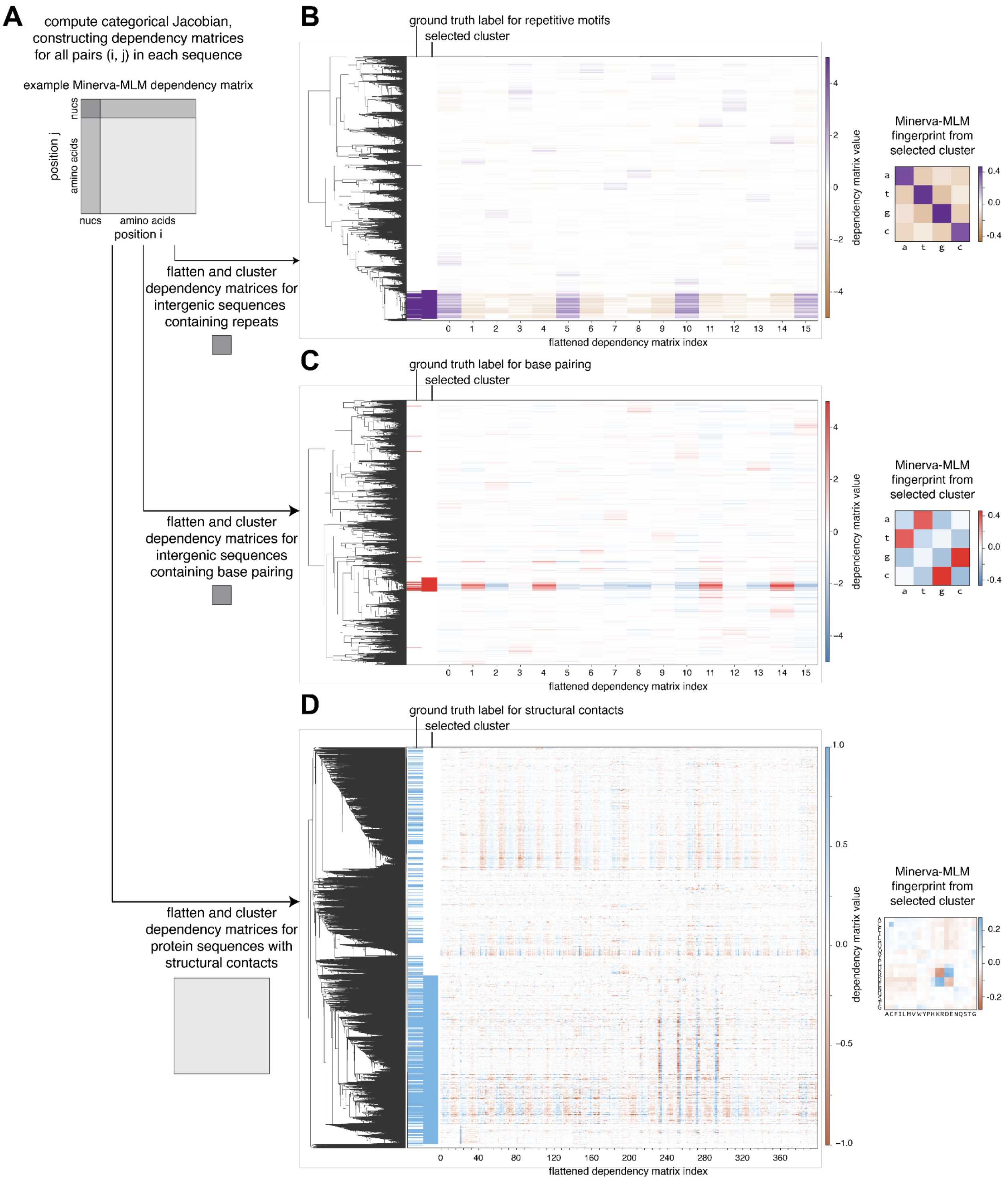


**Figure S3**: **Construction of reference fingerprints for categorical Jacobian fingerprinting using the Minerva-MLM model**

**(A)** Schematic of reference-fingerprint construction for Minerva-MLM categorical Jacobian fingerprinting. Dependency matrices computed for all position pairs in labeled benchmark examples are flattened and hierarchically clustered.
**(B)** Flattened Minerva-MLM dependency matrices for position pairs from the repetitive-motif evaluation. Matrices are hierarchically clustered without reference to their interaction labels, which are subsequently used to select the cluster defining the repetitive-motif fingerprint. The selected cluster and its mean nucleotide dependency matrix are shown.
**(C)** As in **(B)**, for base-paired nucleotide positions in the Rfam evaluation. The selected cluster and its mean nucleotide dependency matrix define the base-pairing fingerprint.
**(D)** As in **(B)**, for contacting amino-acid positions in the PDB protein monomer evaluation. The selected cluster and its mean amino-acid dependency matrix define the protein-contact fingerprint.

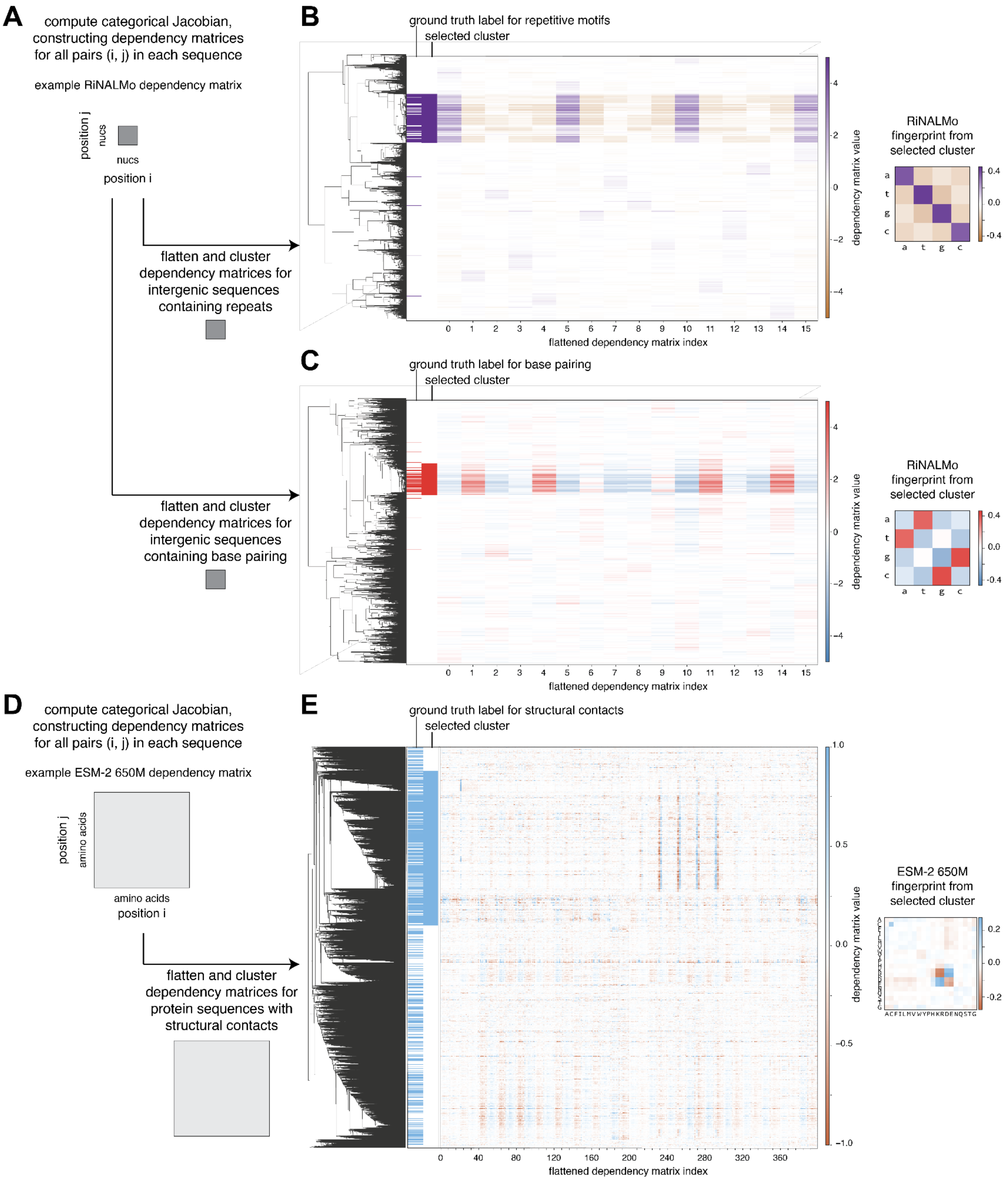


**Figure S4: Construction of reference fingerprints for categorical Jacobian fingerprinting using the RiNALMo and ESM-2 650M models**

**(A)** Schematic of reference-fingerprint construction for RiNALMo categorical Jacobian fingerprinting, as in **Figure S3A**.

**(B)** Flattened RiNALMo dependency matrices for position pairs from the repetitive-motif evaluation. Matrices are hierarchically clustered without reference to their interaction labels, which are subsequently used to select the cluster defining the RiNALMo repetitive-motif fingerprint. The selected cluster and its mean nucleotide dependency matrix are shown.
**(C)** As in **(B)**, for base-paired nucleotide positions in the Rfam evaluation. The selected cluster and its mean nucleotide dependency matrix define the RiNALMo base-pairing fingerprint.
**(D)** Schematic of reference-fingerprint construction for ESM-2 650M categorical Jacobian fingerprinting, as in **Figure S3A**.
**(E)** Flattened ESM-2 650M dependency matrices for residue pairs from the PDB protein monomer evaluation. Matrices are hierarchically clustered without reference to their interaction labels, which are subsequently used to select the cluster defining the ESM-2 650M protein-contact fingerprint. The selected cluster and its mean amino-acid dependency matrix are shown.

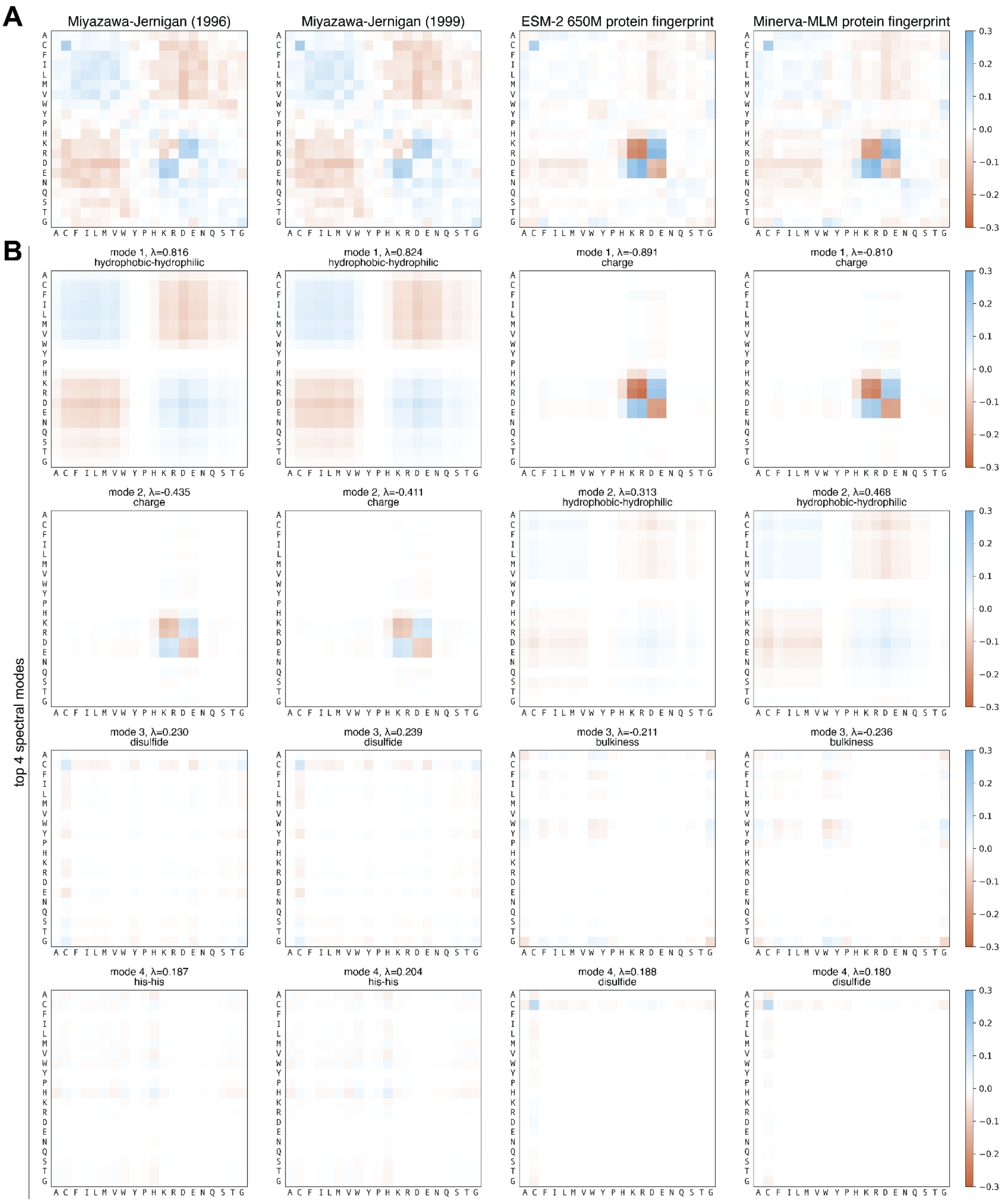


**Figure S5: Protein-contact reference fingerprints resemble the Miyazawa-Jernigan statistical potential**

**(A)** Comparison of the Miyazawa-Jernigan statistical contact potentials from 1996 and 1999 (*37*, *38*) with protein-contact reference fingerprints derived from ESM-2 650M and Minerva-MLM. Amino acids are ordered by physicochemical class.
**(B)** The four leading spectral modes of each matrix in **(A)**. Eigenvalues and the dominant physicochemical property associated with each mode are indicated above the corresponding plot.

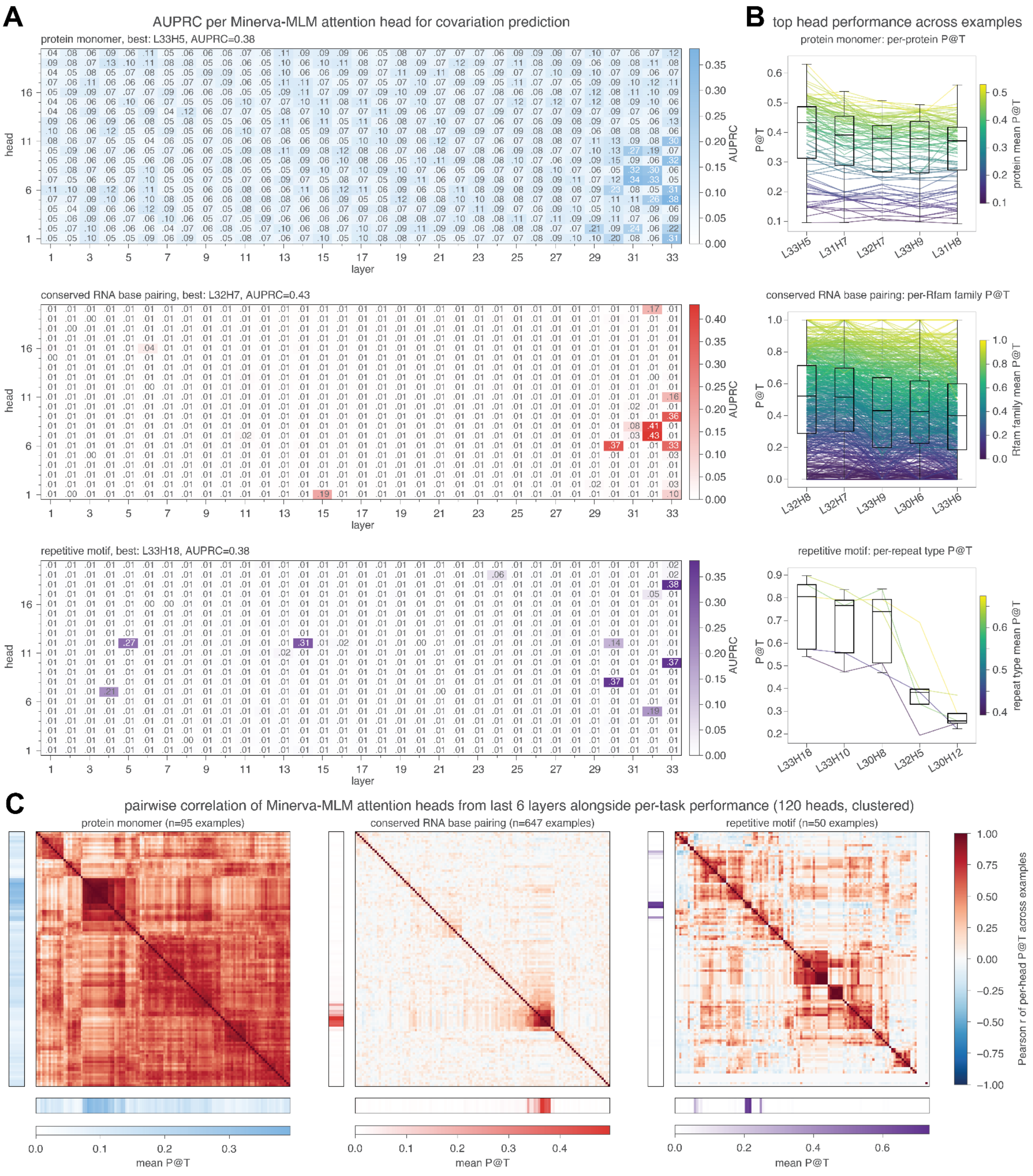


**Figure S6: Individual Minerva-MLM attention heads encode distinct coevolutionary signals**

**(A)** Covariation-prediction AUPRC for every Minerva-MLM attention head, arranged by layer and head index, for the protein monomer-contact (L33H5 best, AUPRC = 0.38), conserved RNA base-pairing (L32H7 best, AUPRC = 0.43), and repetitive motif (L33H18 best, AUPRC = 0.38) evaluations.

**(B)** Per-example precision among the top T highest predictions (P@T, where T is the total number of true interactions) for the five best-performing heads on each evaluation. Boxplots summarize the distribution across examples; connecting lines follow individual examples across heads and are colored by that example's mean P@T.
**(C)** Pairwise Pearson correlation of per-example P@T among the 120 attention heads in the last six layers, hierarchically clustered separately for each evaluation. Protein monomers, n = 95; conserved RNA base pairing, n = 647; repetitive motif, n = 50. Marginal bars show mean P@T for each attention head.

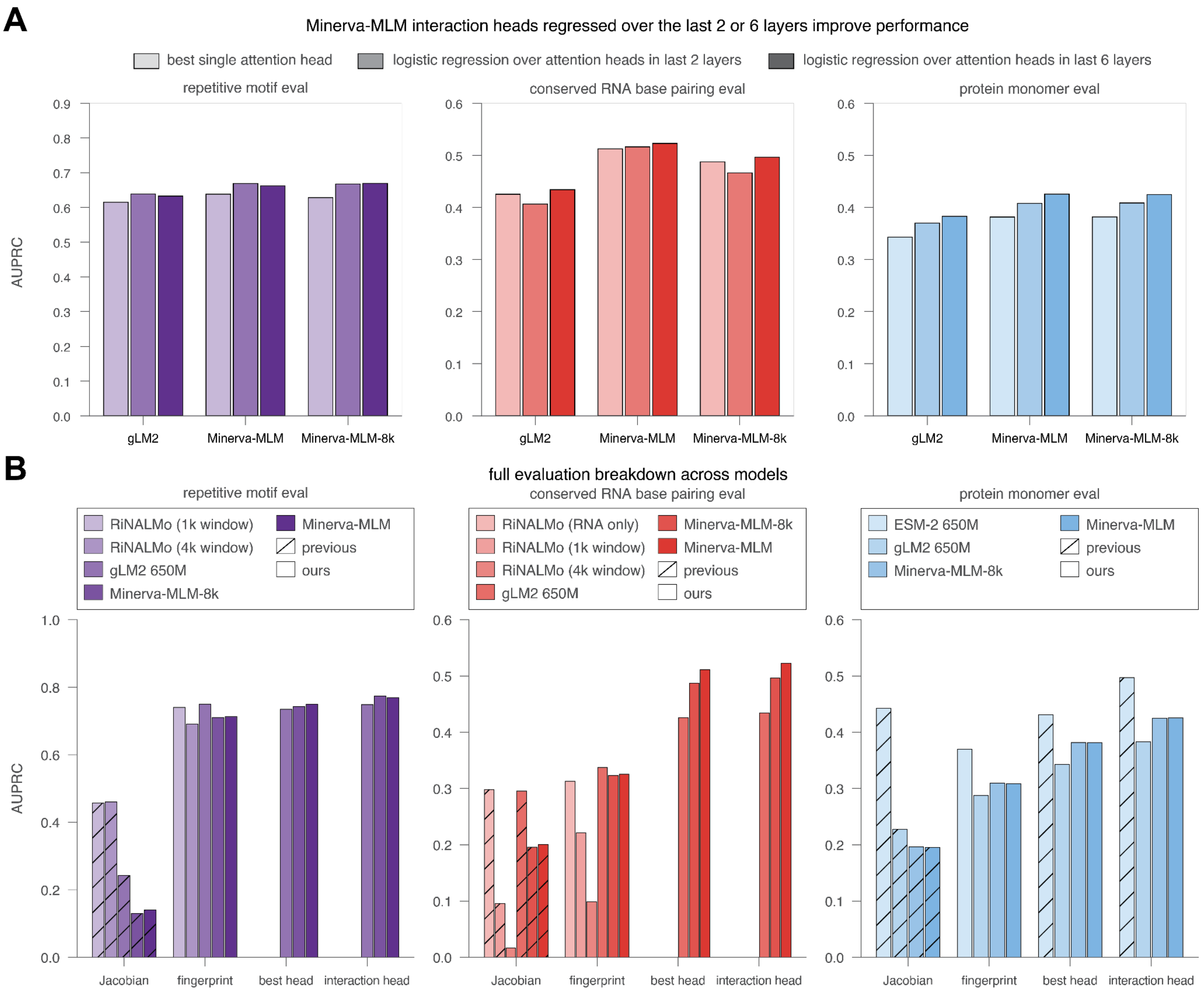


**Figure S7: Performance across models and interaction-extraction methods**
**(A)** Performance as measured by AUPRC of the best individual attention head and logistic-regression interaction heads combining attention maps from the final two or final six layers of gLM2, Minerva-MLM, and Minerva-MLM-8k. Results are shown for repetitive-motif, conserved RNA base-pairing, and protein monomer-contact evaluations.
**(B)** Full performance comparison as measured by AUPRC of categorical Jacobian, categorical Jacobian fingerprinting, the best individual attention head, and last-two-layer interaction heads across the three evaluation sets and across models. Colors denote the underlying language model, and hatching distinguishes previously described methods from methods introduced here.

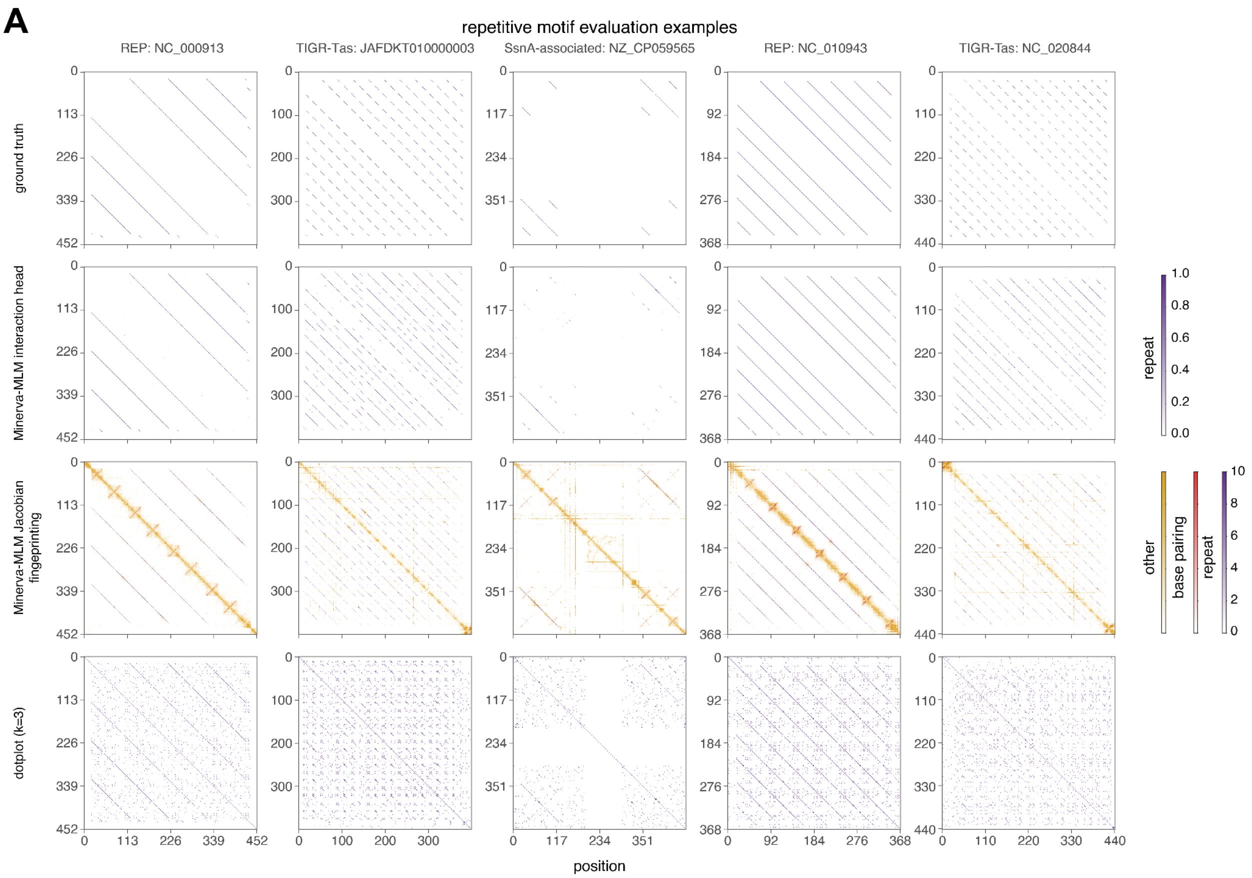


**Figure S8: Repetitive-motif evaluation examples**

**(A)** Five examples from the repetitive-motif evaluation: REP loci from NC_000913 and NC_010943, TIGR-Tas loci from JAFDKT010000003 and NC_020844, and an SsnA-associated repeat locus from NZ_CP059565. For each locus, rows show the labeled reference repetitive-motif interactions, predictions from the Minerva-MLM interaction head, predictions from Minerva-MLM categorical Jacobian fingerprinting, and a sequence dotplot (k = 3, forward matches) for comparison.

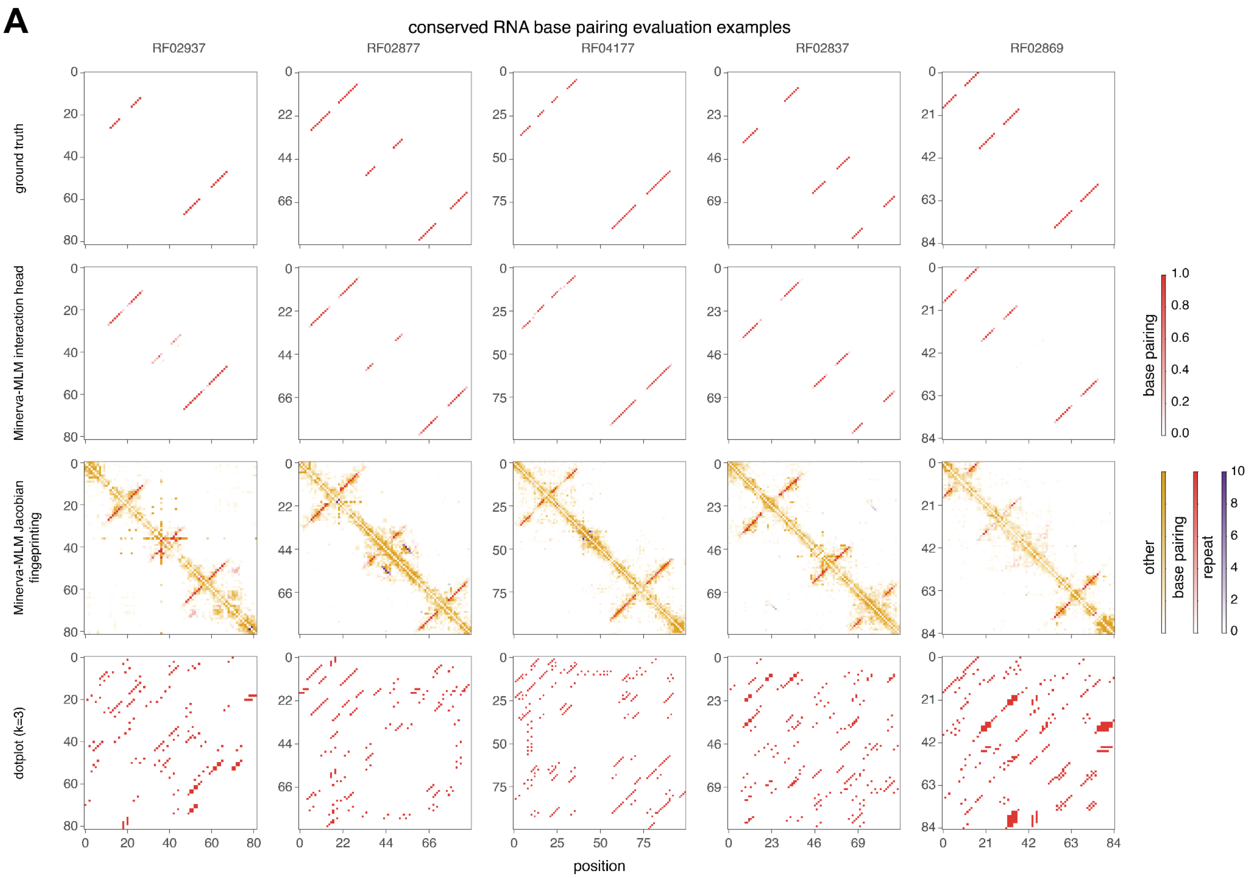


**Figure S9: Conserved RNA base-pairing evaluation examples**

**(A)** Five examples from the Rfam conserved RNA base-pairing evaluation: Rfam families RF02937, RF02877, RF04177, RF02837, and RF02869. For each locus, rows show the labeled reference base-pairing interactions, predictions from the Minerva-MLM interaction head, predictions from Minerva-MLM categorical Jacobian fingerprinting, and a sequence dotplot (k = 3, reverse-complement matches) for comparison.

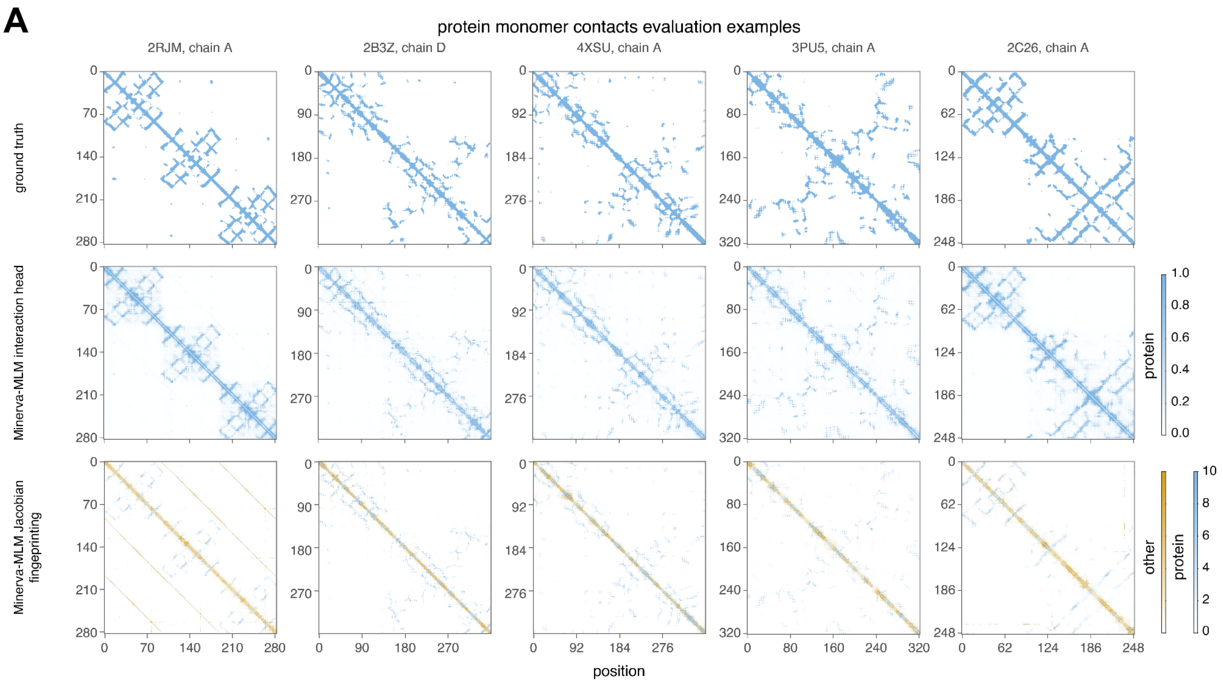


**Figure S10: Protein monomer contact evaluation examples**

**(A)** Five examples from the PDB protein monomer-contact evaluation: 2RJM chain A, 2B3Z chain D, 4XSU chain A, 3PU5 chain A, and 2C26 chain A. For each protein, rows show the labeled reference structural contacts, predictions from the Minerva-MLM interaction head, and predictions from Minerva-MLM categorical Jacobian fingerprinting.

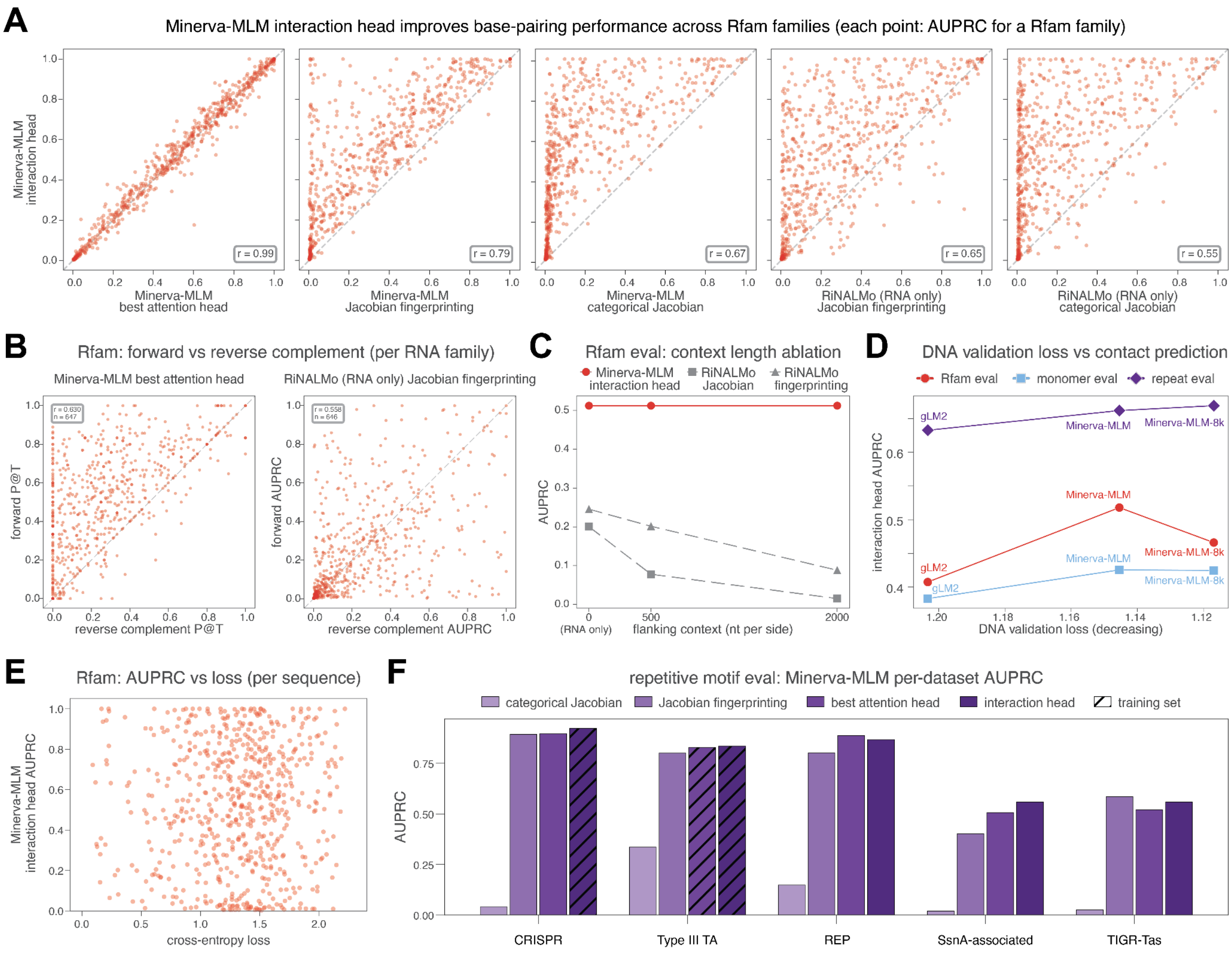


**Figure S11: Evaluation of interaction prediction across models and sequence contexts**
**(A)** Per-family Rfam base-pairing performance of the Minerva-MLM interaction head compared with the Minerva-MLM best individual attention head, Minerva-MLM categorical Jacobian fingerprinting, Minerva-MLM categorical Jacobian, RiNALMo categorical Jacobian fingerprinting on isolated RNAs, and RiNALMo categorical Jacobian scoring on isolated RNAs. Each point represents one Rfam family and reports AUPRC.
**(B)** Base-pairing performance on forward-oriented versus reverse-complemented Rfam sequences for the Minerva-MLM best attention head and RiNALMo categorical Jacobian fingerprinting. Each point represents one RNA family.
**(C)** Effect of genomic context on Rfam base-pairing evaluation performance for the Minerva-MLM interaction head, RiNALMo categorical Jacobian, and RiNALMo categorical Jacobian fingerprinting. RNAs are evaluated in isolation or with increasing flanking genomic context on each side.
**(D)** DNA-validation loss of gLM2, Minerva-MLM, and Minerva-MLM-8k compared with their performance as measured by AUPRC on the Rfam base-pairing, protein monomer-contact, and repetitive-motif evaluations.

**(E)** Per-sequence Minerva-MLM interaction-head AUPRC on the Rfam evaluation compared with language modeling cross-entropy loss for the same sequence.
**(F)** Performance of Minerva-MLM categorical Jacobian, categorical Jacobian fingerprinting, the best attention head, and the interaction head as measured by AUPRC on the repetitive-motif evaluation split across the five different repetitive motif types. Hatched bars denote datasets used to fit the interaction head or select reference fingerprints.

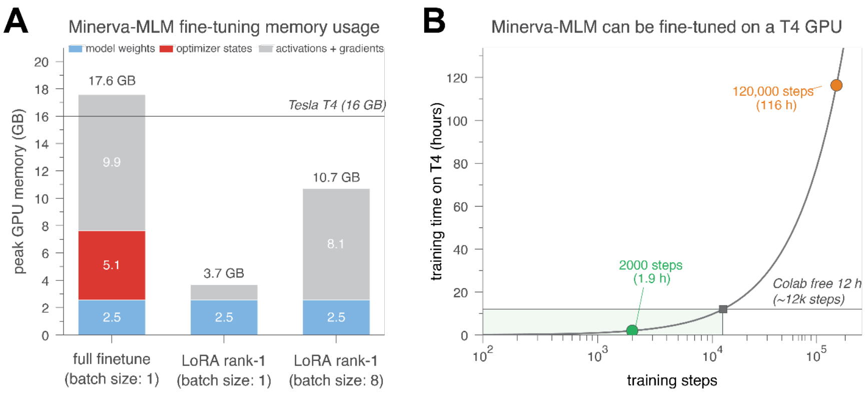


**Figure S12: Compute and memory requirements for fine-tuning Minerva-MLM**
**(A)** Peak GPU-memory use during Minerva-MLM fine-tuning with full-parameter fine-tuning at batch size 1 or rank-1 low-rank adaptation (LoRA) at batch sizes 1 and 8. Memory use is decomposed into model weights, optimizer states, and activations plus gradients. The horizontal line indicates the 16-GB capacity of an NVIDIA Tesla T4 GPU.
**(B)** Minerva-MLM fine-tuning time as a function of training steps estimated on a Tesla T4 GPU. Times for 2,000 and 120,000 steps are marked, together with the approximate number of steps possible within a 12-hour free Google Colab session.

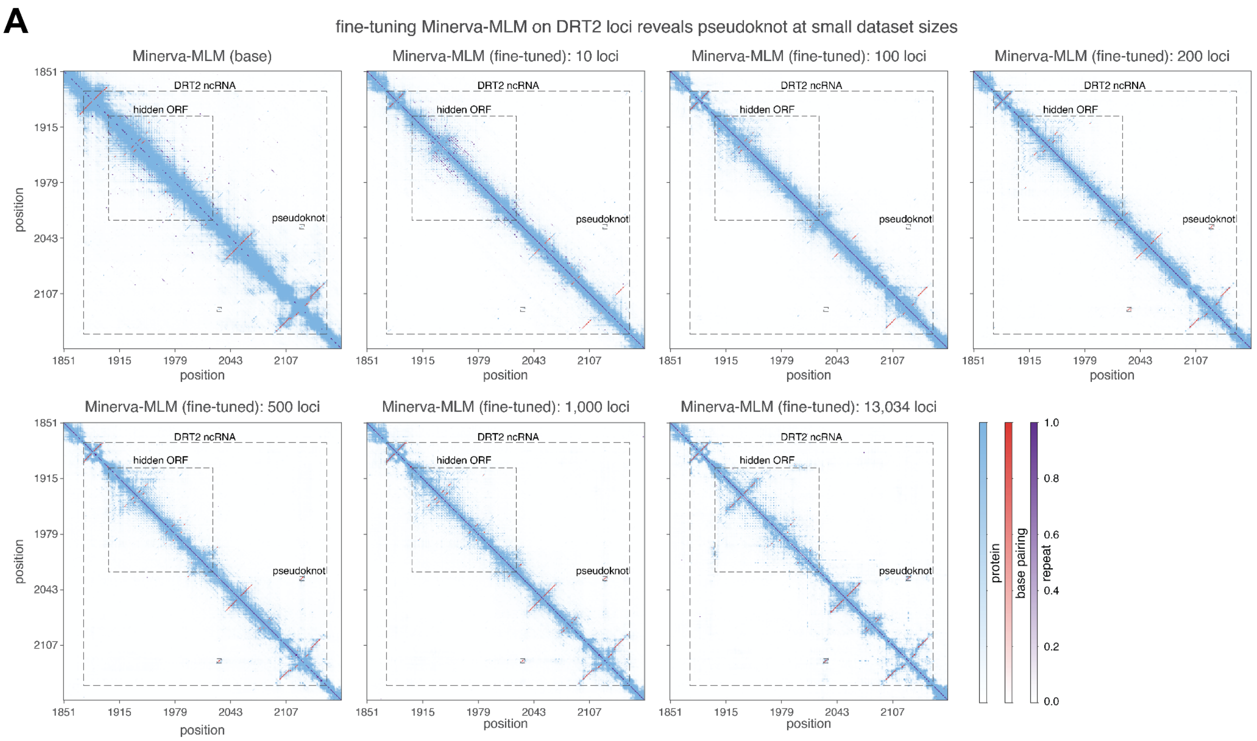


**Figure S13: Fine-tuning Minerva-MLM on small DRT2 datasets recovers the ncRNA pseudoknot**

**(A)** Minerva-MLM interaction-head predictions across the ncRNA and hidden ORF of the *Klebsiella pneumoniae* NTUH-K2044 DRT2 system before fine-tuning and after full-parameter fine-tuning on 10, 100, 200, 500, 1,000, or 13,034 DRT2 loci. The ncRNA, hidden ORF, and pseudoknot are annotated.

A

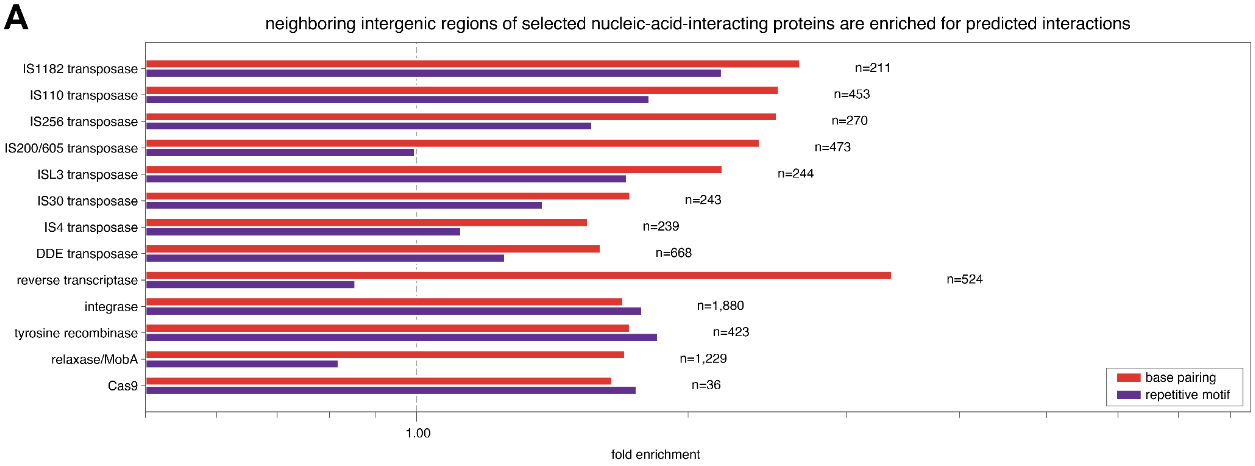


**Figure S14: Intergenic regions neighboring nucleic-acid-interacting proteins are enriched for predicted interactions**

**(A)** Fold enrichment of predicted base-pairing and repetitive-motif interactions in intergenic regions neighboring selected nucleic-acid-interacting protein classes, relative to all intergenic regions across the 150 genomes. The dashed line indicates no enrichment; n indicates the number of occurrences for each protein class.

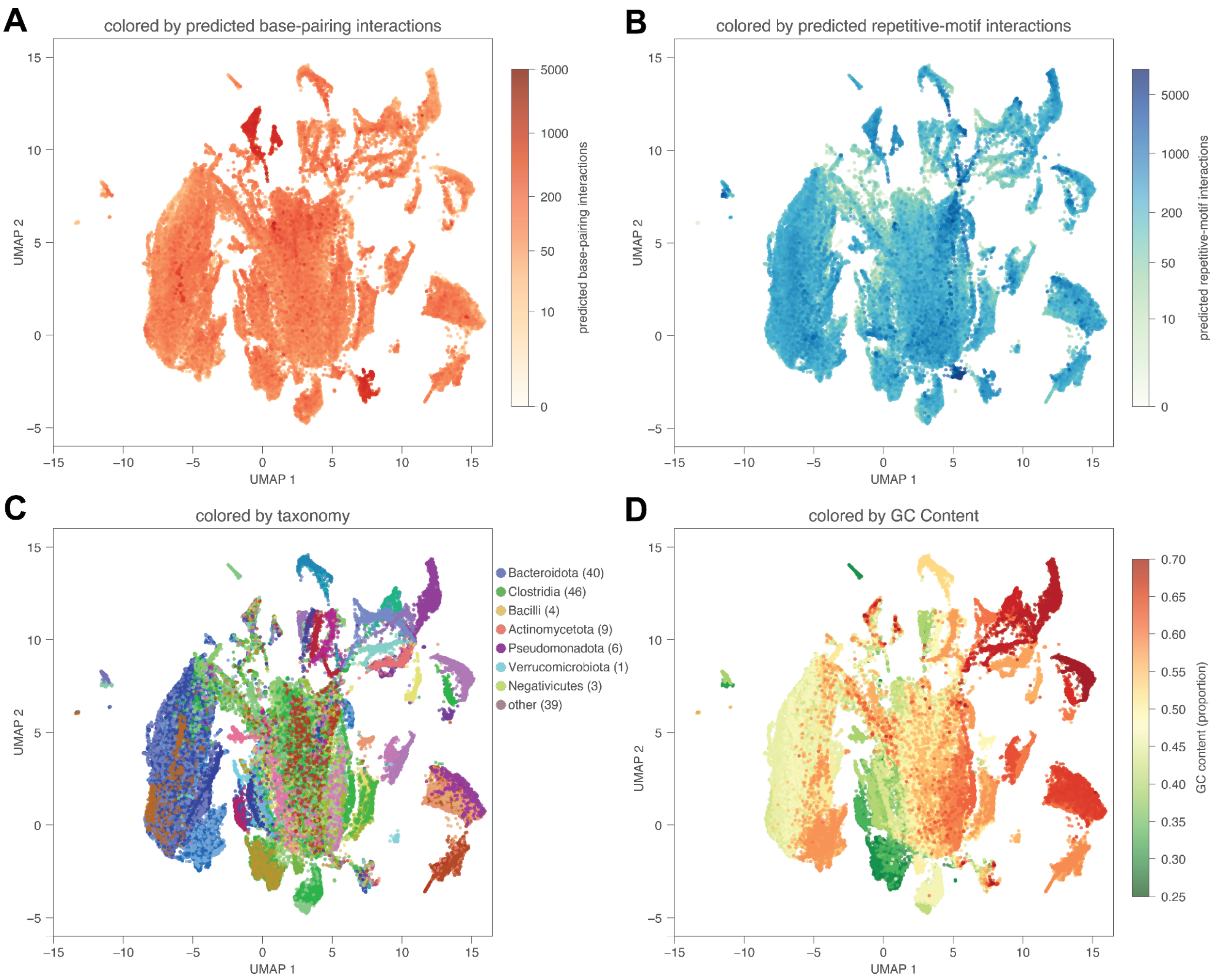


**Figure S15: Minerva-MLM embeddings reflect predicted structure, taxonomy, and sequence composition**

**(A–D)** UMAP of Minerva-MLM embeddings from the top 10% most structured intergenic regions across the 150 genomes as predicted by Minerva-MLM interaction heads, colored by **(A)** the number of interaction-head-predicted base-pairing interactions, **(B)** the number of interaction-head-predicted repetitive-motif interactions, **(C)** the taxonomy of the source genome, or **(D)** GC content. Compare with **Figure 3C**.

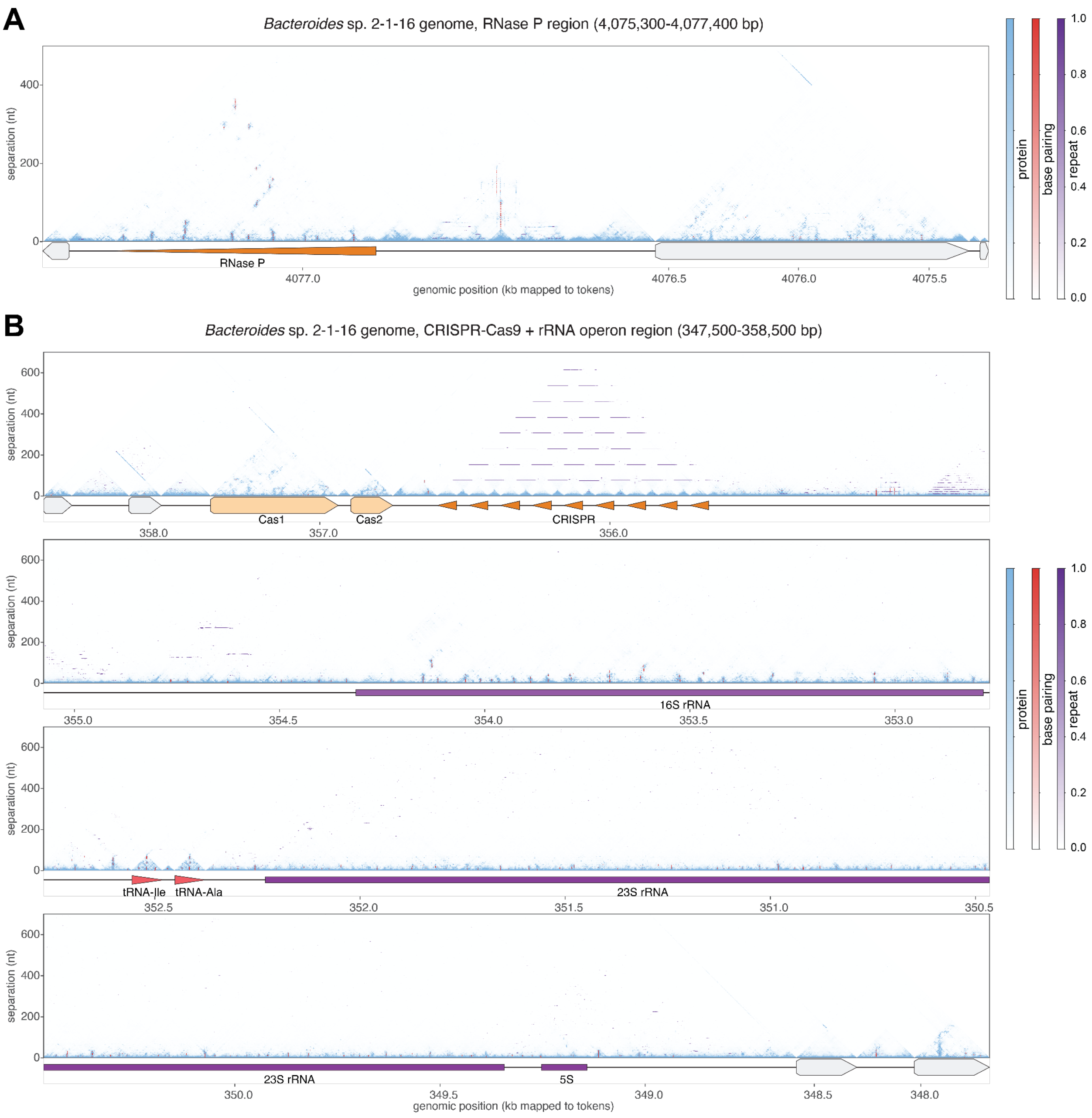


**Figure S16: Genome-scale coevolutionary maps at structured loci in *Bacteroides* sp. 2-1-16**
**(A)** Minerva-MLM interaction-head predictions across an RNase P-containing region (4,075,300–4,077,400 bp) in the *Bacteroides* sp. 2-1-16 genome. Pairwise predictions are plotted by genomic position and nucleotide separation, with the RNase P annotation shown below. Compare with **Figure 3D**.
**(B)** Minerva-MLM interaction-head predictions across a CRISPR-Cas9 locus and neighboring rRNA operon (347,500–358,500 bp) in the *Bacteroides* sp. 2-1-16 genome. Pairwise predictions are plotted by genomic position and nucleotide separation; Cas genes, the CRISPR array, 16S, 23S, and 5S rRNAs, and neighboring tRNAs are annotated below. Compare with **Figure 3D**.

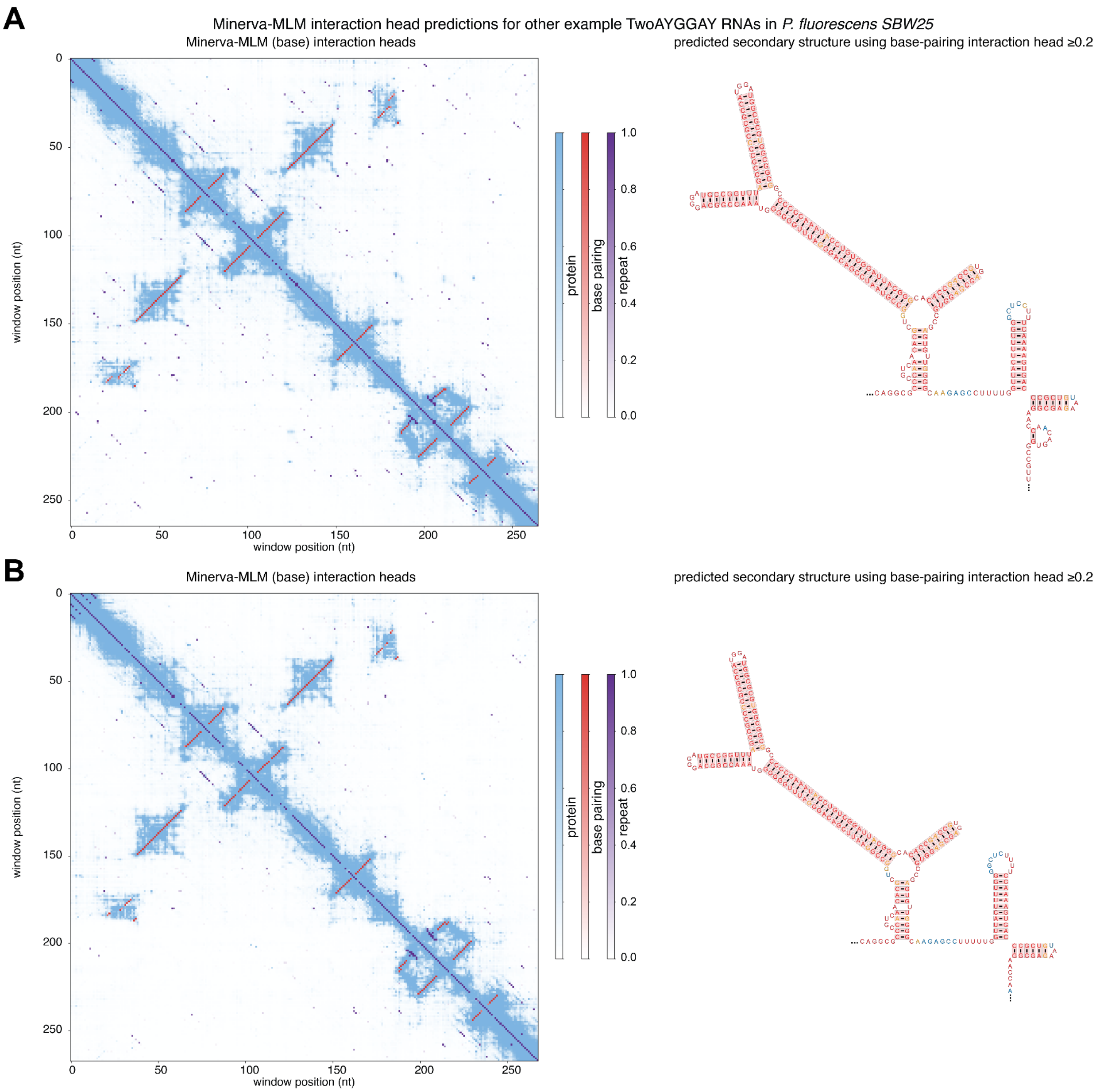


**Figure S17: Additional examples of extended TwoAYGGAY RNA structures in *Pseudomonas fluorescens* SBW25**

**(A–B)** Minerva-MLM base-model interaction-head predictions for two additional *Pseudomonas fluorescens* SBW25 TwoAYGGAY RNAs. Predicted secondary structures generated from base-pairing interaction-head scores of at least 0.2 are shown to the right. Positions predicted to be involved in base-pairing interactions inconsistent with Watson-Crick base-pairing are colored orange. Positions predicted to be involved in a pseudoknot consistent with Watson-Crick base-pairing are colored blue.

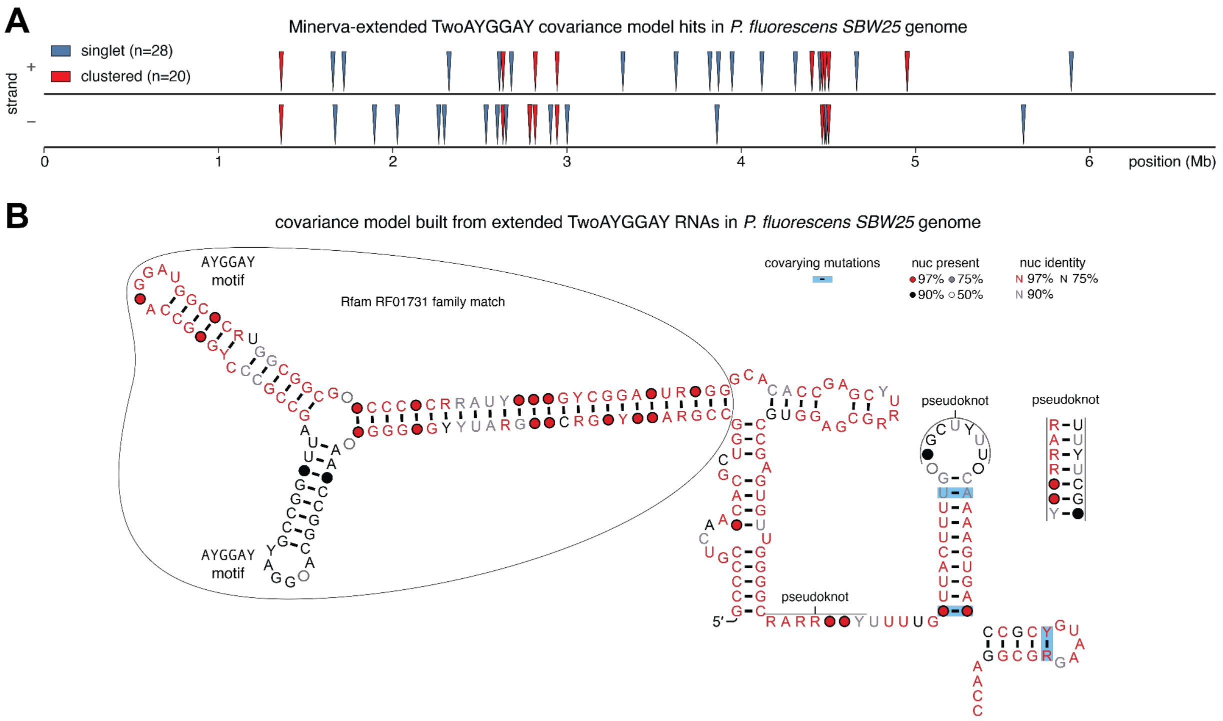


**Figure S18: Extended TwoAYGGAY RNAs in *Pseudomonas fluorescens* SBW25**
**(A)** Genomic positions and orientations of the 48 matches to the Minerva-extended TwoAYGGAY covariance model in the *Pseudomonas fluorescens* SBW25 genome. Matches are classified as singletons or members of local clusters defined using a 2 kb distance cutoff.
**(B)** Covariance model constructed from 41 non-truncated extended TwoAYGGAY RNAs in SBW25. The original Rfam RF01731 match, AYGGAY motifs, extended stems, and predicted pseudoknot are annotated. Nucleotide identity, occupancy, and covarying substitutions are indicated by the accompanying key.

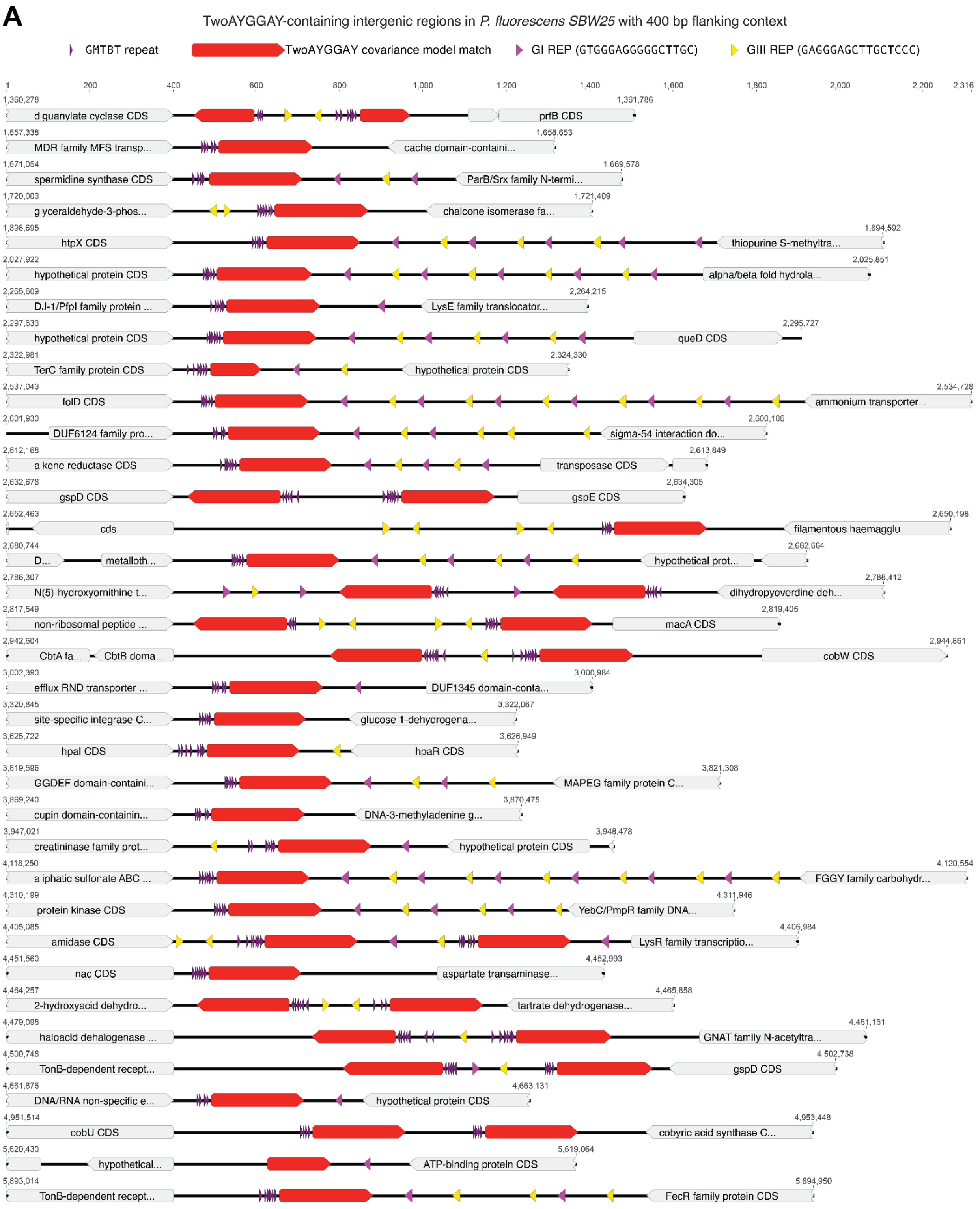


**Figure S19: Genomic contexts of TwoAYGGAY RNAs in *Pseudomonas fluorescens* SBW25**
**(A)** Intergenic regions containing matches to the extended TwoAYGGAY covariance model in SBW25, shown with 400 bp of flanking genomic context. TwoAYGGAY matches, upstream

GMTBT motifs, downstream group I and group III repetitive extragenic palindromic elements (REPs), and neighboring coding sequences are indicated.

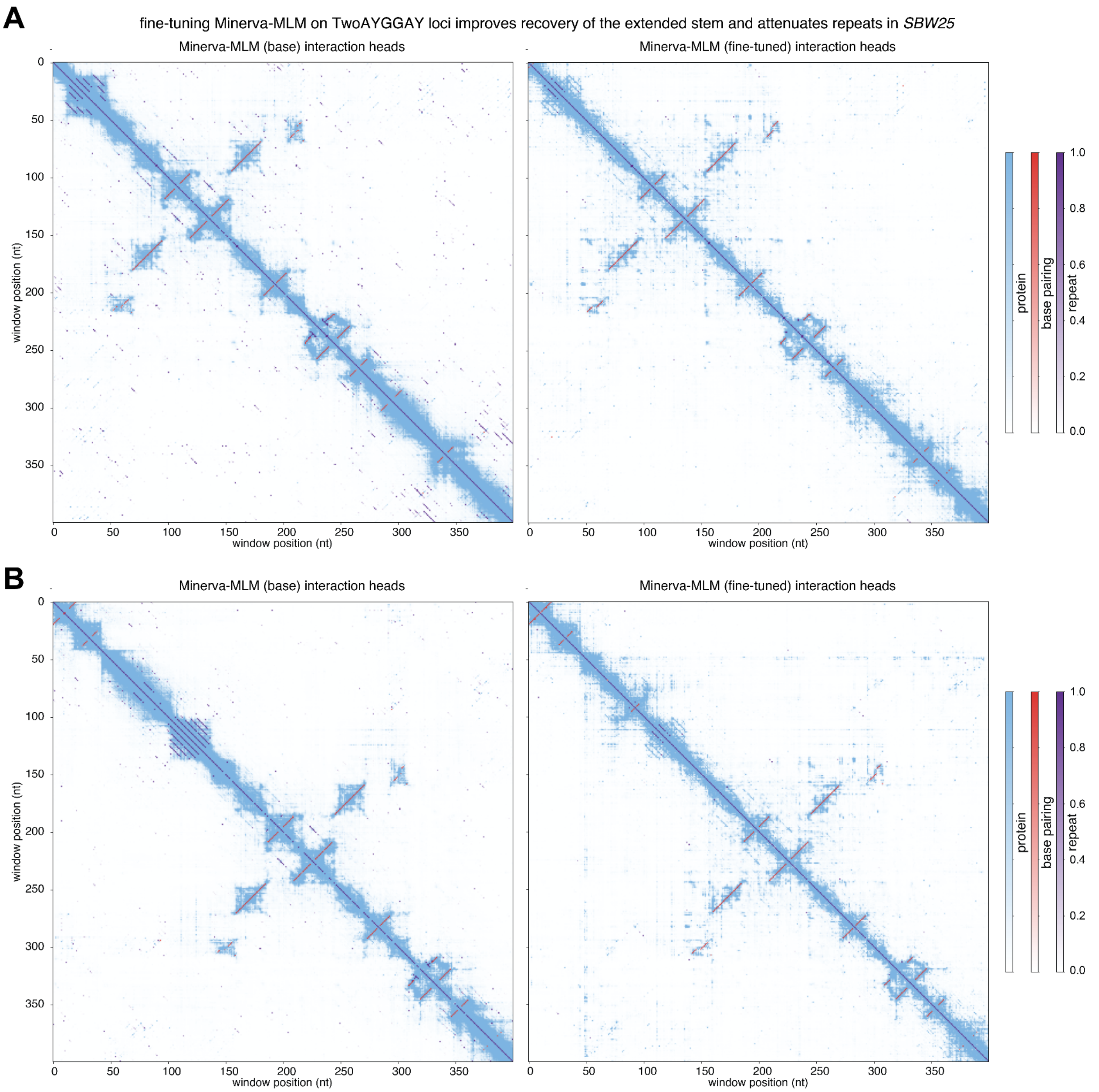


**Figure S20: Fine-tuning Minerva-MLM on TwoAYGGAY loci strengthens the extended TwoAYGGAY stem prediction and attenuates repeat signal**

**(A–B)** Minerva-MLM interaction-head predictions for two representative SBW25 TwoAYGGAY loci before and after family-specific full-parameter fine-tuning.

A

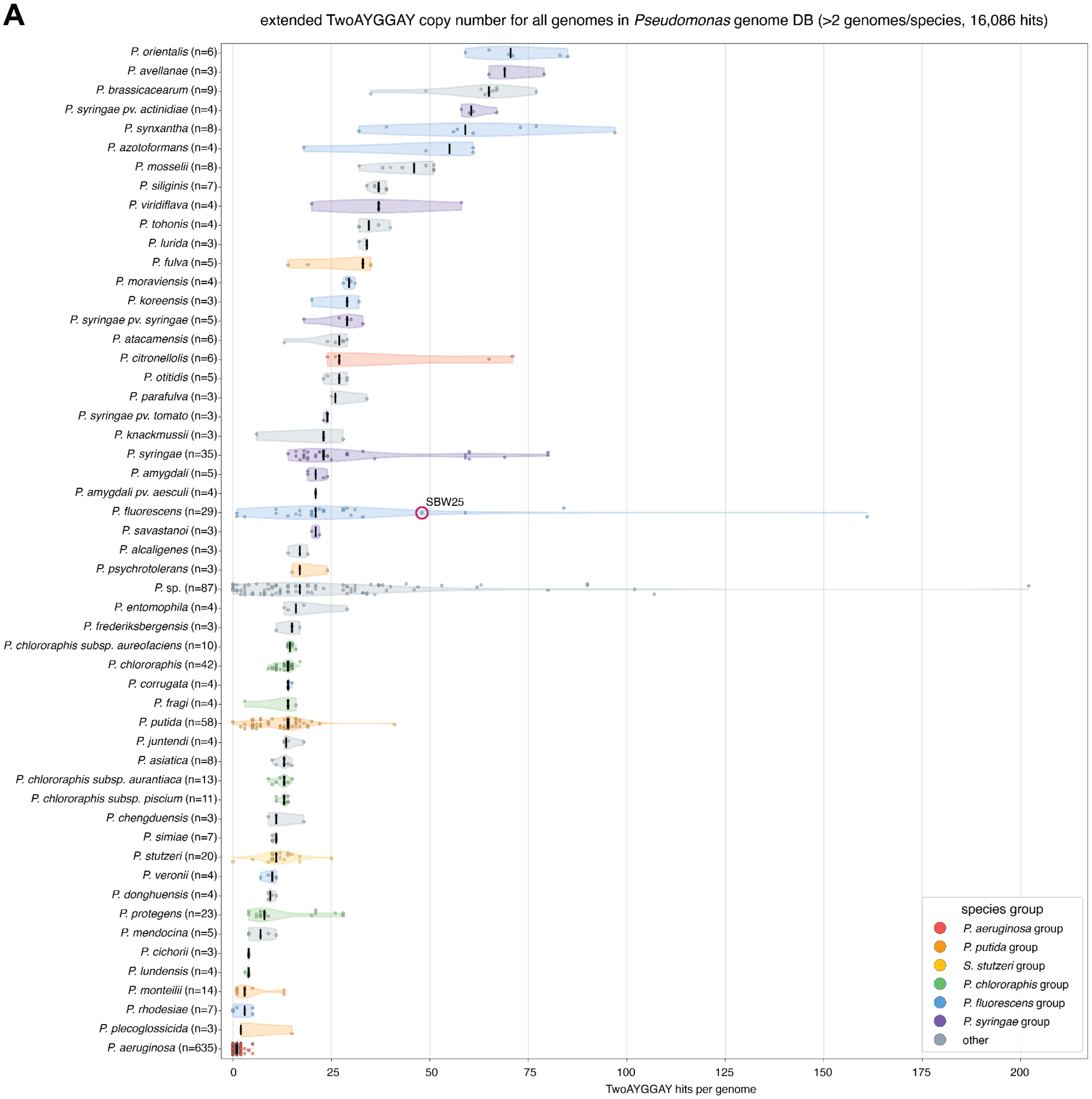


**Figure S21: TwoAYGGAY copy-number variation across *Pseudomonas* species**
**(A)** Number of Minerva-extended TwoAYGGAY covariance-model matches per complete genome for species represented by at least three genomes in the *Pseudomonas* Genome Database. Each point represents one genome; species are grouped by established *Pseudomonas* species group. The SBW25 genome is marked.

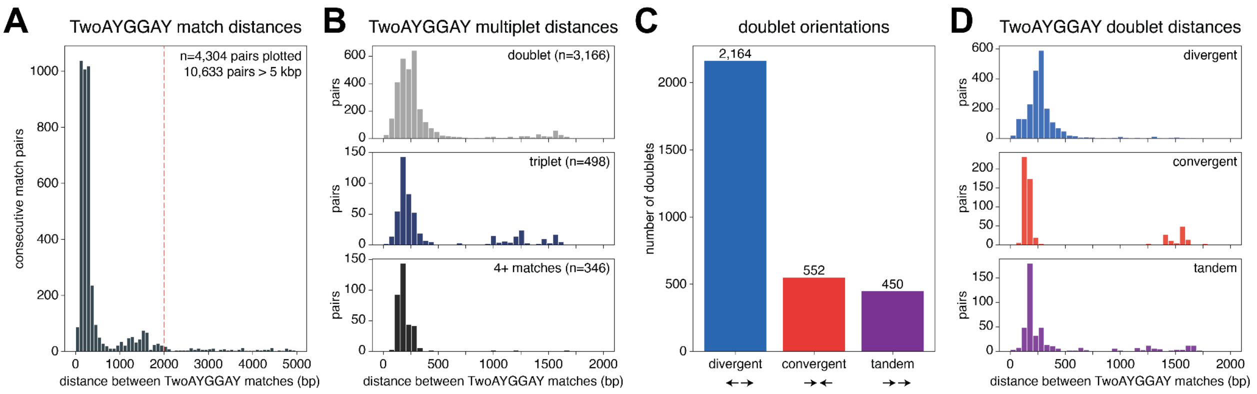


**Figure S22: Local organization and orientation of TwoAYGGAY elements**

**(A)** Distribution of distances between consecutive TwoAYGGAY matches in the *Pseudomonas* Genome Database. Pairs separated by at most 5 kb are shown; the numbers of plotted pairs and pairs separated by more than 5 kb are indicated. Dashed line marks the 2 kb clustering cutoff.

**(B)** Distributions of pairwise distances within loci containing two, three, or at least four neighboring TwoAYGGAY matches, with a cutoff of 2 kb between matches.

**(C)** Counts of divergent, convergent, and tandem TwoAYGGAY doublets.

**(D)** Distance distributions between TwoAYGGAY matches for divergent, convergent, and tandem doublets.

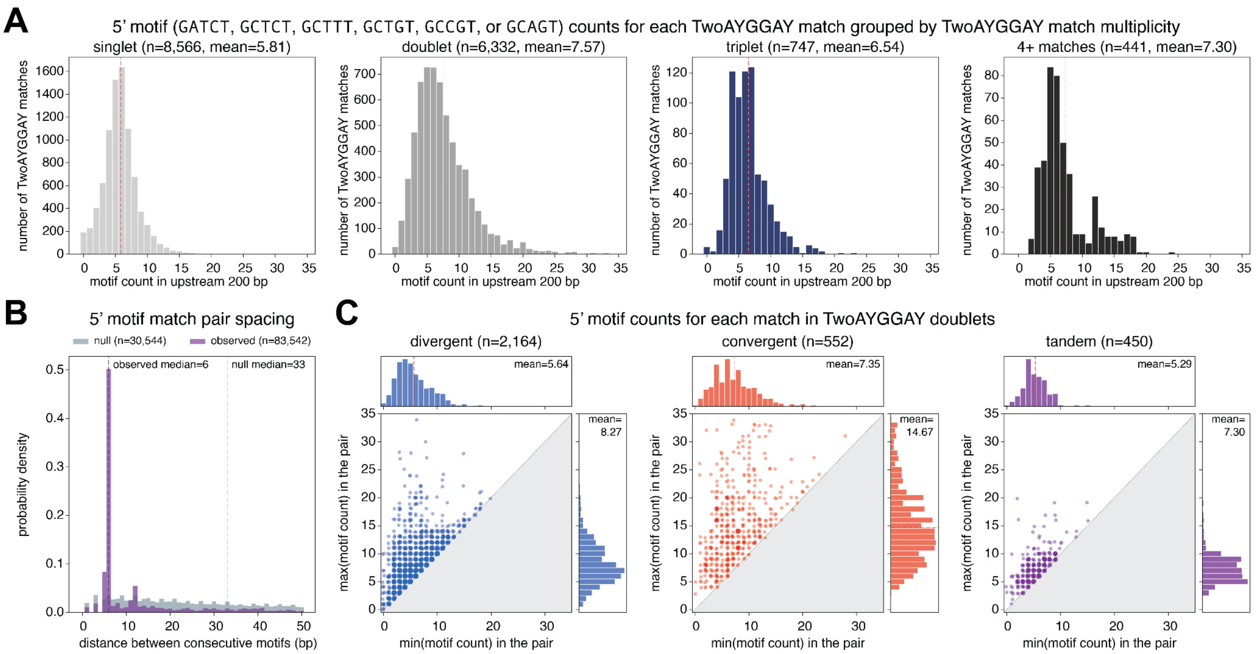


**Figure S23: Organization of upstream short motifs at TwoAYGGAY loci**

**(A)** Distributions of counts of the 5′ motif sequence (GATCT, GCTCT, GCTTT, GCTGT, GCCGT, or GCAGT) within the 200 bp upstream of individual TwoAYGGAY matches, grouped by whether the match occurs as a singleton, doublet, triplet, or higher-order cluster. Sample sizes and mean motif counts are shown.

**(B)** Spacing between consecutive upstream 5′ motif matches compared with spacing in randomly sampled null regions. Sample sizes and median spacings are shown. Related to **Figure 4H**.

**(C)** Numbers of upstream 5′ motifs associated with each member of divergent, convergent, and tandem TwoAYGGAY doublets. For each pair, the smaller motif count is plotted against the larger motif count; sample sizes and mean values are shown.

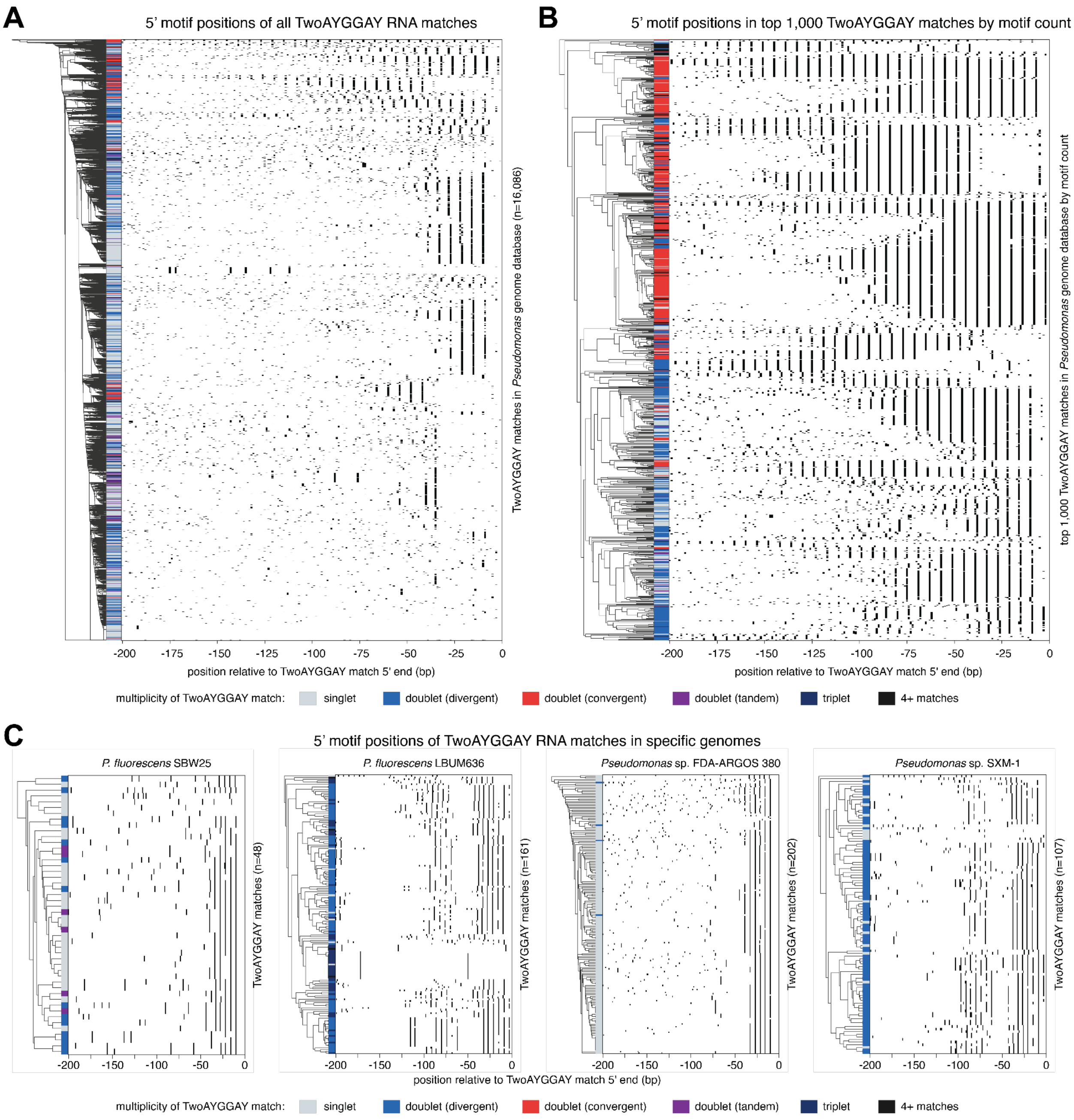


**Figure S24: Positioning of upstream short motifs relative to TwoAYGGAY RNAs is variable**

**(A)** Positions of 5′ motif sequence matches (GATCT, GCTCT, GCTTT, GCTGT, GCCGT, or GCAGT) within 200 bp upstream of all 16,086 extended TwoAYGGAY matches in the *Pseudomonas* Genome Database, as defined in **Figure S23**. Rows are hierarchically clustered and colored by local TwoAYGGAY multiplicity and arrangement.

**(B)** As in **(A)**, for the 1,000 TwoAYGGAY matches with the largest numbers of upstream motifs.

**(C)** Upstream motif positions for TwoAYGGAY matches in *P. fluorescens* SBW25, *P. fluorescens* LBUM636, *Pseudomonas* sp. FDA-ARGOS 380, and *Pseudomonas* sp. SXM-1. Rows are clustered and colored as in **(A)**.

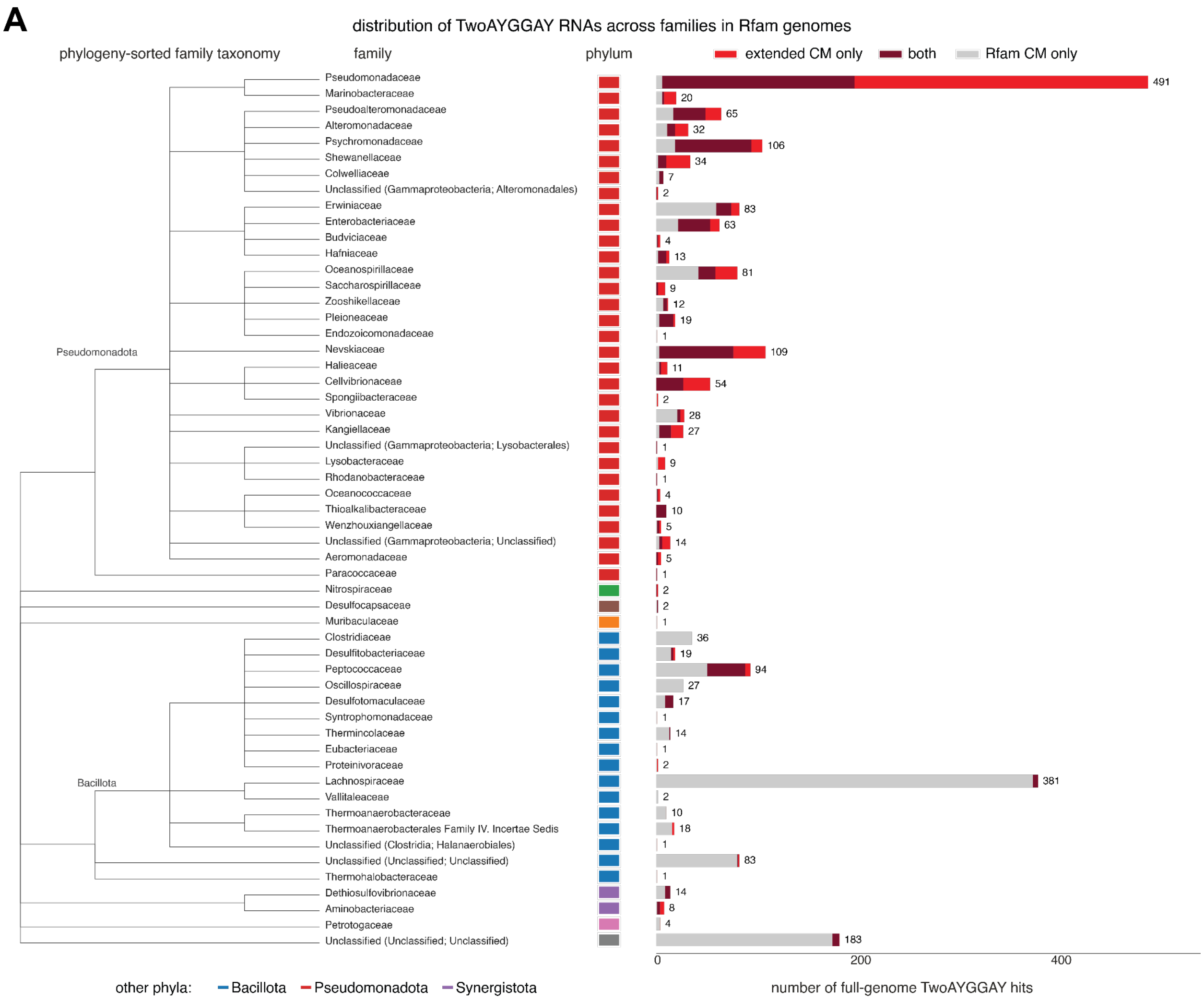


**Figure S25: Taxonomic distribution of extended TwoAYGGAY RNAs**

**(A)** Full-genome TwoAYGGAY covariance-model matches across taxonomic families represented in the Rfam genome collection. Families are ordered by taxonomy and classified by whether matches were detected by the Minerva-extended covariance model only, the original Rfam covariance model only, or both. Phylum is indicated by the adjacent color strip.

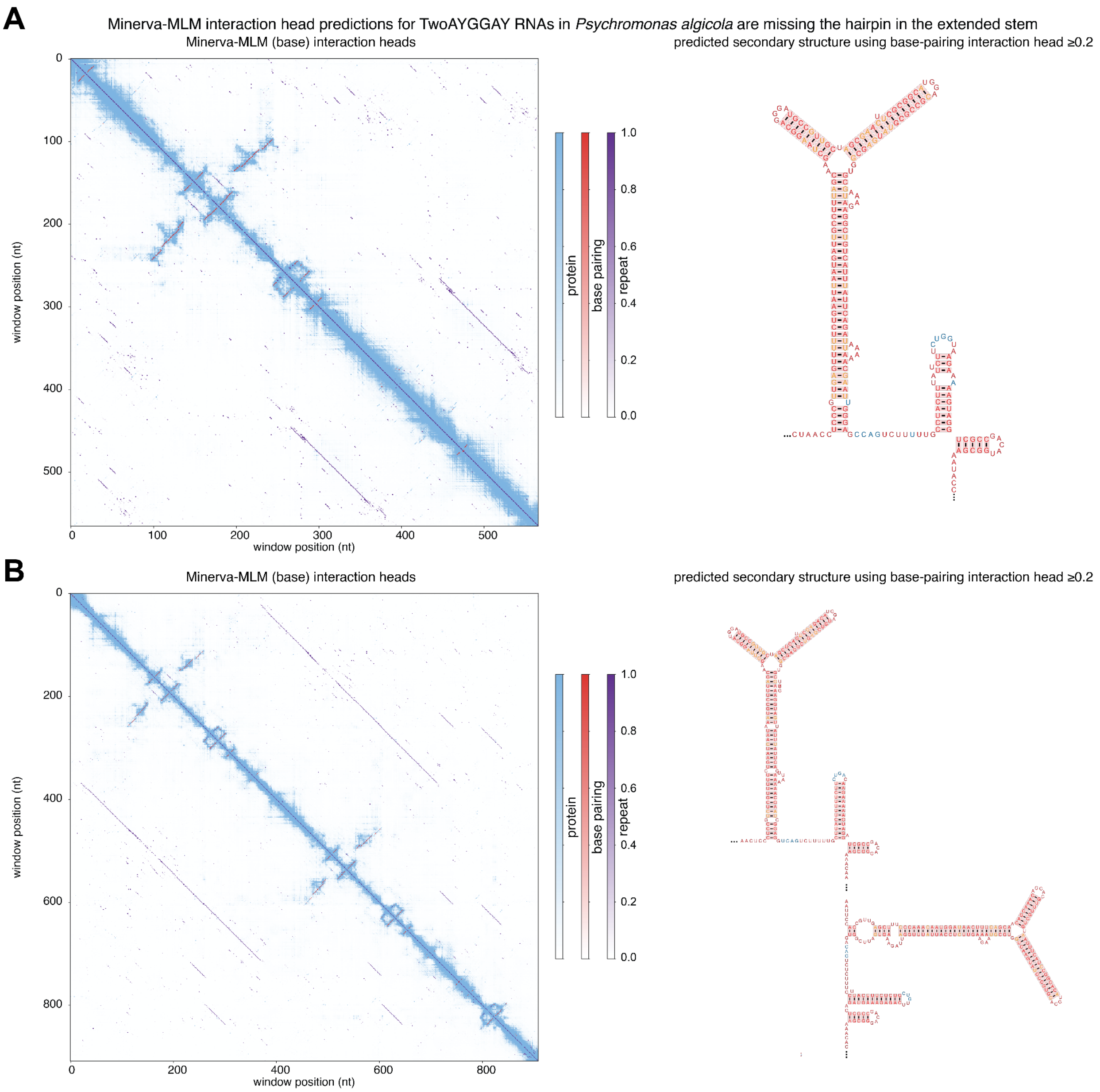


**Figure S26: TwoAYGGAY variants in *Psychromonas algicola* lack a hairpin in the extended stem**

**(A–B)** Minerva-MLM base-model interaction-head predictions for two representative *Psychromonas algicola* TwoAYGGAY RNAs. Predicted secondary structures generated from base-pairing interaction-head scores of at least 0.2 are shown to the right and lack the hairpin within the extended basal stem. Positions predicted to be involved in base-pairing interactions inconsistent with Watson-Crick base-pairing are colored orange. Positions predicted to be involved in a pseudoknot consistent with Watson-Crick base-pairing are colored blue.

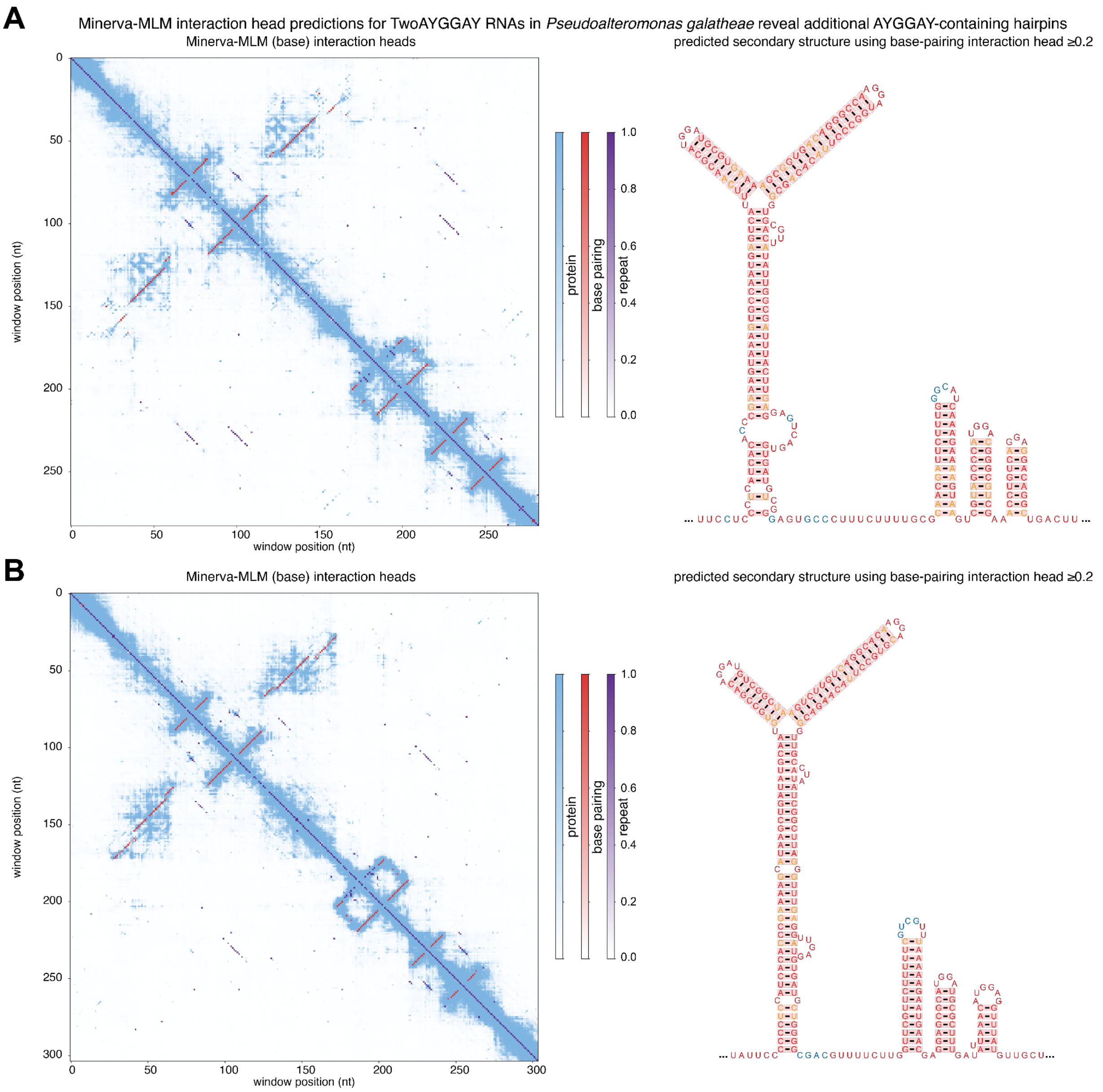


**Figure S27: TwoAYGGAY variants in *Pseudoalteromonas galatheae* contain additional AYGGAY hairpins**

**(A–B)** Minerva-MLM base-model interaction-head predictions for two representative *Pseudoalteromonas galatheae* TwoAYGGAY RNAs. Predicted secondary structures generated from base-pairing interaction-head scores of at least 0.2 are shown to the right and contain additional AYGGAY-motif-bearing hairpins. Positions predicted to be involved in base-pairing interactions inconsistent with Watson-Crick base-pairing are colored orange. Positions predicted to be involved in a pseudoknot consistent with Watson-Crick base-pairing are colored blue.

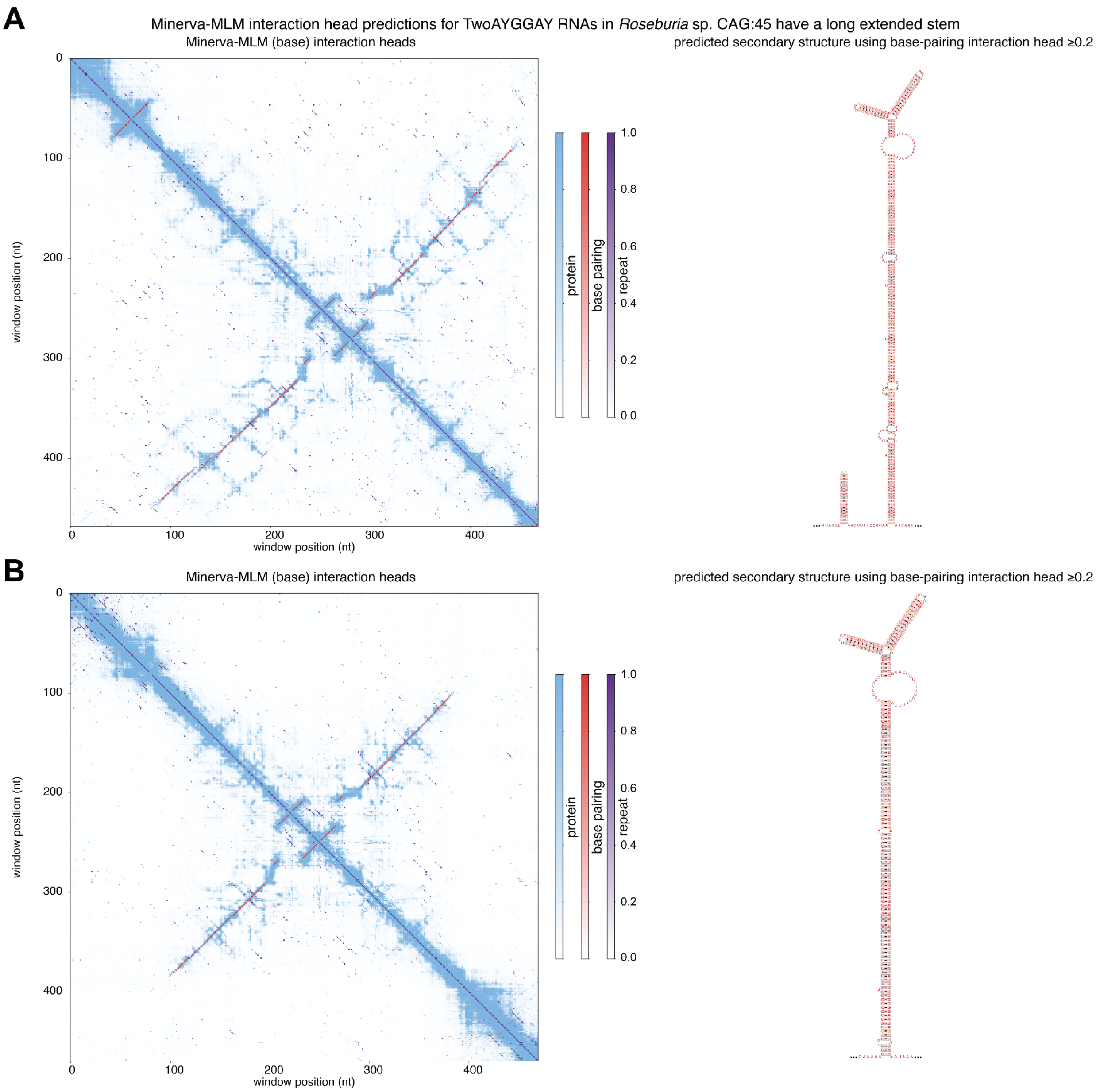


**Figure S28: TwoAYGGAY variants in *Roseburia* sp. CAG:45 contain an elongated basal stem**

**(A–B)** Minerva-MLM base-model interaction-head predictions for two representative *Roseburia* sp. CAG:45 TwoAYGGAY RNAs. Predicted secondary structures generated from base-pairing interaction-head scores of at least 0.2 are shown to the right and contain an elongated basal stem without the Pseudomonadota-associated extension. Positions predicted to be involved in base-pairing interactions inconsistent with Watson-Crick base-pairing are colored orange. Positions predicted to be involved in a pseudoknot consistent with Watson-Crick base-pairing are colored blue.

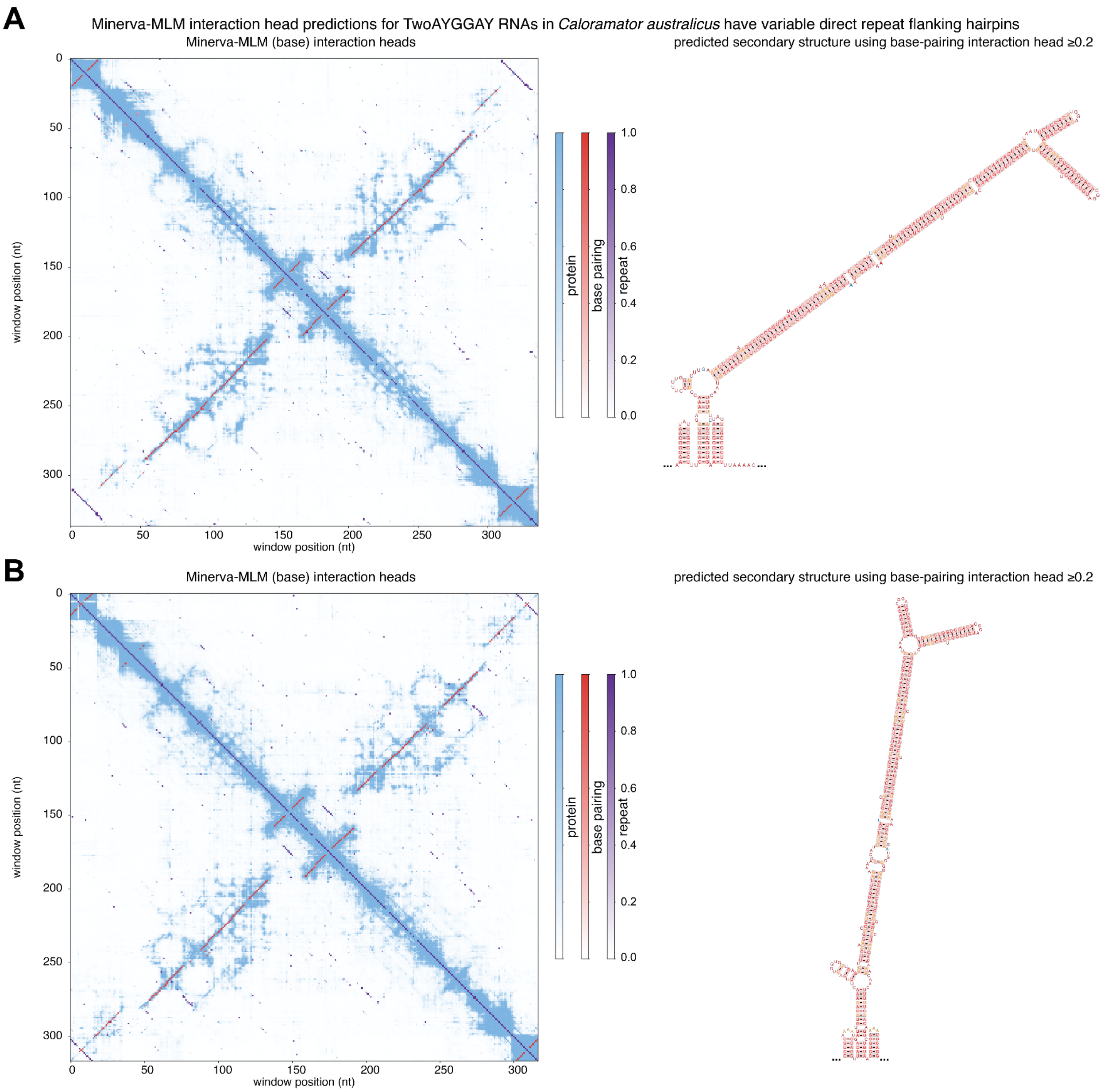


**Figure S29: TwoAYGGAY variants in *Caloramator australicus* contain variable flanking hairpin repeats**

**(A–B)** Minerva-MLM base-model interaction-head predictions for two representative *C. australicus* TwoAYGGAY RNAs. Predicted secondary structures generated from base-pairing interaction-head scores of at least 0.2 are shown to the right and contain variable direct-repeat hairpins flanking the extended basal stem. Positions predicted to be involved in base-pairing interactions inconsistent with Watson-Crick base-pairing are colored orange. Positions predicted to be involved in a pseudoknot consistent with Watson-Crick base-pairing are colored blue.

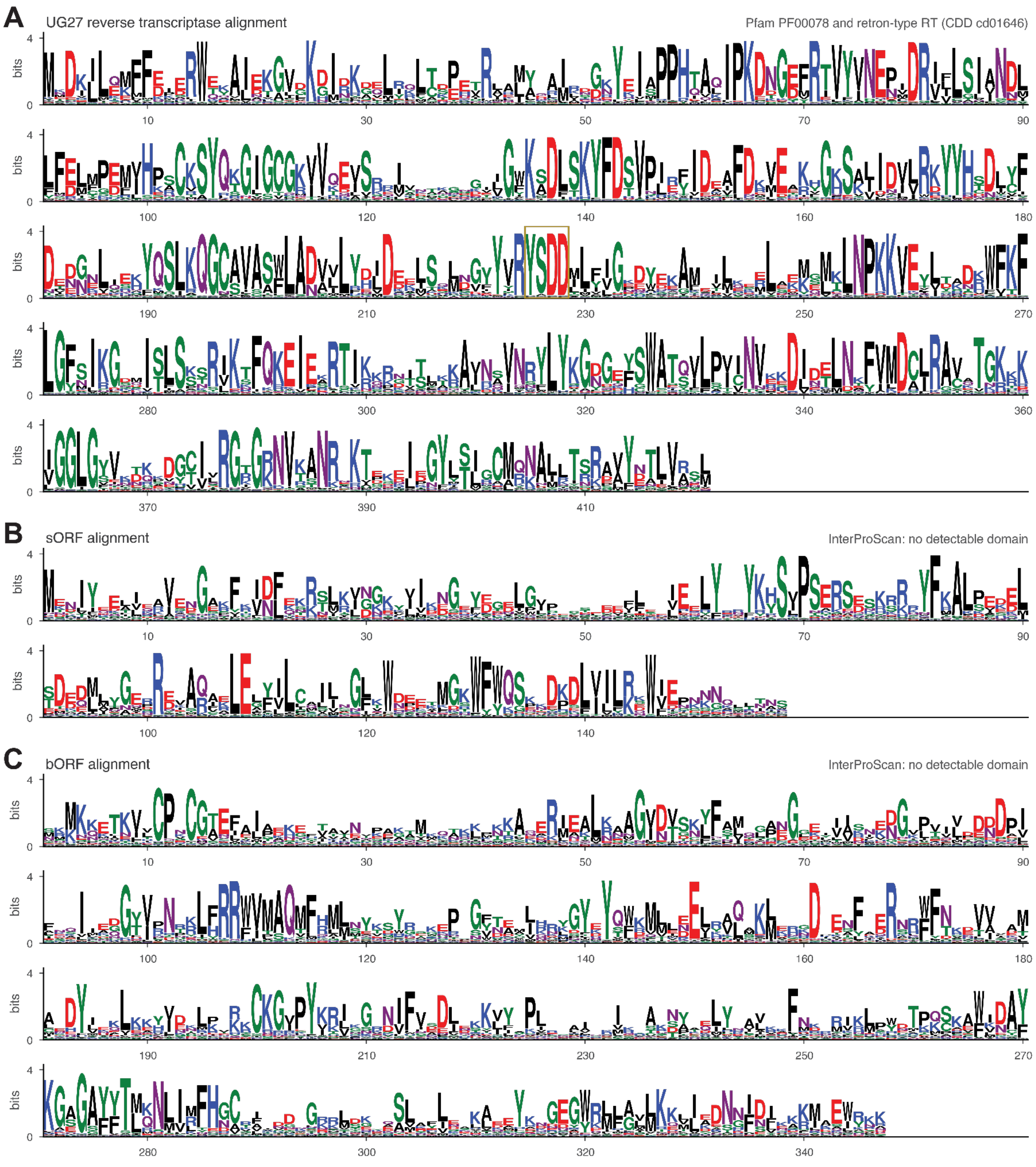


**Figure S30: Sequence conservation of the three UG27 protein components**

**(A)** Sequence logo for the UG27 reverse transcriptase alignment after deduplication. The YSDD RT active-site motif is boxed. The reverse transcriptase contains regions matching Pfam reverse transcriptase family PF00078 and the retron-type reverse transcriptase conserved domain cd01646.

**(B)** Sequence logo for the UG27 small accessory protein (sORF) alignment after deduplication. No conserved domain was detected by InterProScan.
**(C)** Sequence logo for the UG27 large accessory protein (bORF) alignment after deduplication. No conserved domain was detected by InterProScan.

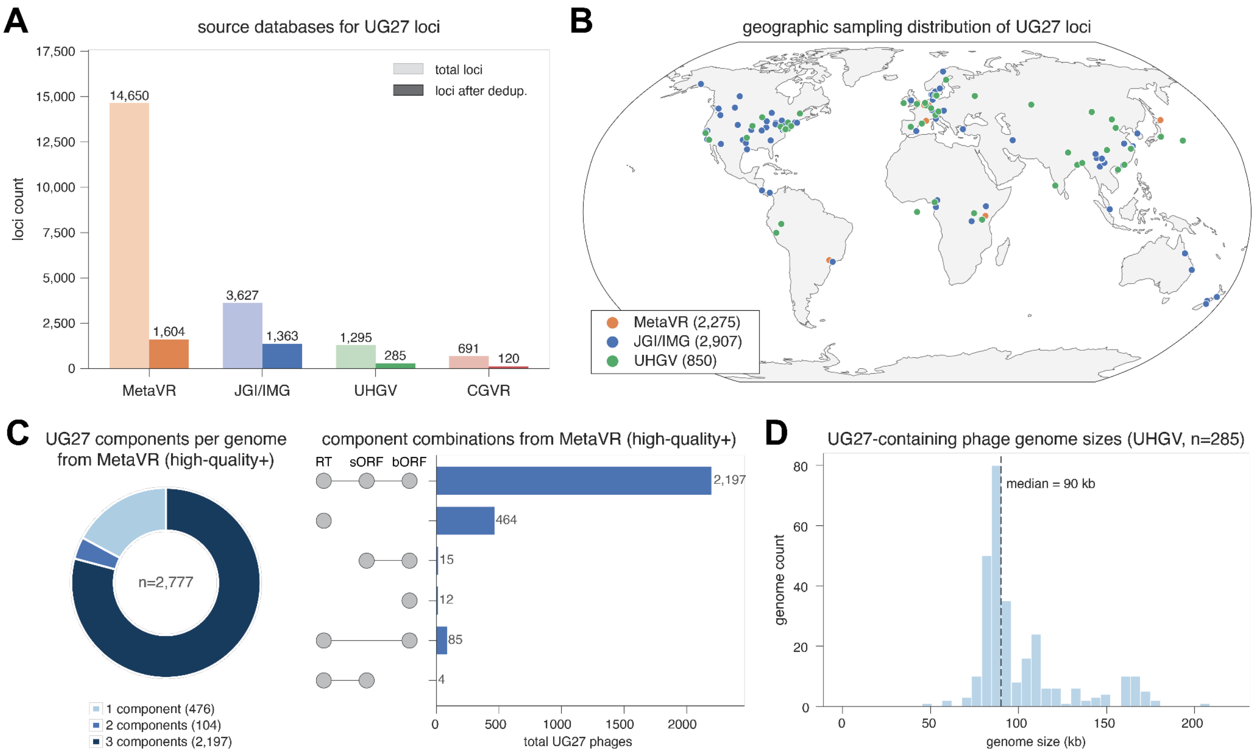


**Figure S31: Assembly and provenance of the UG27 loci dataset**

**(A)** Numbers of UG27 loci recovered from MetaVR, JGI IMG, the Unified Human Gut Virome catalog (UHGV), and the Chinese Gut Viral Reference (CGVR) before and after species-level deduplication.

**(B)** Geographic sampling distribution of UG27 loci from MetaVR, JGI IMG, and UHGV for records with available location metadata.

**(C)** Number of conserved UG27 protein components encoded per high-quality or complete MetaVR phage genome and counts of the observed RT, sORF, and bORF component combinations.

**(D)** Genome-size distribution of UG27-containing phages in UHGV, classified by CheckV as complete or high-quality (≥90% completeness). The median genome size is indicated.

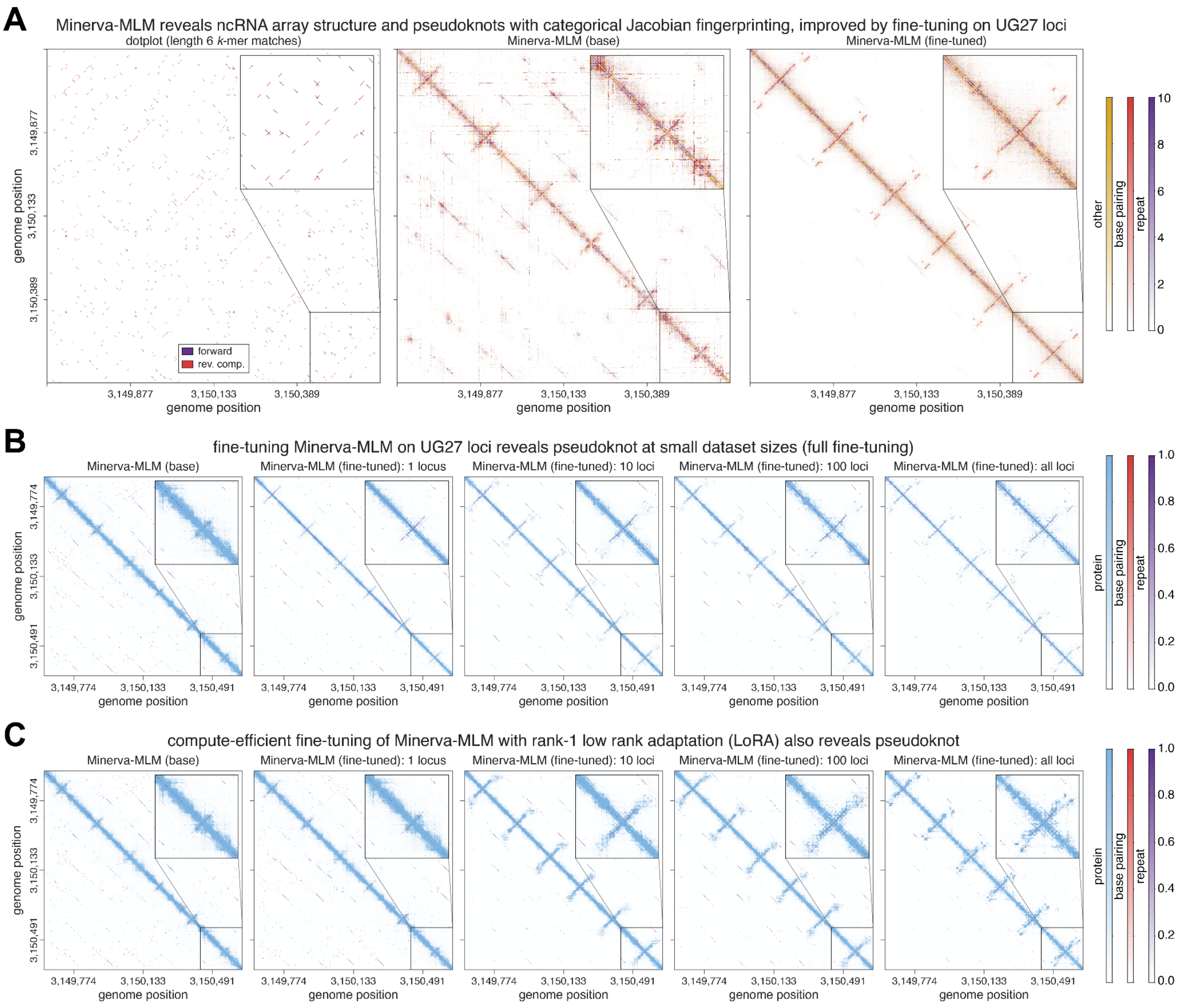


**Figure S32: Fine-tuning Minerva-MLM on UG27 loci resolves the ncRNA pseudoknot, including with rank-1 LoRA**

**(A)** Sequence dotplot showing exact 6-mer matches in the forward and reverse-complement orientations, alongside Minerva-MLM categorical Jacobian fingerprinting maps for the ncRNA array in an intergenic region of the *Bacteroides* sp. 2-1-16 UG27 system. Categorical Jacobian fingerprinting predictions shown are before and after UG27 family-specific fine-tuning. Insets enlarge a single ncRNA unit.

**(B)** Minerva-MLM interaction-head predictions for the same region using the base model and models fine-tuned on 1, 10, 100, or 2,549 UG27 loci (deduplicated at 100% protein identity) using full-parameter fine-tuning.

**(C)** As in **(B)**, using rank-1 low-rank adaptation instead of full-parameter fine-tuning.

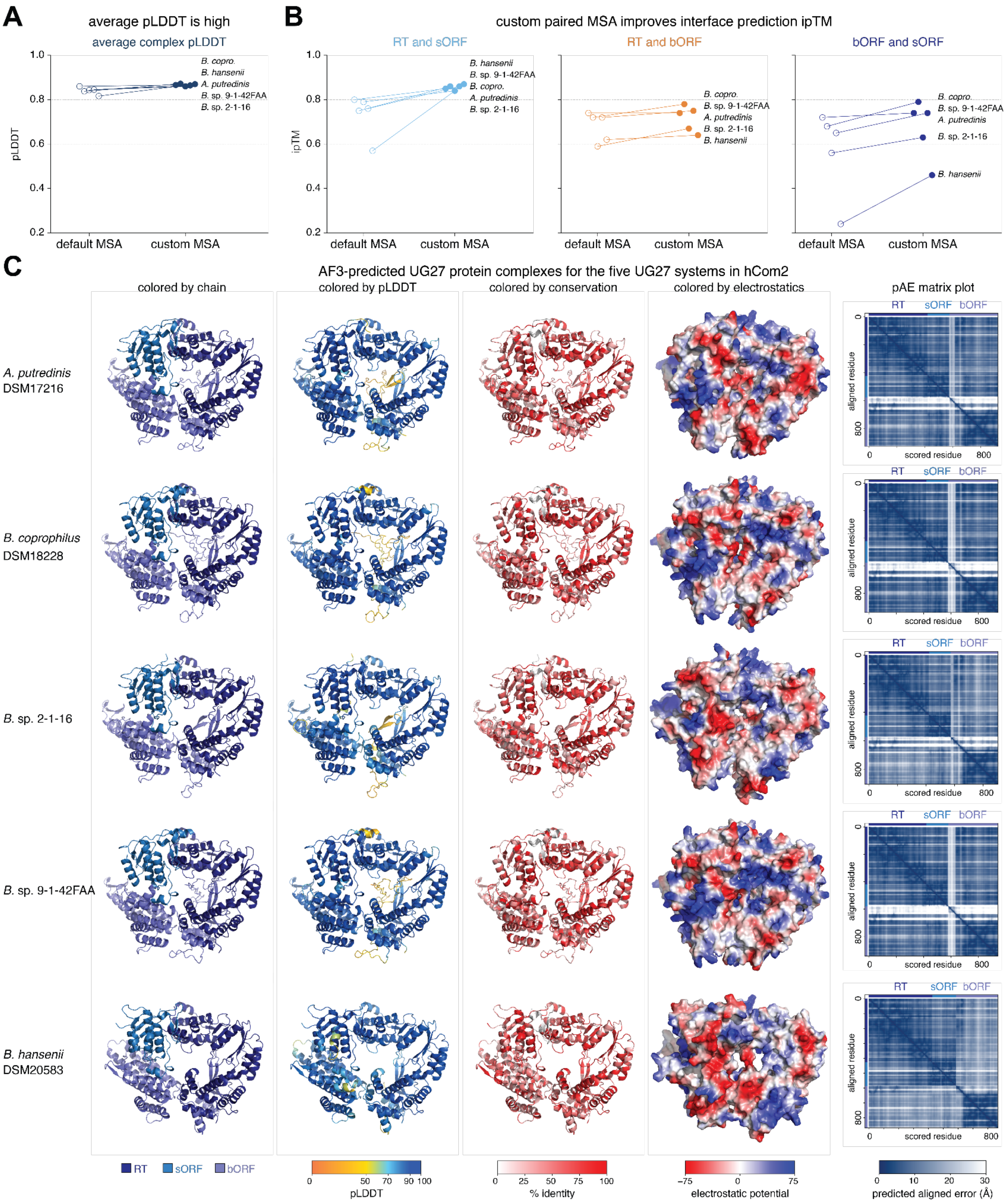


**Figure S33: AlphaFold3 predictions support a three-protein UG27 complex**
**(A)** Mean complex predicted local distance difference test (pLDDT) score for AlphaFold3 predictions of the five hCom2 UG27 protein systems generated using default AF3 MSAs or custom paired alignments incorporating metagenomic loci.

**(B)** Pairwise interface predicted template modeling (ipTM) scores for the RT–sORF, RT–bORF, and bORF–sORF interfaces using default AF3 MSAs or custom paired alignments incorporating metagenomic loci.
**(C)** AlphaFold3-predicted UG27 protein complexes for the five hCom2 systems, colored by protein chain, pLDDT, sequence conservation, or electrostatic potential. Predicted aligned error matrices are shown on the right, with RT, sORF, and bORF boundaries indicated.

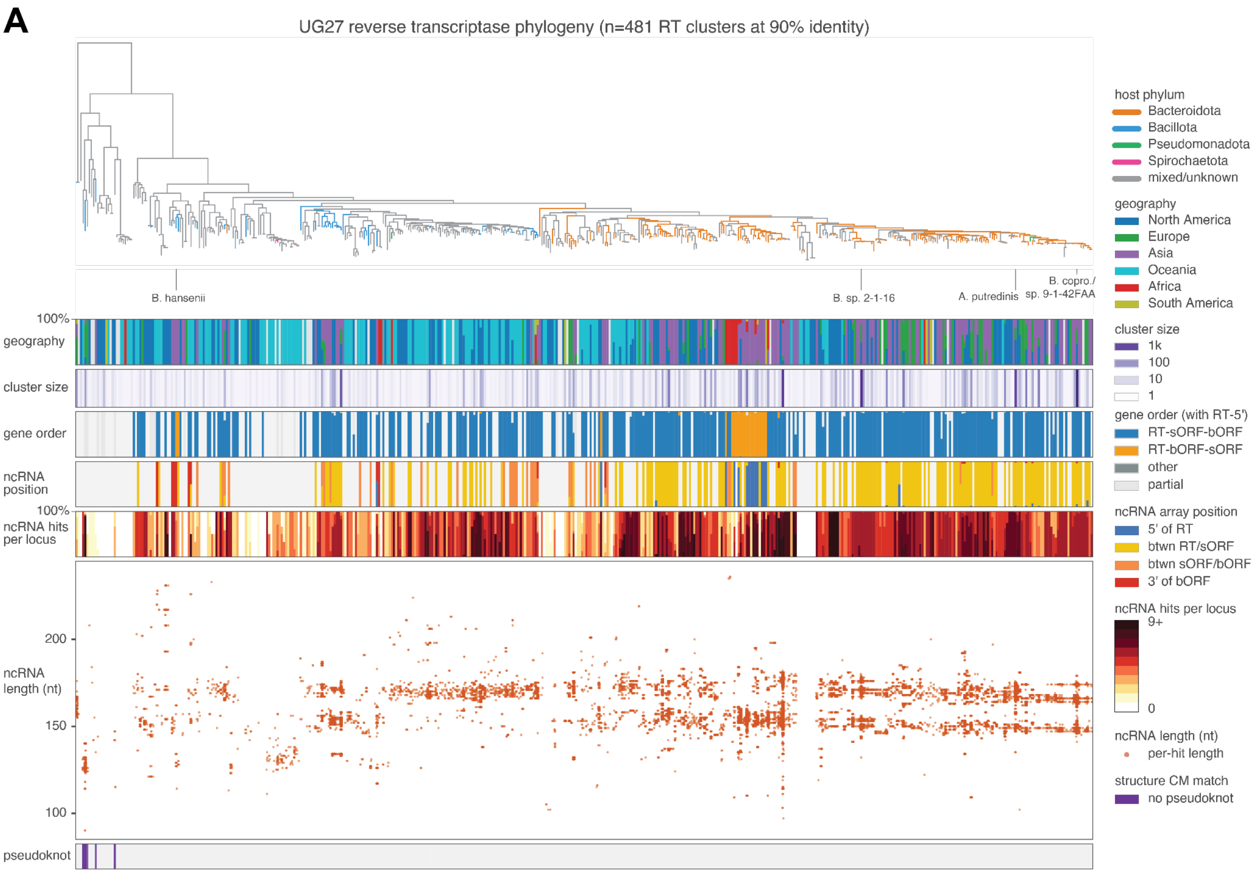


**Figure S34: Phylogenetic and architectural diversity of UG27 systems**
**(A)** Maximum-likelihood phylogenetic tree of 481 UG27 reverse transcriptase clusters defined at 90% amino-acid identity, with branches colored by predicted host phylum. Annotation tracks show (i) the geographic distribution of the data sources from which the constituent loci were drawn; (ii) cluster size, defined as the number of sequences represented by each leaf; (iii) gene-order architecture of the UG27 elements, oriented with the RT 5′-most; (iv) the position of the ncRNA array relative to the protein-coding genes; (v) the distribution among constituent loci of the number of ncRNA hits recovered by the two covariance models; (vi) the lengths of individual ncRNA units within each cluster; and (vii) clusters in which the covariance model lacking the pseudoknot produced a more significant match than the pseudoknot-containing model, consistent with ncRNAs lacking the pseudoknot. The five experimentally tested hCom2 systems are labeled.

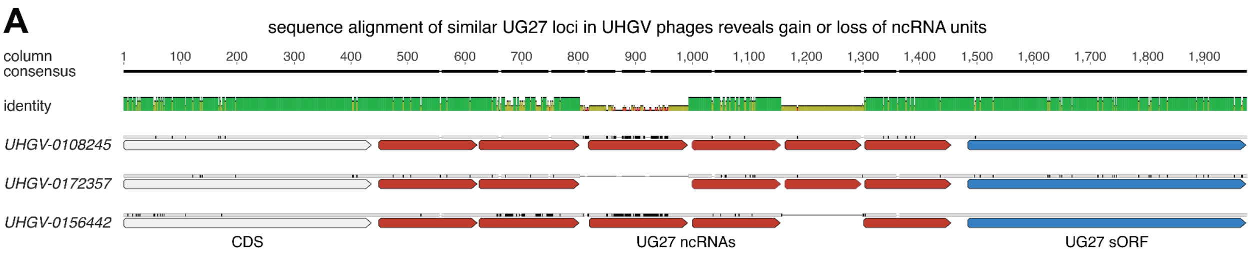


**Figure S35: Closely related UG27 loci differ by the gain or loss of individual ncRNA units** **(A)** Nucleotide alignment of three closely related UG27 loci from UHGV phages (UHGV-0108245, UHGV-0172357, UHGV-0156442), shown with column consensus and per-column identity. Annotated features are an upstream CDS, the array of UG27 ncRNA units, and the UG27 sORF. The loci differ by the presence or absence of individual ncRNA units.

**A**

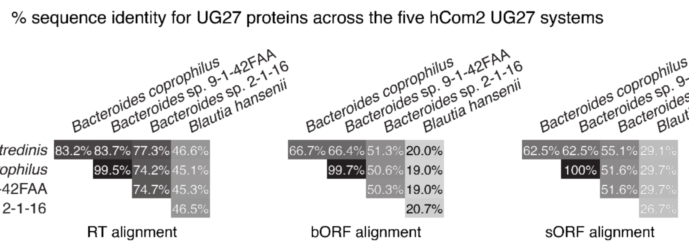


**Figure S36: Protein sequence identity among the five experimentally tested UG27 systems** **(A)** Pairwise amino-acid sequence identity among the RT, bORF, and sORF proteins from the five hCom2 UG27 systems. Percent identities were calculated from the corresponding protein alignments.

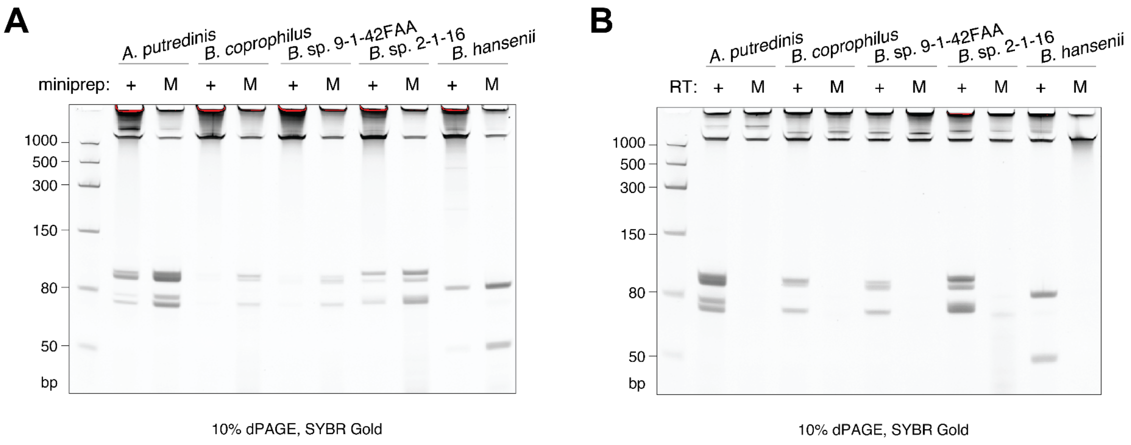


**Figure S37: UG27 systems produce short RT-dependent cDNA products, with capture enhanced by a modified miniprep protocol**

**(A)** Denaturing 10% TBE-urea PAGE of nucleic acids purified from *E. coli* heterologously expressing wild-type versions of each of the five UG27 systems using a standard (+) or modified (M) miniprep protocol. Nucleic acids were visualized with SYBR Gold.

**(B)** Denaturing 10% TBE-urea PAGE of nucleic acids purified from *E. coli* heterologously expressing wild-type (+) or catalytically inactive RT YSAA mutant (M) versions of each of the five UG27 systems. Nucleic acids were visualized with SYBR Gold. This experiment is an independent biological replicate of **Figure 6B**.

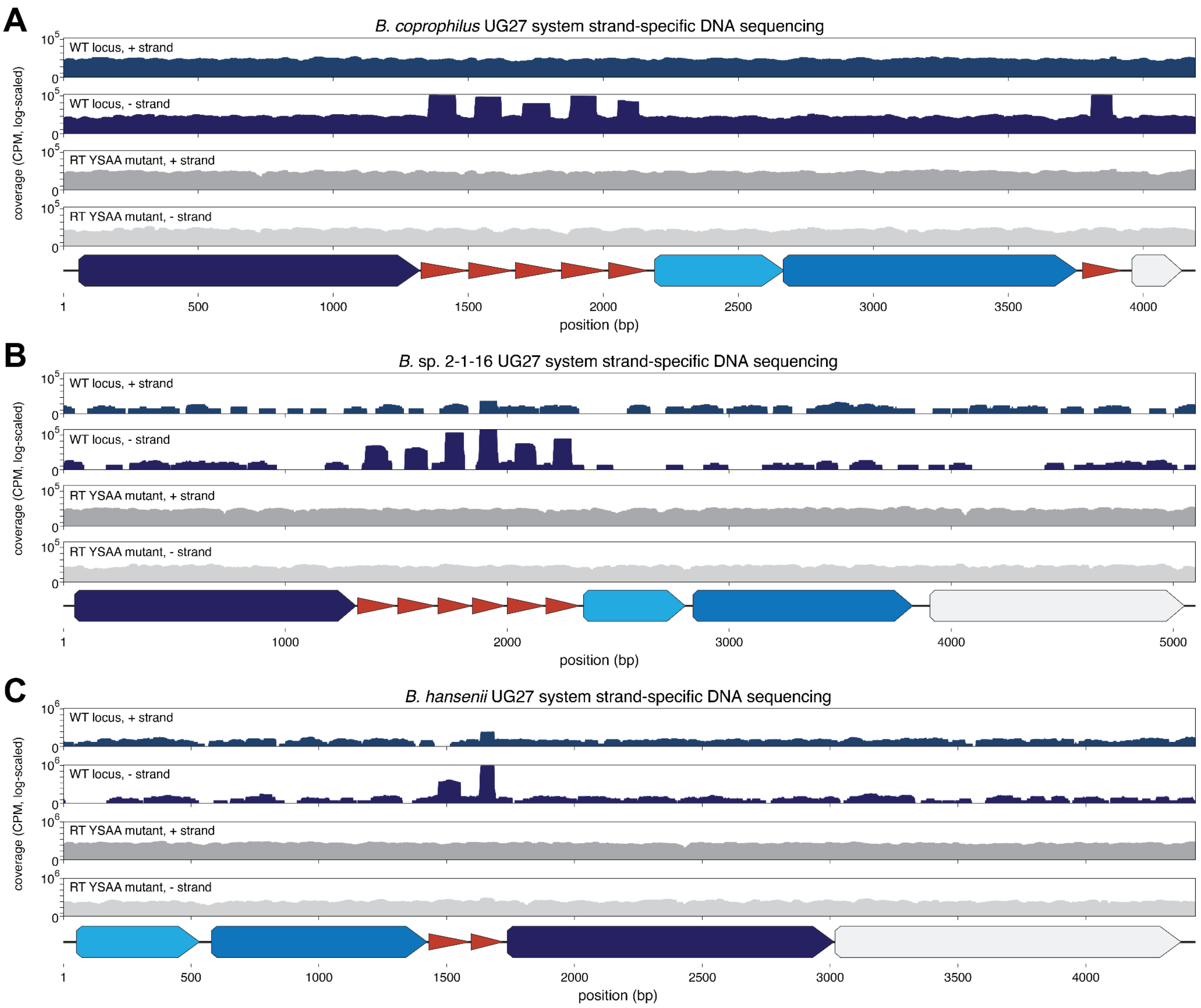


**Figure S38: Strand-specific sequencing maps UG27 cDNA products to central hairpins within the ncRNA arrays**

**(A–C)** Strand-specific DNA sequencing coverage (in counts per million, CPM) across the UG27 loci from **(A)** *Bacteroides coprophilus*, **(B)** *Bacteroides* sp. 2-1-16, and **(C)** *Blautia hansenii*. Coverage from wild-type systems and catalytically inactive RT YSAA mutants is shown separately for the positive and negative strands on a logarithmic scale. UG27 protein-coding genes and ncRNA units are annotated below each locus.

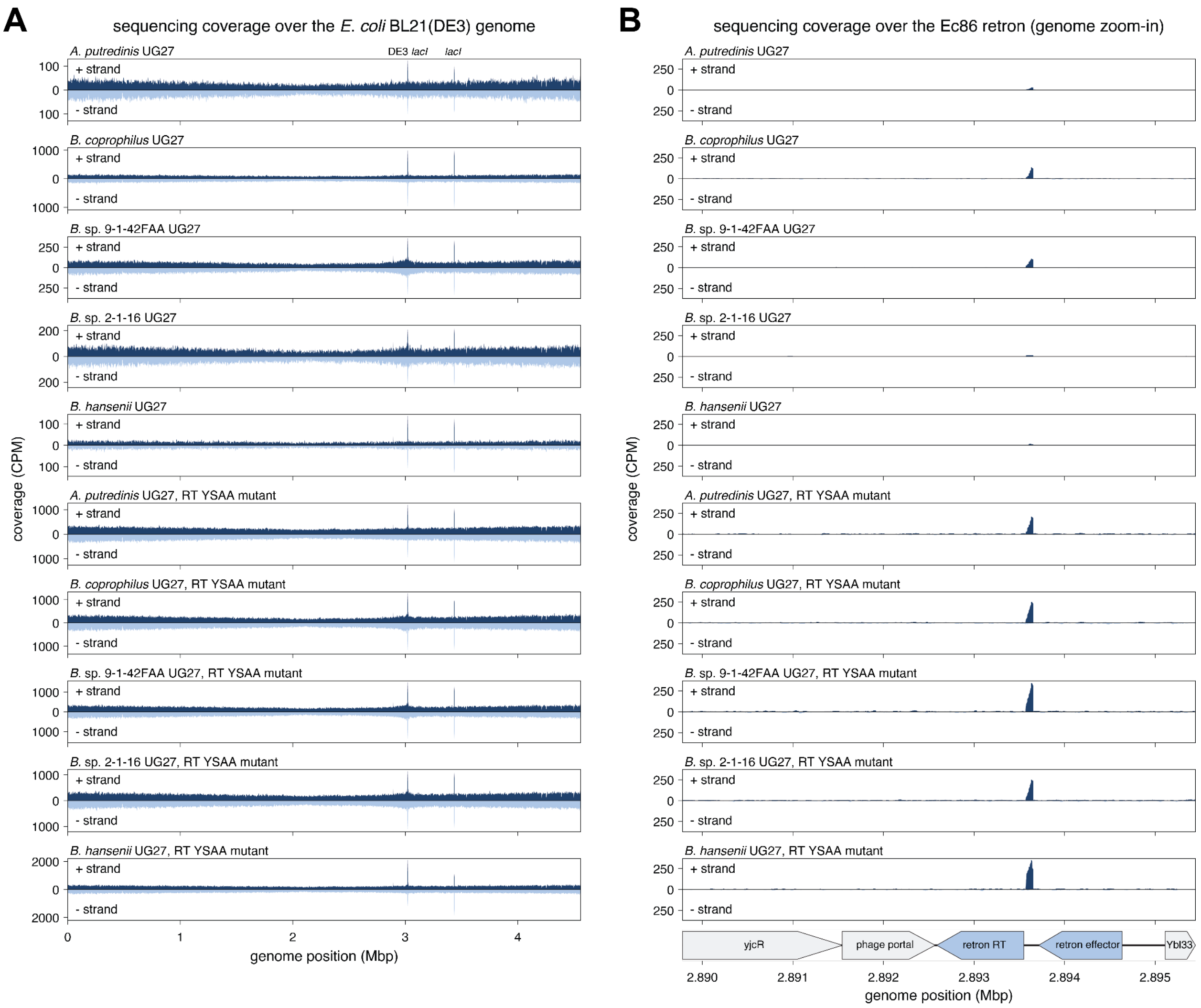


**Figure S39: Genome-wide sequencing coverage and recovery of the endogenous Ec86 retron product**

**(A)** Positive- and negative-strand DNA sequencing coverage (in counts per million, CPM) across the *E. coli* BL21(DE3) genome for cells expressing each wild-type UG27 system or its catalytically inactive RT YSAA mutant. The DE3-associated and native chromosomal *lacI* loci are annotated.

**(B)** Enlarged view of coverage (in counts per million, CPM) across the endogenous Ec86 retron locus for the same samples. The retron reverse transcriptase, retron effector, and neighboring genes are annotated below.

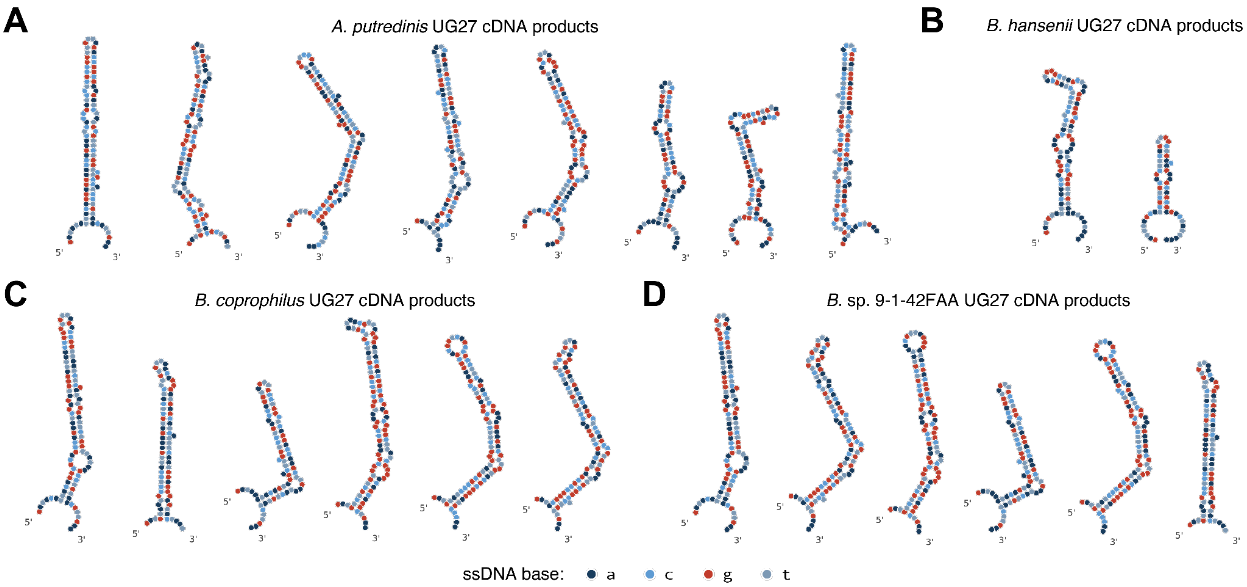


**Figure S40: Predicted secondary structures of UG27 cDNA products**

**(A–D)** Predicted secondary structures of sequenced cDNA products from the **(A)** *Alistipes putredinis*, **(B)** *Blautia hansenii*, **(C)** *Bacteroides coprophilus*, and **(D)** *Bacteroides* sp. 9-1-42FAA UG27 systems. Structures were predicted using ViennaRNA with DNA parameters; nucleotide identity is indicated by color.

**Table S1:** The 150 bacterial genomes scanned in this study (see Supplementary File).

**Table S2:** Model and training hyperparameters for Minerva-MLM and Minerva-MLM-8k (see Supplementary File).

**Table S3:** Primers and constructs used in this study (see Supplementary File).
